# SARS-CoV-2 Nucleocapsid protein is decorated with multiple N- and O-glycans

**DOI:** 10.1101/2020.08.26.269043

**Authors:** Nitin T. Supekar, Asif Shajahan, Anne S. Gleinich, Daniel Rouhani, Christian Heiss, Parastoo Azadi

## Abstract

Severe acute respiratory syndrome coronavirus 2 (SARS-CoV-2), which causes coronavirus disease (COVID-19) started at the end of 2019 in Wuhan, China has spread rapidly and became a pandemic. Since there is no therapy available that is proven as fully protective against COVID-19, a vaccine to protect against deadly COVID-19 is urgently needed. Nucleocapsid protein (N protein), is one of the most abundant proteins in coronaviruses and is a potential target for both vaccine development and point of care diagnostics. The variable mass of N protein (45 to 60 kDa), suggests the presence of post-translational modifications (PTMs), and it is critical to clearly define these PTMs to gain the structural understanding necessary for further vaccine research. There have been several reports suggesting that the N protein is phosphorylated but lacks glycosylation. Our comprehensive glycomics and glycoproteomics experiments confirm that the N protein is highly O-glycosylated and also contains significant levels of N-glycosylation. We were able to confirm the presence of O-glycans on seven sites with substantial glycan occupancy, in addition to less abundant O-glycans on four sites. We also detected N-glycans on two out of five potential N-glycosylation sites. Moreover, we were able to confirm one phosphorylation site. Recent studies have indicated that the N protein can serve as an important diagnostic marker for coronavirus disease and a major immunogen by priming protective immune responses. Thus, detailed structural characterization of the N protein may provide useful insights for understanding the roles of glycosylation on viral pathogenesis and also in vaccine design and development.

## Introduction

In early December 2019 in Wuhan, China, an outbreak of disease caused by severe acute respiratory syndrome coronavirus 2 (SARS-CoV-2), commonly known as 2019 coronavirus disease (COVID-19), spread rapidly and became a worldwide pandemic.^1^ According to the World Health Organization (WHO) report, as of August 25, 2020, there have been 23.7 million confirmed cases of COVID 19 globally, including over 814,000 deaths. Along with other medically advanced countries like Italy, Spain, France and Germany, the United States of America (USA) is one of the most affected countries in the world. To date, the USA alone has over 5.7 million confirmed cases and over 176,000 deaths due to COVID-19.^2^ Therefore, a vaccine to protect against COVID-19 is urgently needed. Multiple preclinical studies have shown positive results, but despite the continuous efforts by the scientific community towards a vaccine or a specific treatment, no vaccine is currently available in USA or Europe that is proven effective and protective against COVID-19.^3–4^

The coronavirus virion consists of a nucleocapsid that is surrounded by a lipid envelope in which the membrane glycoprotein (M) and small transmembrane protein (E) are embedded. Protrusions composed of trimeric glycoproteins (Spike protein, S) are anchored in the lipid envelope and extend radially (Figure 1). These Spike proteins give the spherical viral particles an appearance reminiscent of the solar corona, hence the name for this group of viruses. The S protein is one of the most studied proteins in SARS-CoV-2 and has been under evaluation as a vaccine target.^4^ The S protein is highly glycosylated and binds to angiotensin converting enzyme 2 (hACE2) for entry into the cells. A number of studies have highlighted the importance of spike protein glycans, which play a critical role in the mechanism of virus attachment to the hACE2 receptor. CD8^+^ T cells are known to detect and neutralize virus-infected cells by recognizing viral protein-derived peptide epitopes presented by major histocompatibility complex class I (MHC-I) proteins on the cell surface.^5^ Recently, however, a study involving SARS-CoV-2 patients identified the majority of immunodominant CD8^+^ T cell epitopes from virus proteins other than the S protein, and leading to the conclusion that more protein targets need to be included for new and more effective vaccine design.^6^

**Figure 1.**
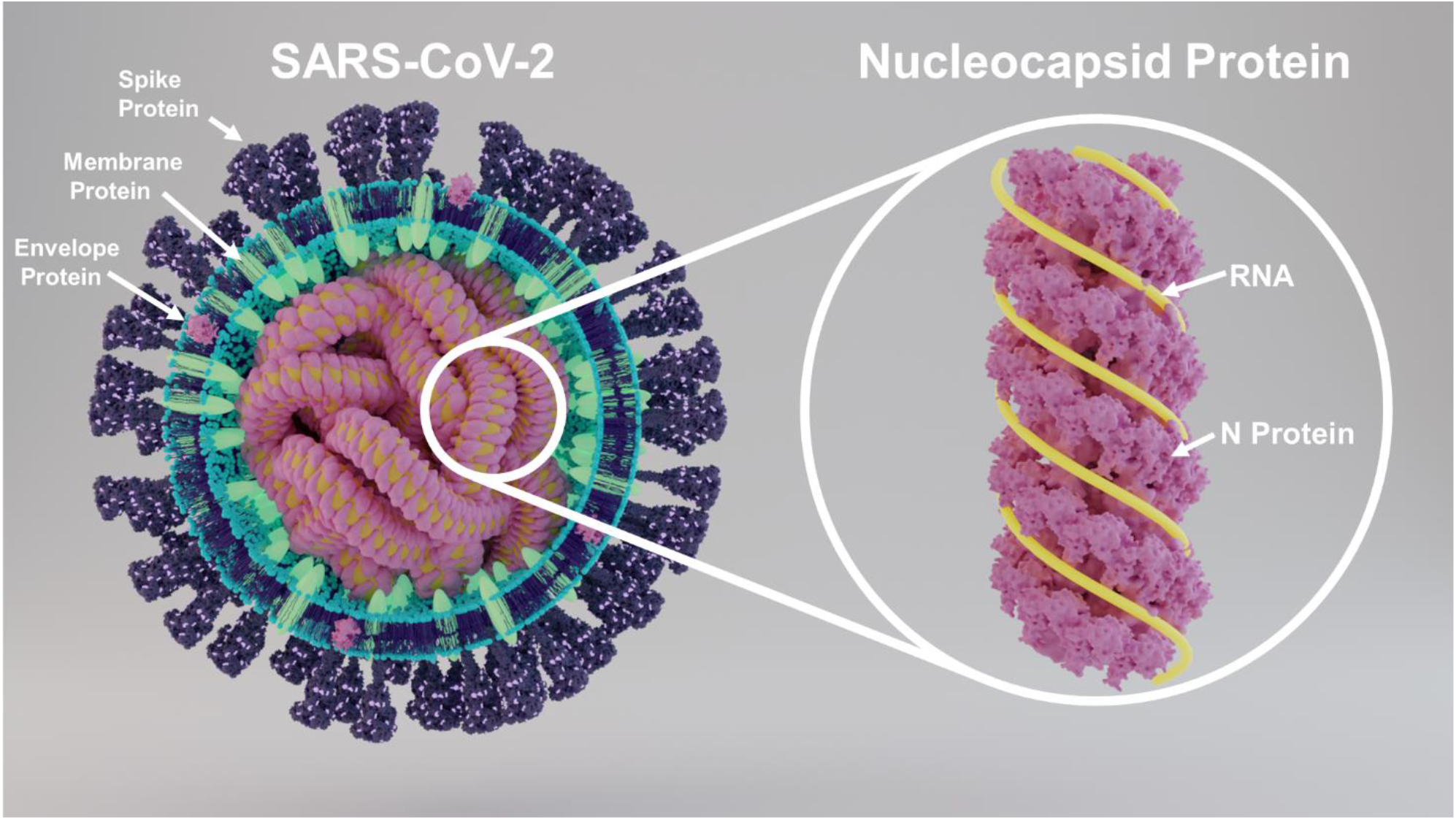
Structure of SARS-CoV-2 showing key proteins and structure of nucleocapsid protein

The nucleocapsid (N) protein is one of the most abundant proteins in the coronavirus. Numerous copies of the N protein interact with RNA molecules during the viral life cycle. The N protein, in conjunction with the membrane (M) protein, also contributes towards genome condensation and packaging. The C-terminal interactions between the N and M proteins result in specific genome encapsidation during the budding process of the viral particle. The N protein has a molecular mass ranging from 45 to 60 kDa - possibly due to PTMs - and is highly basic in nature.^7^ After sequence comparison of various strains, it is believed that the majority of N proteins are composed of three highly conserved domains known as Domains 1, 2 and 3.^7–8^ Domains 1 and 2 are rich in arginine and lysine, a characteristic feature of several viral RNA-binding proteins. The carboxy-terminal Domain 3 is short and contains a net negative charge due to multiple acidic residues.^7^

Mapping studies of SARS-CoV-2 and other coronaviruses, including the closely related SARS coronavirus (retrospectively named SARS-CoV-1), which was the cause of an epidemic in 2003, have linked the RNA-binding function to a fragment of 55 amino acids located at the N-terminal half that resides in Domain 1, and the dimerization function to its C-terminal half from Domain 3 of the N protein. The linker region (LKR) is located in the middle of the protein sequence, connects the N- and C-terminal domains of the protein and includes a Ser/Arg-rich (S/R-rich) region.^9^ Studies have shown that the S/R-rich LKR contains multiple putative sites of phosphorylation that may play a role in regulating N protein function and N-M protein interactions.^10–13^ The N protein interacts with viral genome RNA to form long, flexible, helical ribonucleoprotein (RNP) complexes. One of the structural models of coronaviruses (CoV) proposes that numerous copies of the N protein are present in the form of a helical nucleocapsid as well as in the internal spherical or icosahedral core.^14^ The internal core of CoV is composed of N protein, RNA and the C-terminal dimerization domain (CTD) of M protein.^9^ The M protein is the main core shell component, and the CTD of M protein binds directly to the N protein through an ionic interaction resulting in specific genome encapsidation in the budding viral particle. This way, the N protein plays an important structural role in the CoV virion through a series of interactions with the genomic RNA, M protein and other N proteins. Along with the structural roles, the N protein is also associated with other processes such as mRNA transcription, replication, and host cell modulation during viral infection. Interestingly, nucleocapsid proteins of MERS-CoV and SARS-CoV-1 were found to antagonize interferons (INFs).^15–17^ Early studies investigating the SARS-CoV-2 N protein show similar results in the suppression of interferon induction.^18–19^ INFs are signaling molecules of the innate immune system of the host and have been identified as a crucial component in the early phase of viral disease. In detail, type I and type III interferons serve as the first important step of antiviral defense mechanisms that induce downstream antiviral processes.^20^ Studies found that the antagonistic role of the N protein leads to a modulation of the innate immune response of the host, weakening primary antiviral responses. An initial viral evasion from innate immunity affects the course of the disease tremendously.^3, 21^ The N protein can also serve as an important diagnostic marker for coronavirus disease and a major immunogen by priming protective immune responses. The recent mass spectrometry (MS) -based studies of SARS-CoV-2 proteins from gargle solution samples and nasal swab samples of COVID-19 patients detected the presence of N protein peptides, making them a potential diagnostic biomarker for COVID-19.^22–23^ The presence of N protein peptides in the gargle and nasopharyngeal swabs could be used to develop a point of care high throughput test for fast detection of SARS-CoV-2. The authors demonstrated in these studies that SARS-CoV-2 peptidome detection through tandem mass spectrometry can be used as alternative methodologies to PCR and immunodiagnostics. The clinical study on gargle solution showed that peptide ^41^RPQGLPNNTASWFTALTQHGK^61^ from the N protein which comprises an N-glycosylation site can only be detected on COVID-19 positive cases after the removal of N-glycosylations by PNGase F treatment.^22^ Whereas in the latter study, the authors argued that the N protein peptides ADETQALPQR and GFYAQGSR can be detected with intense signal within short retention time on COVID-19 clinical samples.^23^ These recent findings on COVID-19 clinical samples show the presence of N protein by the virus on the host and the potential as a diagnostic biomarker at point of care.

A study of SARS-CoV-2 patients noted that 26 of 29 specific T cell epitopes came from proteins other than the S protein.^6^ Importantly, out of these 29 immunodominant epitopes, 6 derived from the N protein. This makes the N protein a potential target for a new vaccine design which would comprise more than one antigen to elicit a CD8^+^ T cell response against SARS-CoV-2.^6^

Early studies, including recent review articles about the N protein, suggested that the N protein is a phosphoprotein and does not bear glycans on its backbone.^24^ Glycan-detecting software and amino acid sequence analysis have lately indicated the considerable likelihood of the presence of N- and O-glycans. To our knowledge, however, glycan profiling and site-specific information of N- and O-glycans in the N protein have not been described. To discover the PTMs of N protein, we explored several high-resolution MS-based approaches, including glycomics, glycoproteomics, and phosphoproteomics analyses. Our study has revealed new information about the PTMs of N protein: We demonstrate that the SARS-CoV-2 N protein is not only a phosphoprotein but also carries significant amounts of N- and O-glycosylation. MS experiments to determine occupancy and site-specific microheterogeneity, sialic acid linkages, fucosylation and investigation of bisecting GlcNAc of N-linked glycans were also conducted. In our study, we detected mostly high-mannose and complex type glycans, as well as minor percentages of hybrid N-glycans through the glycomics approach. These findings were confirmed in our glycoproteomic investigation: We confirmed presence of N-glycosylation on two of five consensus sequons, which were modified with mostly either high-mannose or complex type N-glycans. Along with N-linked glycosylation, we also observed O-linked glycosylation on the N protein via glycomics and glycoproteomics. Proteomic analysis for O-glycosylation and phosphorylation resulted in the detection of 8 glycopeptides with 11 potential O-glycan sites and one phosphate modification site.

## Results

### Glycomics analysis of N- and O-glycans released from nucleocapsid protein

We accomplished detailed structure characterization of the glycans of N protein that was recombinantly expressed in HEK293 cells using full MS and MS^n^ experiments of the released and permethylated glycans. The N protein was initially purified using SDS-PAGE for both N- and O-glycomics experiments (Fig. S1). The N-glycans were released from the cut gel band by treatment with PNGase F enzyme. The released N-glycans were extracted from the gel pieces and were purified by C18 SPE. The gel pieces containing the remaining O-glycoprotein were subjected to enzymatic digestion via trypsin and were extracted from gel bands with acetonitrile-water mixture. The obtained O-glycopeptides were used for the release of O-glycans by β-elimination. Both N- and O-glycans were permethylated and analyzed by MALDI-MS and ES-IMS^n^ for the detailed structural characterization.

We observed mostly high-mannose type N-glycans (~73%), along with smaller amounts of complex (~22%) and hybrid type (~5%) glycans. Sialylated and non-sialylated glycan structures with multiple fucosylation were observed in lesser abundance (~7%) (Fig. 3A and Table S1). We confirmed the structure of the permethylated N-glycans by ESI-MS^n^. Interestingly, we also observed terminal GalNAc structures and lack of branching other than biantennary, and these are discussed in the corresponding sections below.

The O-glycans detected on the N protein were Core 1, Core 2, Core 3, and Core 4 types; however, Core 1 and Core 2 types were predominant, comprising 96% of total O-glycans (Figure 3B and Table S2). As shown in, Core 3 and Core 4 structures, which are formed by the activity of the Core 3 GlcNAcT and Core 4 GlcNAcT enzymes,^25^ were relatively low (Figure 3B and Table S2). The structural features such as branching, sialic acid linkage, position of fucose on O-glycans were also confirmed by ESI-MS^n^.

### Evaluation of the type of branching and presence of bisecting GlcNAc on the N protein N-glycans

We evaluated the type of branching on the released N-glycans from N protein by ESI-MSn analysis. Interestingly, we did not observe any evidence for the presence of common tri- or tetra-antennary and bisecting GlcNAc structures. Instead, we observed uncommon N-glycans with terminal GalNAc and sialic acid linked GalNAc structures. Such GalNAc-GlcNAc-R (LacdiNAc, LDN) glycan structures are known to play critical role in many biological processes and are considered key determinants that can be recognized by the adaptive and innate immune systems via multiple carbohydrate-binding proteins.^26^’^31^ The structural confirmation of LDN was demonstrated by the MSn fragmentation of a representative N-glycan structure in Figure 4A. The dissociation of the N-glycan with mass 2285 produced a doubly charged fragment with m/z 922. Subsequent fragmentation of m/z 922 generated a fragment with m/z 1317, supporting the presence of a biantennary structure with GalNAc termini. Interestingly, any fragments with m/z 1306, which would support the presence of both triantennary and bisecting GlcNAc structures, are completely absent in the spectra. Similar observations were made on other branched N-glycans, indicating that only biantennary N-glycan structures are present on the N protein (Figure 4A).

### Determination of core and antennal fucosylation on N-glycans released from N protein

We evaluated the presence of core and antennal fucosylation on N-glycan structures and observed that core fucosylations were predominant (Table S1). However, we observed mono-fucosylated structures that presented both antennal and core fucosylation. A representative example is the mono-fucosylated N-glycan with m/z 2040. Upon performing MS2 fragmentation of this glycan, we observed the presence of both core and antennal fucosylation as a mixture of isoforms. The m/z 474 fragment and 451 Da neutral loss to form m/z 1588 supports the presence of core fucosylation. On the other hand, presence of fragments with m/z 442 and m/z 701, and 678 Da neutral loss to generate fragment m/z 1361 shows evidence for the presence of antennal fucosylation (Figure 4B).

### Determination of sialic acid linkage on N- and O-glycans released from N protein

Our N-glycan analysis showed that only biantennary structures are present on N protein and gave evidence for the linkage of terminal sialic acids to both galactose and GalNAc on both arms of glycans. We have determined the linkage of sialic acids attached to both galactose and GalNAc by fragmenting terminal trisaccharides on each arm of the N-glycans. We found that the sialic acids could be linked to both O-3 and O-6 of GalNAc (fragments with m/z 150 for 2,6 and m/z 164 for 2,3 linkages) but were exclusively linked to O-3 of galactose (fragments with m/z 109 and 137) (Figure 4C).

We observed only 2,3 linked sialic acid on the O-glycans released from N protein (Figure 4D).

### Characterization of N-glycosylation on nucleocapsid protein by glycopeptide analysis

Since our glycomics analysis confirmed the presence of N-glycans on the N protein, we were interested in determining the site-specific glycan distribution. For this purpose, we used trypsin and elastase proteases both separately and sequentially to produce three glycopeptide samples. The LC-MS/MS data of these protease digests were analyzed by using search algorithms in the Byonic software, employing all possible mammalian N-glycans, O-glycans, and phosphorylation as possible post-translational modifications.

There are five potential N-glycosylation sites on the N protein, and we identified N-glycosylation on two of the sites. Both confirmed N-glycosylation sites have a very distinct N-glycosylation profile. The N-glycosylation site N47 had a glycan occupancy of about 53% (Figure 7A Table S3). We observed only complex type glycan structures at site N47, which is part of glycopeptide ^41^RPQGLPNNTASWF^53^. Based on the common biosynthetic pathway and glycan neutral loss pattern in MS/MS experiments, the glycoform containing NeuAc1GalNAc1Gal1GlcNAc2Man3GlcNAc2Fuc1 at m/z 3590.51 was found to be most abundant with 6.56% glycan occupancy relative to other glycoforms at site N47. Moreover, all the glycans detected at this site featured core fucosylation. We also observed multiple doubly fucosylated glycans and a minor triply fucosylated structure (0.45%) (Tables S1 and S3). The Byonic N-glycopeptide search followed by manual validation for accurate precursor mass (5 ppm) and precise glycan neutral loss in MS/MS experiments (<20 ppm) resulted in detection of N-glycan on peptide ^267^AYNVTQAFGR^276^ at site N269. The glycan occupancy at site N269 was found to be higher (94%) than that of site N47 (53%). At site N269, we detected mostly high-mannose type glycosylation comprising ~85% of total glycan. The relative percentage of complex and hybrid type N-glycans was lower (~15%) (Tables S1 and S3). The tryptic peptide ^69^GQGVPINTNSSPDDQIGYYR^88^ was detected only in non-glycosylated form (Figure S25), indicating that N77 is not occupied. The N-glycosylation sites N192 and N196 occur in the linker region, which is rich in serine and arginine (SR-rich). Investigation of the N-glycosylation at these sites was extremely difficult because the two N-glycan sequons are located next to each other, separated by only one arginine residue (Figure 2A). For site N192, the trypsin digest possibly generated the tripeptide ^192^NSS^195^ (Figure 2A) whose small size would make it difficult to fragment in mass spectrometry and thus manual detection as well as detection through software was challenging. For site N196, *in silico* trypsin digestion generated the peptide ^196^NSTPGSSR^203^, which also contains four potential O-glycan sites (Figure 2A). The software searches for the possible N-glycan combined with four O-glycans were not successful due to software limitations. Similarly, *in silico* analysis of the elastase digest of N protein produced the longer peptide ^183^SSRSSSRSRNSSRNSTPGSSRGTSPA^208^, comprising two potential N-glycosylation sites and fourteen potential O-glycosylation sites, presenting a formidable challenge to software-based detection (Figure 2A). All these factors made the characterization of N- and O-glycosylation at this SR-rich linker region extremely difficult. To overcome this obstacle, we set out to locate the O-glycans on this peptide first, using the de-N-glycosylated N protein. We reasoned that searching for O-glycosylation in the SR-rich region would later be helpful in finding N-glycans since confirmed O-glycan information from this de-N-glycosylation experiment would enable manual/software assisted search for N-glycosylation in non-de-N-glycosylation experiments. To investigate O-glycosylation in the SR-rich region, we first removed the N-glycans from the N protein by PNGase F treatment and subsequently digested the protein by trypsin/elastase. The resulting O-glycopeptides were subjected to LC-MS/MS analysis, followed by manual and software-assisted searches for O-glycans. This eliminated interference of N-glycans while searching for O--glycans and reduced the number of possible combinations for manual/software-based glycan search. We detected the O-glycosylated ^204^GTSPAR^209^ glycopeptide from the linker region and observed several O-glycans on the peptide (see detection of O-glycosylation section below). However, even with this stepwise approach, we could not identify any N-glycans at sites N192 and N196. We also evaluated for the presence of phosphorylation but could not identify any in the linker region.

**Figure 2.**
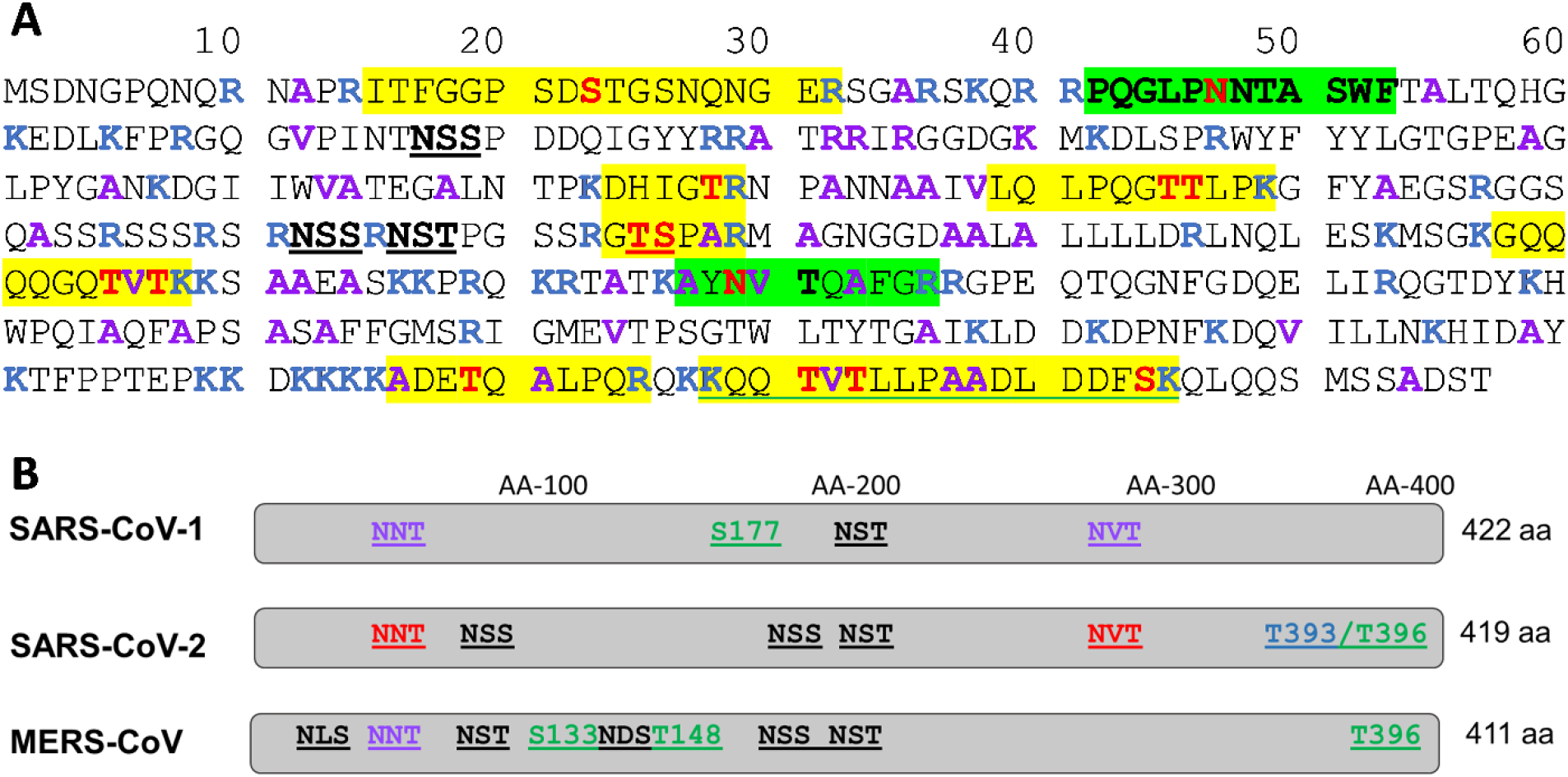
**A.** SARS-CoV-2 N protein sequence, protease cleavage sites and detected glycosylation sites. Legend: Amino acids in red color bold font - detected glycosylation sites; Amino acids in blue color bold font - Trypsin cleavage sites; Amino acids in purple color bold font - elastase cleavage sites; Peptide sequence in green highlight - detected N-glycopeptides; Peptide sequence in yellow highlight - detected O-glycopeptides; Peptide sequence in yellow highlight & green underline - detected O-glycopeptide with phosphate modification; peptide sequence in black color underline – non-glycosylated sites. **B.** Schematic amino acid sequence alignment of the nucleocapsid protein from coronaviruses SARS-CoV-1, SARS-CoV-2 and MERS-CoV (UniProt identifiers SARS-CoV-1 [P59595], SARS-CoV-2 [P0DTC9], MERS-CoV [K9N4V7]). The number of N-linked glycosylation motifs (-NXS/T-with X≠P), accounts for 3 potential sites in SARS-CoV-1, 5 potential N-linked glycosylation sites in SARS-CoV-2, and 6 potential N-glycan sites in MERS-CoV. Highlighted in red are the N-linked glycosylation sites in SARS-CoV-2 that we found occupied in our study. Highlighted in purple are the conserved N-glycan motifs in the depicted coronaviruses; so far, no studies illuminated the glycosylation of the N proteins in SARS-CoV-1 and MERS-CoV. Highlighted in green are the predicted phosphorylation sites Highlighted in green are the predicted phosphorylation sites Highlighted in blue are the detected phosphorylation site in our study

### ^18^O labelling experiment for N-glycosylation site confirmation

To further investigate the possibility of glycosylation at site N192 and N196 and to confirm the glycosylation pattern of the N protein, we performed an ^18^O labelling experiment where N-glycans are removed from glycoproteins PNGase F in the presence of H2^18^O, followed by enzymatic digestion with trypsin/elastase. In the process of de-glycosylation, the N-glycan bearing Asn is converted to ^18^O-Asp, which can be detected by mass spectrometry. In this experiment, we observed conversion of N47 and N269 to ^18^O labelled Asp, confirming the presence of N-glycans in these positions. LC-MS/MS analysis of the trypsin & elastase digests detected the peptides, ^41^RPQGLPD*NTA^50^ and ^268^YD*VTQAFGR^276^ with ^18^O-Asp (D*) at positions 47 and at 269, respectively (Figures S7 and S27). However, in either digest, we did not observe any ^18^O labels at site N196 but detected the unlabeled peptide ^196^NSTPGSSRGTSPA^208^, confirming absence of an N-glycan at N196 (Figure S26). For site N192, we observed neither ^18^O labeled nor unlabeled peptide in any of the enzymatic digests.

### Detection of O-glycosylation on nucleocapsid protein

For the detection of O-glycosylation, we performed LC-MS/MS experiments on tryptic and elastase digests separately and sequentially. We also confirmed O-glycosylation on peptides by removing N-glycans by treatment of PNGase F, followed by enzymatic digestion with trypsin and elastase. We identified the O-glycosylation on the N protein by performing higher energy collisional dissociation (HCD) and collision-induced dissociation (CID) fragmentations, and O-glycosylation site occupancy by a targeted electron-transfer dissociation (ETD) MS^2^ fragmentation on O-glycopeptides. ETD is an ionization technique that induces fragmentation along the glycopeptide backbone generating c and z ions with the intact glycan, which helps pinpointing the position of the glycan in the peptide.^32^ We were able to confirm O-glycosylation on 8 distinct peptides comprising 11 O-glycan sites and determined the site of O-glycan attachment (Figure 6). The glycan occupancy on three glycopeptides that comprise 4 O-glycosylation sites in total was lower (<10%) at each site (Figures S113-S115). We observed ions with the mass corresponding to the peptide ^15^ITFGGPSDSTGSNQNGER^32^ with multiple glycans, such as m/z 2771.1381 for peptide+HexNAc1Hex1NeuAc2 and m/z 3137.2547 for peptide+HexNAc2Hex2NeuAc2. In HCD and CID MS experiments, presence of oxonium ions for HexNAc (m/z 204.0865) and NeuAc (m/z 292.1026) and respective b and y peptide fragment ions confirmed the identity of the glycopeptide (Figures S42-S46). The ETD experiments suggested that glycosylation is present at site S23 (fragments c8, c9, z9 and c10) (Figure S42). However, we observed that the glycan occupancy at S23 was only about 2% as shown in Figure S113. HCD and CID MS^2^ analysis of glycopeptide ^144^DHIGTR^149^ confirmed the O-glycosylation at T148 by the presence of glycan oxonium ions, peptide fragments (b and y ions) and glycan neutral losses (Figures S47-S51). The overall glycan occupancy at the site T148 was determined with ~81%, out of which mono- and di-sialyl Core 2 type glycans (~70%) were the predominant ones (Figure 7B, Table S3). The Byonic glycopeptide search showed a strong indication for the presence of glycan on the peptide ^159^LQLPQGTTLPK^169^. To further evaluate the glycosylation in this peptide, we manually validated full mass, HCD and CID MS^2^ spectra that showed presence of oxonium ions, b and y peptide fragments along with peaks for mass of peptide and peptide+HexNAc (Figure S53). This glycopeptide contains two possible O-glycosylation sites, the ETD experiment for the site mapping of T165 or T166 in the peptide ^159^LQLPQGTTLPK^169^ detected two spectra that showed c and z ions indicating that some of these peptides are O-glycosylated on T165 and others on T166 (Figures S68 and S69). As shown in ETD MS^2^ spectra, the diagnostic peaks z5 at m/z 908.57 and c7 at m/z 1120.57 for the presence of glycan at site T165 were observed (Figure S68). Similarly, the diagnostic peaks z4 at m/z 807.36 and c8 at m/z 1221.65 for presence of glycans at site T166 (Figure S69) were detected. It was found that the peptide ^159^LQLPQGTTLPK^169^ is almost fully occupied with glycans and Core 1 type structures were predominant ones by comprising 77% of total glycans at these sites (Figure 7B). However, we also detected Core 2 to Core 4 type of glycans including extended Core 2 glycan with fucose and sialic acid on the peptide ^159^LQLPQGTTLPK^169^ (Figures S52-S69). The peptide ^204^GTSPAR^209^, located in the LKR region, showed presence of O-glycans at both the sites, T205 and S206. Interestingly, this peptide ^204^GTSPAR^209^ is about 85% occupied with sialylated Core 1 and Core 2 types of glycans in total (Figures 7B and S70 – S76). The Byonic and manual glycoproteomics search also found presence of the peptide ^238^GQQQQGQTVTK^248^ at m/z 1858.8428 with glycans HexNAcHexNeuAc that contains sites T245 and T247 at the CTD region (Figure S79). Based on full mass and fragment ions, another peptide ^376^ADETQALPQR^385^ that contains O-glycosylation site T379 showed indication of presence of glycan (Figures S81-S90). Nonetheless, the peptides ^238^GQQQQGQTVTK^248^ and ^376^ADETQALPQR^385^ were mostly unoccupied (>90%, Figures S114-S115). Towards the C-terminus end of N protein, the peptide ^388^KQQTVTLLPA^397^ which includes O-glycosylation sites T391 and T393, was found with attached glycans. The ETD experiments detected corresponding c and z peptide fragment with intact glycan suggesting that the site T391 is glycosylated (Figure 5D). At site T391 di-sialylated Core 1 to Core 2 type glycans were predominant over other glycans (Figures 7B and S91-S105, Table S4). Interestingly, site S404 was found to contain only mono- and di-sialylated Core 2 type structures as shown in Figures S106-S108. We confirmed the O-glycan structures and glycosylation sites through both Byonic software search, as well as manual analysis. In summary, as shown in Figure 7B the sites T148, T165, T166, T205, S206, T391, and S404 showed the presence of significant levels of O-glycosylation (47%-90%) and the sites S23, T245, T247 and T379 indicated a lower level (1%-9%) of O-glycosylation (Figures S113-S115). We identified Core 1, Core 2, Core 3, and Core 4 type O-glycans at these sites, confirming the glycomics analysis findings (Figure 3B and Table S2).

**Figure 3.**
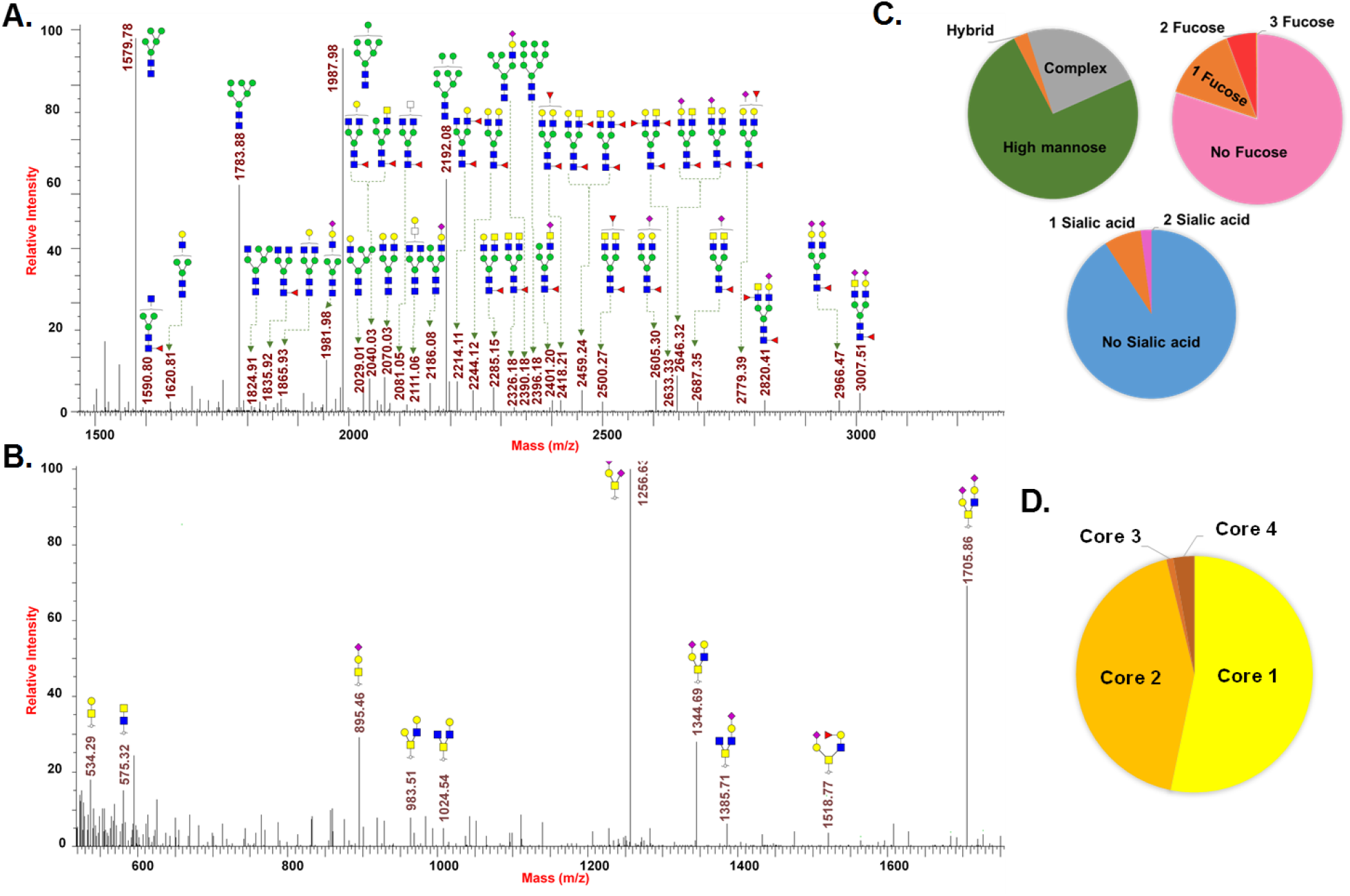
N- and O-glycomics analysis of N protein: **A.** ESI-MS spectrum of permethylated N-glycans released by PNGase F; **B.** ESI-MS spectrum of permethylated O-glycans released by β-elimination; **C.** Relative percentages of N-glycan types, fucosylation and sialylation; **D.** Relative percentage of O-glycan types.

### Phosphorylation on nucleocapsid protein

N proteins from coronaviruses such as SARS-CoV-1 (Uniprot ID P59595) and MERS-CoV (Uniprot ID K9N4V7, uniprot.org) have been known as phosphoproteins.^33^ However, to the best of our knowledge, there are no reports of site-specific phosphorylation for SARS-CoV-2, therefore we included the identification of phosphorylation sites present on SARS-CoV-2 N protein.^13^ We conducted the Byonic based phosphate search, along with O-glycans in the elastase digest of the N protein sample. In this experiment we identified phosphorylation on glycopeptide ^388^KQQTVTLLPA^397^ that contains two potential sites – T391 and T393, for glycosylation and phosphorylation. As shown in the HCD MS^2^ spectrum (Figure S110), the glycopeptide identity was confirmed by validating b and y ions and glycan neutral losses. The strong peaks for presence of peptide, peptide+HexNAc and peptide+2HexNAc at m/z 1098.65, 1301.73 and 1504.81 further confirmed the glycosylation on this peptide. The presence of a phosphate in the peptide was confirmed by CID MS^2^ experiments using the high-resolution precursor mass and neutral losses of phosphate (loss of m/z 79.9663) from the glycopeptide (Figures 4C and S111). As shown in the deconvoluted CID MS^2^ spectrum (Figure S11), the peak at m/z 1584.77 represents the glycopeptide fragment ^388^KQQTVTLLPA^397^ with a phosphate group and glycan, and the peak at m/z 1504.81 represents the same peptide with only glycan showing neutral loss of one phosphate group of 79.9663 Da. Similarly, peaks at m/z 975.45 and at 935.98 show loss of a phosphate group from the peptide fragment bearing a HexNAc3Hex glycan. In the site mapping ETD experiments of this peptide, we observed characteristic c4, c6 and z5, z7 ions at m/z 1565.76, 1846.15, 578.26 and 1840.85, respectively, to unambiguously assign the site of phosphorylation at site T393 and glycan at site T391 (Figures 4D and S109). Interestingly, the peptide with phosphorylation modification also has Core 1 to Core 4 type O-linked glycosylation, including fucosylated glycans, as shown in Figure 7B and Table S4. Additionally, we identified a longer peptide with phosphorylation on glycopeptide ^389^QQTVTLLPAADLDDFSK^405^ that contains three potential sites – T391, T393 and S404 – confirming the glycosylation and phosphorylation. The presence of a phosphate in the peptide was confirmed by CID MS2 experiments using the high-resolution precursor mass and neutral losses of phosphate (loss of m/z 79.9663) from the glycopeptide (Figure S112). As shown in the deconvoluted CID MS2 spectrum in Figure S112, the peak at m/z 2964.30 represents the glycopeptide fragment ^389^QQTVTLLPAADLDDFSK^405^ bearing the glycan HexNAc2Hex2NeuAc1 and phosphate, and peak at m/z 2884.34 represents the same peptide with glycan HexNAc2Hex2NeuAc1 showing neutral loss of one phosphate group.

**Figure 4.**
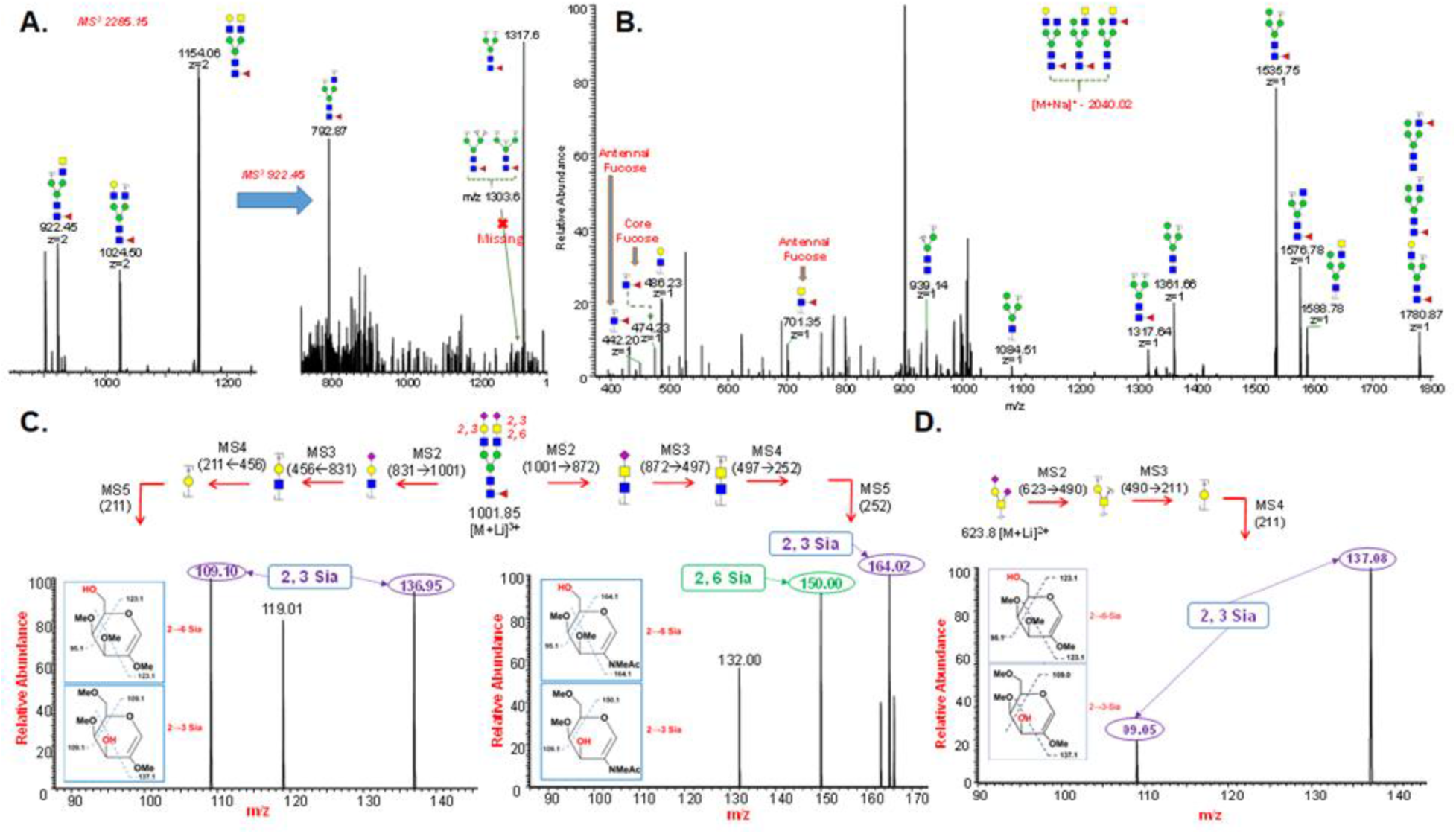
ESI-MSn analysis was performed on permethylated N- and O-glycans for the structure and linkage determination. **A.** ESI-MS2 spectrum of permethylated N-glycan of N protein with mass 2285.15 and ESI-MS3 of fragment 922.45 showing absence of triantennary and bisecting GlcNAc structures. This also provided the confirmation for the presence of terminal GalNAc on N-glycans; **B.** ESI-MS2 spectrum of permethylated N-glycan of N protein showing both core and antennal fucosylation. **C.** ESI-MS5 spectrum of permethylated N-glycan of N protein for sialic acid linkage analysis; **D.** ESI-MS5 spectrum of permethylated O-glycan of N protein for sialic acid linkage analysis.

**Figure 5.**
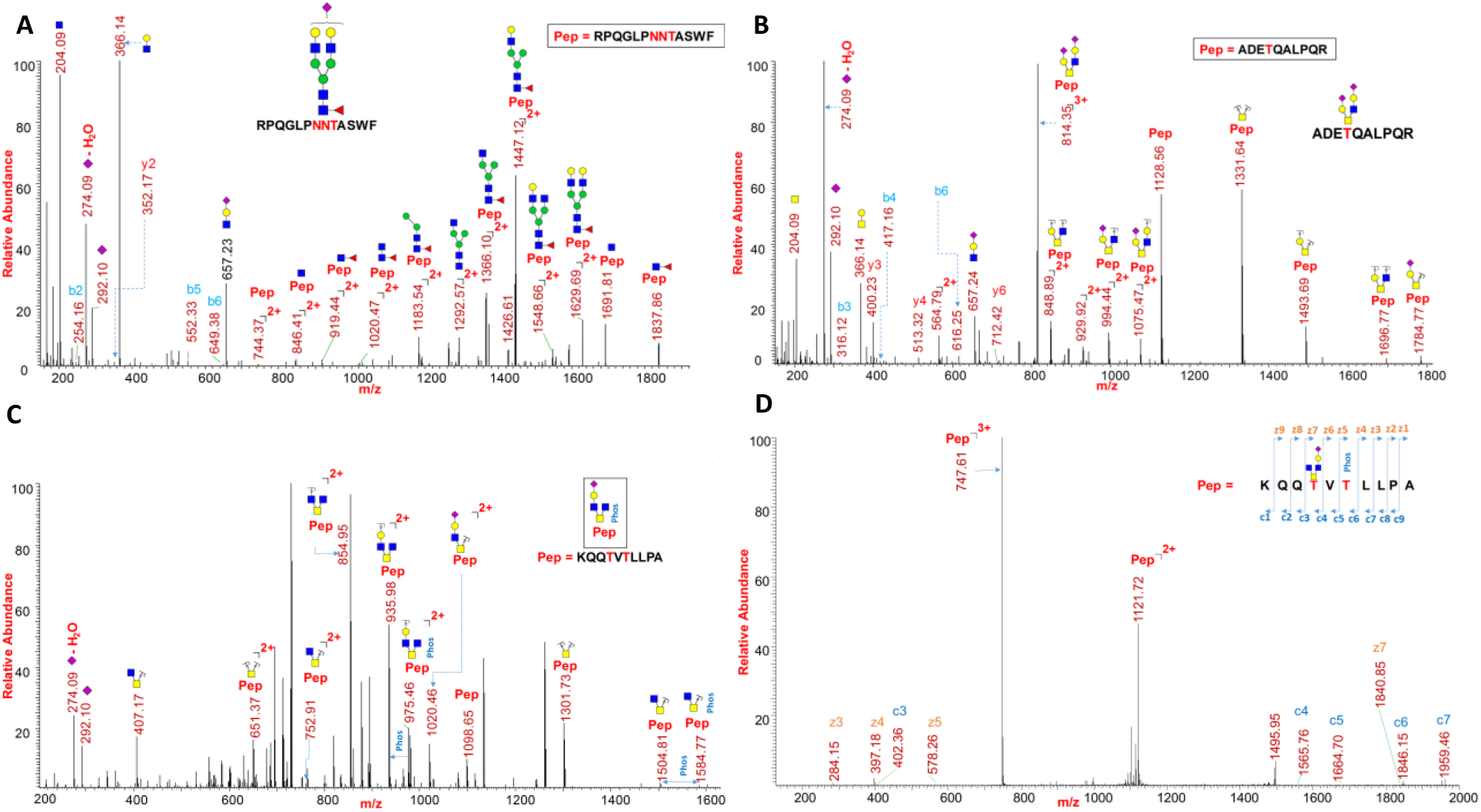
PTM analysis of N protein via proteomics, glycoproteomics, and phosphoproteomics. **A.** HCD MS2 spectrum of N-glycopeptide 41RPQGLPNNTASWF53 containing HexNAc(4)Hex(5)Fuc(1)NeuAc(1); **B.** HCD MS2 spectrum of O-glycopeptide 376ADETQALPQR385 containing HexNAc(2)Hex(2)NeuAc(1) at site T379; **C.** CID MS2 spectrum of O-glycophosphopeptide 388KQQTVTLLPA397 containing HexNAc(3)Hex(1)NeuAc(1) and phosphorylation at T391 and T393 showing neutral loss of phosphate group; **D.** ETD MS2 spectrum of O-glycophosphopeptide 388KQQTVTLLPA397 showing presence of HexNAc(3)Hex(1)NeuAc(1) at site T391 and phosphorylation at T393.

**Figure 6.**
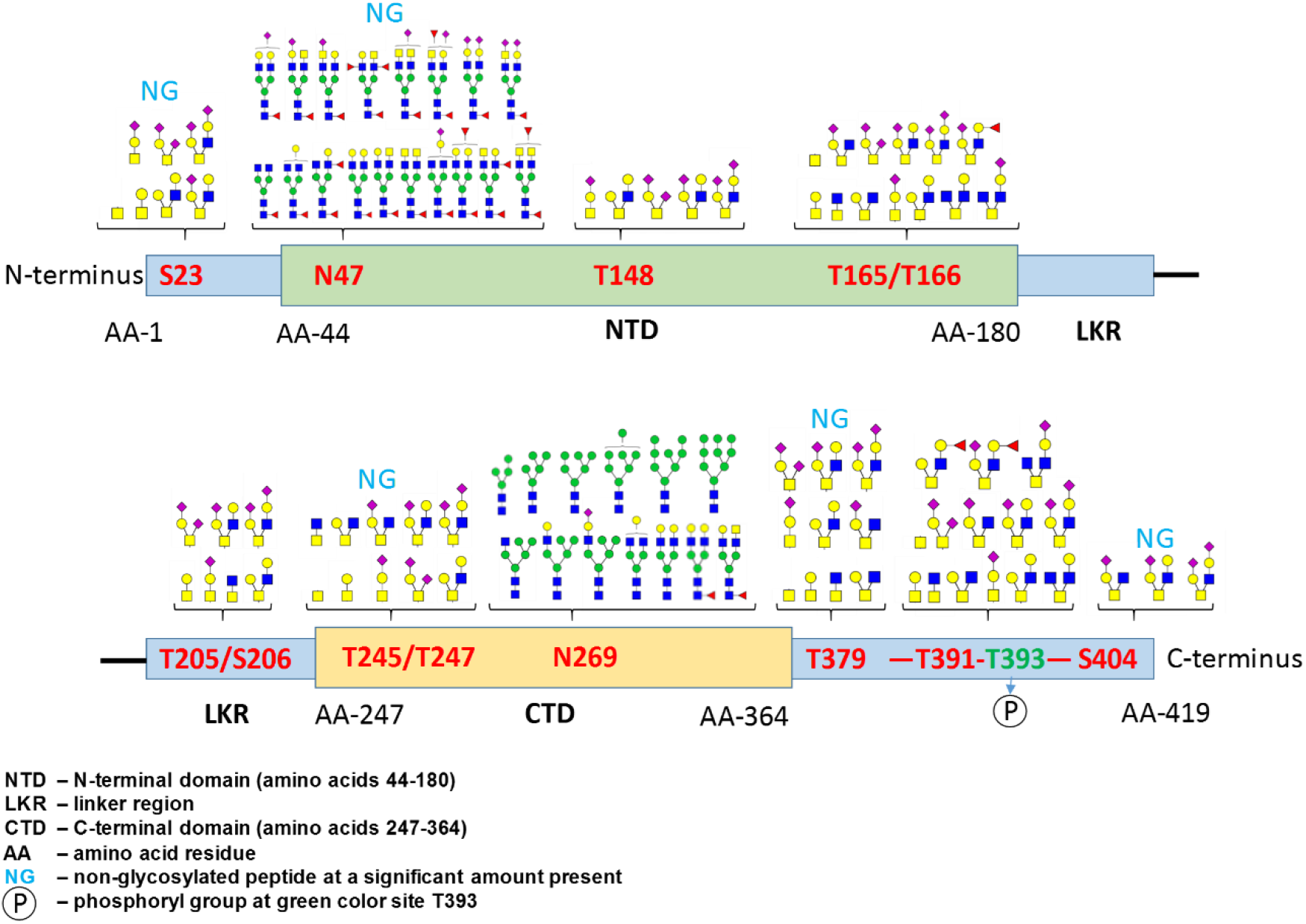
Schematic overview of the PTM modifications identified on N protein of SARS-CoV-2. Complex type N-glycosylation at N47 and major high mannose type of N-glycosylation at site N269 were observed. O-glycosylation sites show Core 1 to Core 4 type of glycans including an O-glycopeptide with phosphorylation site (T393) towards the C terminus of protein.

**Figure 7.**
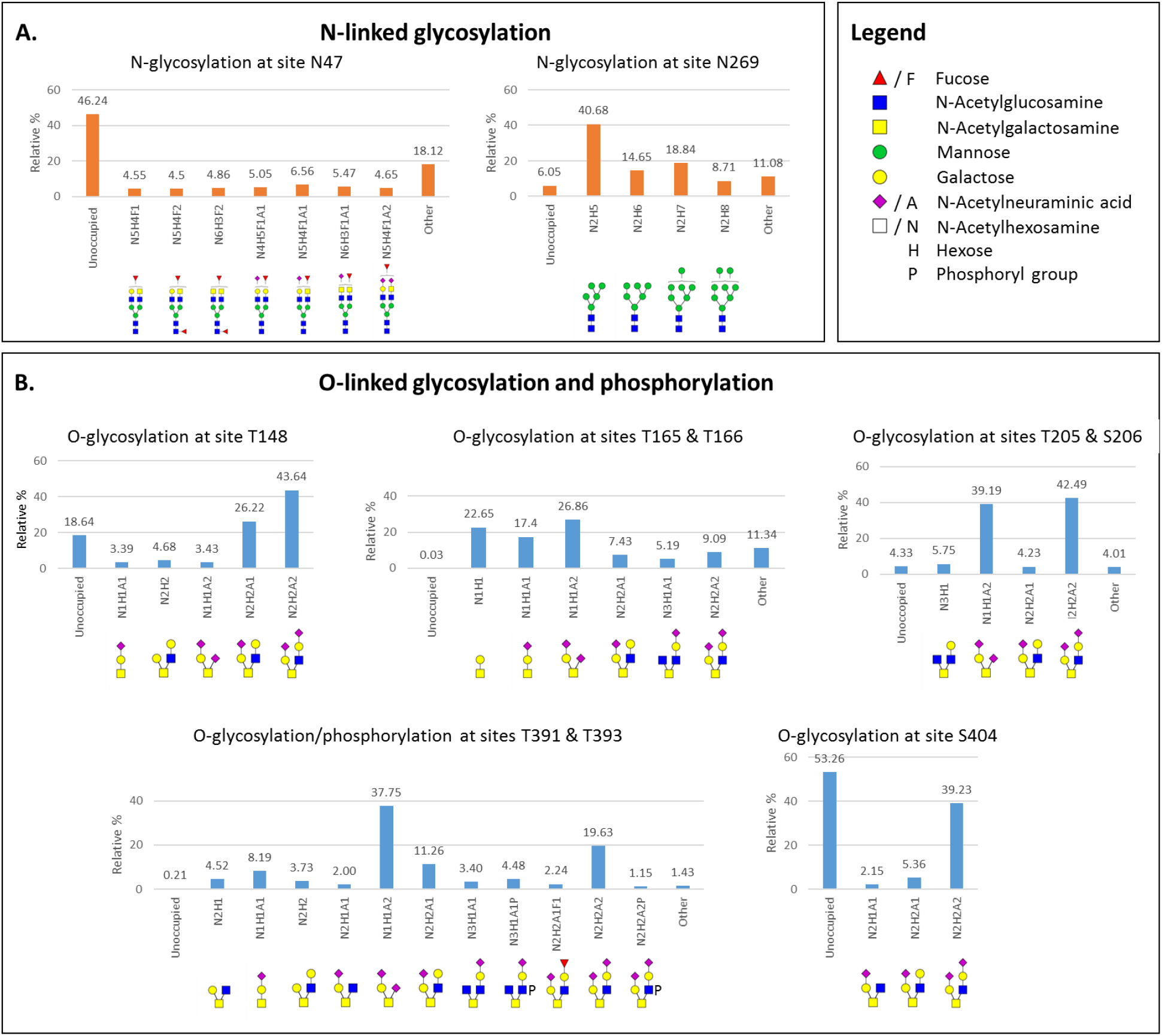
Site specific relative quantification of N- and O-linked glycosylation; **A.** Relative percentage of N-glycans at sites N47 shows complex type glycans and N269 shows majority of high mannose type of glycan with Man5 structure is the most abundant; **B.** Relative percentage of O-glycans at sites T148, T165, T166, T205, S206, T391, and S404. Phosphorylation is detected on T393. These sites showed major glycan occupancy ranging from 47%-99%. Minor O-glycosylated site distribution is shown in SI. Other – minor glycoforms with less than 2 % abundances.

### Sequence alignment of nucleocapsid proteins of SARS-CoV-1, MERS-CoV and SARS-CoV-2

The alignment of the N protein sequences of SARS-CoV-1 (ID P59595, 422 amino acids), MERS-CoV (ID K9N4V7, 411 amino acids) and SARS-CoV-2 (ID P0DTC9, 419 amino acids) via UniProt reveals that the N protein from SARS-CoV-1 and SARS-CoV-2 is highly conserved, sharing about 90% similarity. This is in agreement with other studies that used slightly different sequences/alignments.^34–35^ The N protein sequence of MERS-CoV shares only 42% to 44% similarity with the SARS-CoV-1 and SARS-CoV-2 N protein, respectively.

As shown in Figure 2B, the N protein sequence of SARS-CoV-1 carries 3 N-linked glycosylation motifs (-NXS/T-with X≠P), whereas SARS-CoV-2 carries 2 additional sites – totaling 5 potential N-glycan sites for SARS-CoV-2. MERS-CoV carries 6 N-linked glycosylation motifs in total: Two of the potential N-glycan sites are unique to MERS-CoV, two overlap with motifs found in SARS-CoV-2, and the remaining two motifs are found in all three sequences of MERS-CoV, SARS-CoV-1 and SARS-CoV-2. As mentioned before, neither MERS-CoV nor SARS-CoV-1 have been reported as possessing N-glycans on the N protein in previous studies. Overall alignment of potential N-glycosylation sites among the three N proteins and the data presented in this study indicate the high likelihood of the presence of N-glycans in the N proteins of all three coronavirus species.

## Discussion

The coronavirus N protein consists of three distinct but highly conserved domains, including an N-terminal RNA-binding domain (NTD), a C-terminal dimerization domain (CTD), and a central Ser/Arg (SR)-rich linker domain, which displays an intrinsically disordered structure and facilitates molecular movements to aid interactions. The NTD is reportedly responsible for RNA binding, CTD for oligomerization, and the SR-rich linker is generally known to be primarily involved in phosphorylation events.^36–38^ However, no site specific glycosylation nor phosphorylation was reported in any of these domains in the current literature on SARS-CoV-2. Our LC-MS/MS studies have shown that the N protein contains N- and O-glycosylation, in addition to phosphorylation. The coronavirus N protein-NTD associates with the 3’ end of the viral RNA genome through electrostatic interactions.^39^ In the NTD, we identified one N-glycosylation site (N47) and three O-glycosylation sites (T148, T165 and T166), which possibly play a role in regulating the RNA-binding activity. In the CTD, we observed minor O-glycosylation at sites T245 and T247 and N-glycosylation at site N269. Considering the role of CTD in oligomerization and self-association, future studies on the role of the glycosylation is of vital importance.^40^ The detailed structural analysis of the PTMs on the N protein of the newly emerged SARS-CoV-2 could be a key aspect in the development of therapeutic agents that specifically and efficiently block the coronavirus replication, transcription, and viral assembly.^36^

To our knowledge, the CoV N protein is the only phosphorylated structural protein on the virus, and this phosphorylation has been proposed to play a role in regulating its functions.^41–43^ Evidence of significant conformational changes in the N protein structure due to phosphorylation has been reported.^44^ Importantly, N protein from SARS-CoV-1 have been shown to elicit a well-defined immunological response, which underscores the importance of the N protein as a potent target for a vaccine against COVID-19 infection.^45^ A recent study aimed to investigate effect of early SARS-CoV-2-specific humoral immune responses on disease outcome found that deceased patient had stronger antibody responses towards N protein while survivors had much more stronger antibody response to S protein highlighting the importance of N protein in disease outcome.^46^ Therefore, understanding the N protein glycosylation patterns described in this study may ultimately help in new vaccine design to prevent SARS-CoV-2 infection.

## Materials and methods

Sequencing-grade modified trypsin and elastase were purchased from Promega (Madison, WI). Peptide-N-Glycosidase F (PNGase F) was purchased from New England Biolabs (Ipswich, MA). All other reagents were purchased from Sigma Aldrich unless indicated otherwise. Data analysis was performed using Byonic 2.3 software and manually using Xcalibur 4.2 and GlycoWorkbench 1.1. The N protein expressed in HEK293 cells (Cat. No. NUN-C5227) was purchased from AcroBiosystems (Newark, DE).

### N- and O-linked glycan release, purification, and permethylation

The N protein was purified using SDS-PAGE prior to glycomic analysis (Figure S1). The 60-65 kDa gel band was cut into smaller pieces (1 mm squares approx.) and transferred to clean microcentrifuge tubes. The gel pieces were de-stained by adding 500 μL acetonitrile (ACN): 50mM NH4HCO3 (1:1) and incubated at room temperature (RT) for about 30 min. Tubes were centrifuged, and the supernatant was discarded. Then 250 μL ACN was added, and the gel pieces were incubated for 20-30 min. Samples were centrifuged, and supernatant was discarded. The gel pieces were then suspended in 500 μL 50mM NH4HCO3 and 3 μL PNGase F added. The sample was incubated for 18 h at 37 °C. The released N-glycans were extracted out by adding 1:2 H2O: ACN (500 μL) at room temperature for 15 min, and the supernatant was collected. The released N-glycans were speed dried, dissolved in with 0.1% formic acid solution (FA), purified by passing through a C18 SPE cartridge and eluted with 0.1% formic acid solution (3 mL) and dried by lyophilization. The gel pieces containing O-glycoprotein were digested by adding sequence grade trypsin in digestion buffer (50 mM NH4HCO3) for 18 hr. at 37 °C. The O-glycopeptides were extracted out from gel by the addition of 1:2, H2O: ACN containing 5% formic acid (500 μL) and kept at RT for 15 min and supernatant was collected. The released O-glycopeptides were speed dried and used for the release of O-glycans by β-elimination.

The O-glycans were released from the glycopeptide peptide by reductive *β*-elimination reported elsewhere.^32^ Briefly, eluted N protein glycopeptides were treated with a solution of 19 mg/500 μL of sodium borohydride in 50mM sodium hydroxide. The reaction mixture was heated to 45°C for 16 h, then neutralized with a solution of 10% acetic acid. The sample was desalted on a hand-packed ion exchange resin (DOWEX H+) by eluting with 5% acetic acid and dried by lyophilization. The borates were removed by the addition of a solution of methanol:acetic Acid (9:1) and evaporation under a steam of nitrogen.

The released N- and O-linked glycans were then permethylated using NaOH/DMSO-methyl iodide method published previously.^32^

### N- and O-linked glycomic profiling by MALDI-MS and ESI-MS^n^

The permethylated N- and O-glycans were dissolved in 2 μL of methanol. 0.5 μL of sample was mixed with equal volume of DHB matrix solution (10 mg/mL in 1:1 methanol-water) and spotted on to a MALDI plate. MALDI-MS spectra were acquired in positive ion and reflector mode using an AB Sciex 5800 MALDI-TOF-TOF mass spectrometer.

2 μL of permethylated N- and O-glycans dissolved in methanol were mixed with 98 μL of ESI-MS infusion buffer (1:1:1 – 0.1% formic acid in water (with 1 mM NaOH): methanol: acetonitrile) and infused directly into an Orbitrap Fusion Tribrid mass spectrometer through a nanospray ion source. ESI-MS^n^ spectra of glycans were acquired by both total ion monitoring and manual MS^n^ fragmentation.

Original glycoform assignments were made based on full-mass molecular weight. Additional structural details were determined by ESI-MS^n^ and analysis with GlycoWorkbench 1.1 software.

### Protease digestion of N protein for glycoproteomics

The purified N protein (20 μg) expressed on HEK293 cells was dissolved in 50 mM ammonium bicarbonate solution and digested with trypsin and elastase separately as well as sequentially by incubating for 1 h at 37 °C. The digest was filtered through 0.2 μm filter and directly analyzed by LC-MS/MS.

### ^18^O labeling via de-glycosylation and subsequent protease digestion of N protein

1 μg of N protein was dissolved in 36 μL H2^18^O and 2 μL 10x glycobuffer (NEB). To this solution 2 μL PNGase F was added and the reaction mixture was incubated at 37 °C for 16 h. Enzymatic activity of PNGase F was deactivated by heating mixture to 95 °C for 5 min followed by lyophilizing the sample. The de-N-glycosylated protein was then suspended 48 μL 50mM NH4HCO3 and added 2 μL Elastase (0.5 μg/μL). The protein was then digested at 37 °C for 16 h. The enzyme was deactivated by heating to 95 °C for 5 min and solvents were removed by speed vacuum. The peptides were reconstituted in 0.1% FA, filtered through 0.2 μm filter and analyzed via LC-MS/MS.

### Data acquisition of protein digests using nano-LC-MS/MS

The glycoprotein digests were analyzed on an Orbitrap Fusion Tribrid mass spectrometer equipped with a nanospray ion source and connected to a Dionex binary solvent system (Thermo Fisher, Waltham, MA). Pre-packed nano-LC column (Cat. No. 164568, Thermo Fisher, Waltham, MA) of 15 cm length with 75 μm internal diameter (id), filled with 3 μm C18 material (reverse phase) were used for chromatographic separation of samples. The precursor ion scan was acquired at 120,000 resolution in the Orbitrap analyzer and precursors at a time frame of 3 s were selected for subsequent MS/MS fragmentation in the Orbitrap analyzer at 15,000 resolution. The LC-MS/MS runs of each digest were conducted for 180 min. 0.1% formic acid and 80% acetonitrile-0.1 % formic acid was used as mobile phase A and B, respectively, in order to separate the glycopeptides. The threshold for triggering an MS/MS event was set to 1000 counts, and monoisotopic precursor selection was enabled. MS/MS fragmentation was conducted with stepped HCD (Higher-energy Collisional Dissociation) product triggered CID (Collision-Induced Dissociation) (HCDpdCID) program. The O-linked glycopeptides were also analyzed for site mapping by targeted Electron Transfer Dissociation (ETD) and acquired on both Orbitrap and ion trap analyzers. Charge state screening was enabled, and precursors with unknown charge state or a charge state of +1 were excluded (positive ion mode). Dynamic exclusion was also enabled (exclusion duration of 30 s).

### Data analysis of glycoproteins

The LC-MS/MS spectra of combined tryptic /elastase digest of N protein were searched against the FASTA sequence of N protein using the Byonic software 2.3 by choosing appropriate peptide cleavage sites (semi-specific cleavage option enabled). Oxidation of methionine, deamidation of asparagine and glutamine, and possible common mammalian N-glycans, O-glycan and phosphorylation masses were used as variable modifications. The LC-MS/MS spectra were also analyzed manually for the glycopeptides with the support of the Thermo Fisher Xcalibur 4.2 software, GlycoMod tool, and ProteinProspector v6.2.1. The HCDpdCID and ETD MS^2^ spectra of glycopeptides were evaluated for the glycan neutral loss pattern, oxonium ions and glycopeptide fragmentations to assign the sequence and the presence of glycans in the glycopeptides.

## Data Availability

The raw data files and search results can be accessed from glycopost repository - https://glycopost.glycosmos.org/preview/11350896155f46ae870bcc3; Pin: 2697.

## Acknowledgements

Financial support from the US National Institutes of Health (S10OD018530) is gratefully acknowledged.

## Abbreviations

N protein: Nucleocapsid protein
M: membrane
E: envelope
S: Spike
SARS-CoV-2: severe acute respiratory syndrome coronavirus 2
NTD: N-terminal RNA-binding domain
CTD: C-terminal dimerization domain
COVID-19: coronavirus disease
hACE2: human angiotensinconverting enzyme 2
HEK: human embryonic kidney
DTT: dithiothreitol
PNGase F: Peptide-N-Glycosidase F
ACN: acetonitrile
HCD: Higher-energy Collisional Dissociation
pd: product triggered
CID: Collision-Induced Dissociation
HCD: Higher-energy Collisional Dissociation
ETD: Electron-transfer dissociation
IT-MS: ion trap-mass spectrometry

## Contributions

P.A., N.S and A.S. conceived of the paper; N.S. performed glycoproteomics and glycomics sample processing, A.S conducted glycomics data acquisition, N.S, A.S, A.G and D.A. performed data analysis; everyone contributed toward writing the manuscript; P.A. and C.H. monitored the project.

## Supplemental material

SDS-PAGE-purification (S1), Glycomics and Glycoproteomics workflow of N protein (S2), N- and O-glycan MALDI/ESI-MS data (S3-S5), Annotated glycopeptide MS/MS spectra (S6 – S112), O-Glycoform quantification for S23, T245, T247 and T379 – (S113-S115) and are incorporated as a separate supplemental material file. Supplemental Tables 1-4 are also included as separate files for relative quantification via glycomics and glycoproteomics.

## Supplemental Data Included

Figures S1 – S115, Supplemental Tables 1-4

