## Supplemental Figures for "SARS-CoV-2 Nucleocapsid protein is decorated with multiple N- and O-glycans"

Supplemental Data Included: Figures S1 – S115, Supplemental Tables 1-4

Running Title: Glycosylation of Nucleocapsid (N) protein

**Supplementary Figures:**

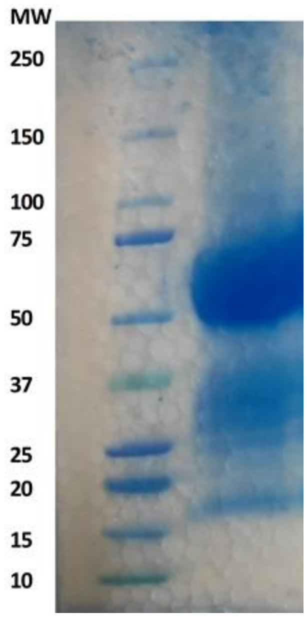


**Supp. Figure S1** SDS PAGE purification of N-protein

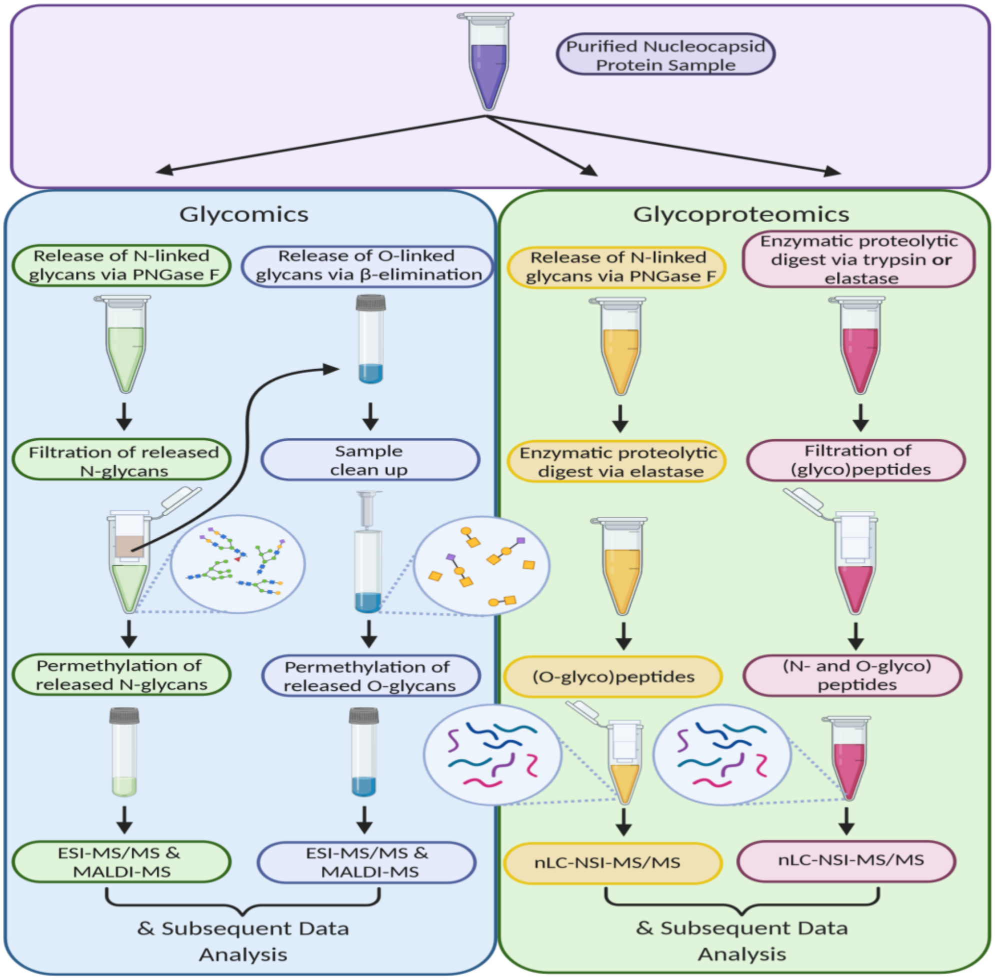


**Supp. Figure S2** Glycomics and Glycoproteomics workflow of N protein

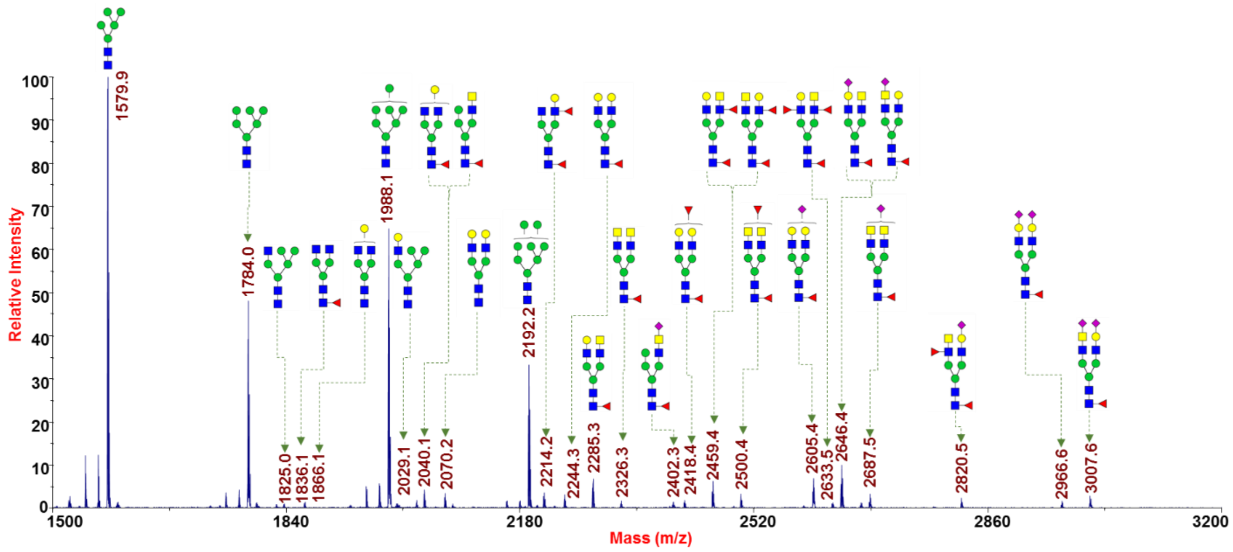


**Supp. Figure S3** MALDI-MS spectrum of permethylated N-glycans released by PNGase F

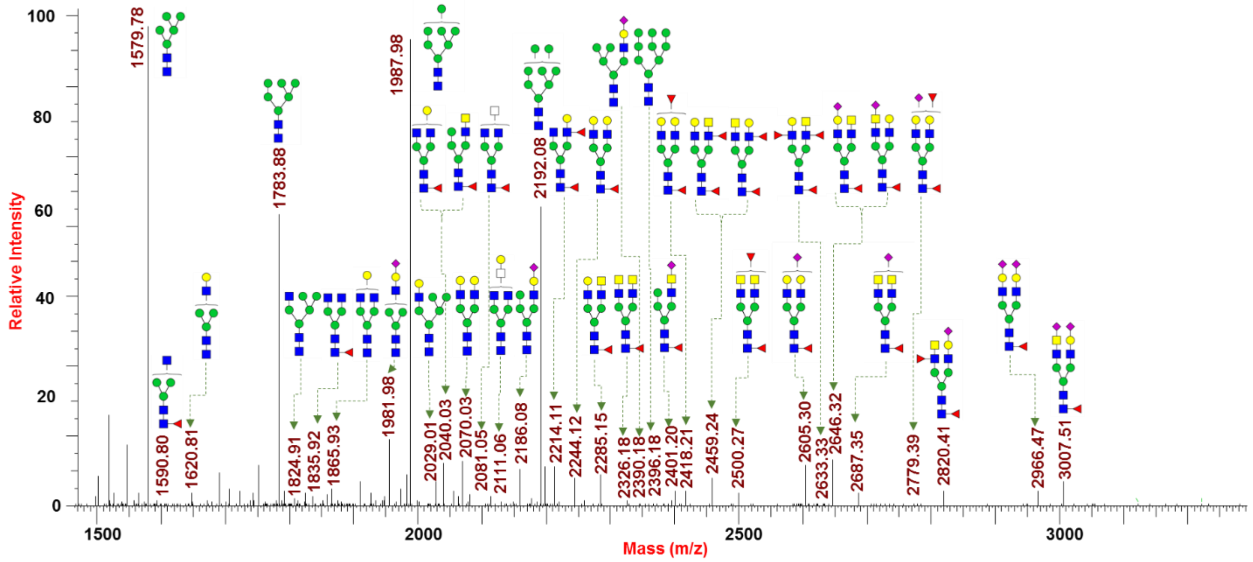


**Supp. Figure S4** Full ESI-MS spectrum of permethylated N-glycans released by PNGase F

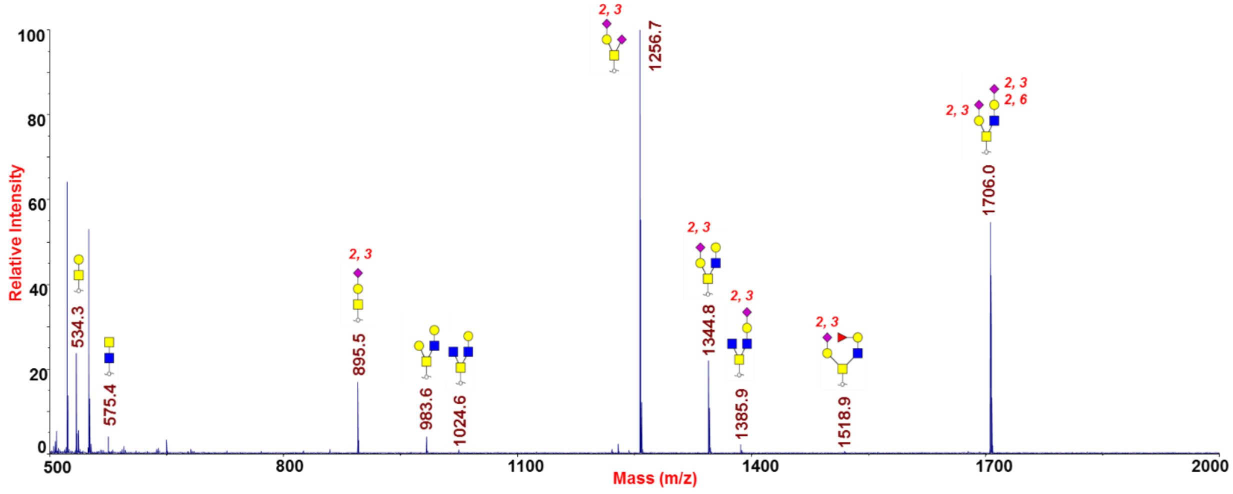


**Supp. Figure S5** MALDI-MS spectrum of permethylated O-glycans released by β-elimination.

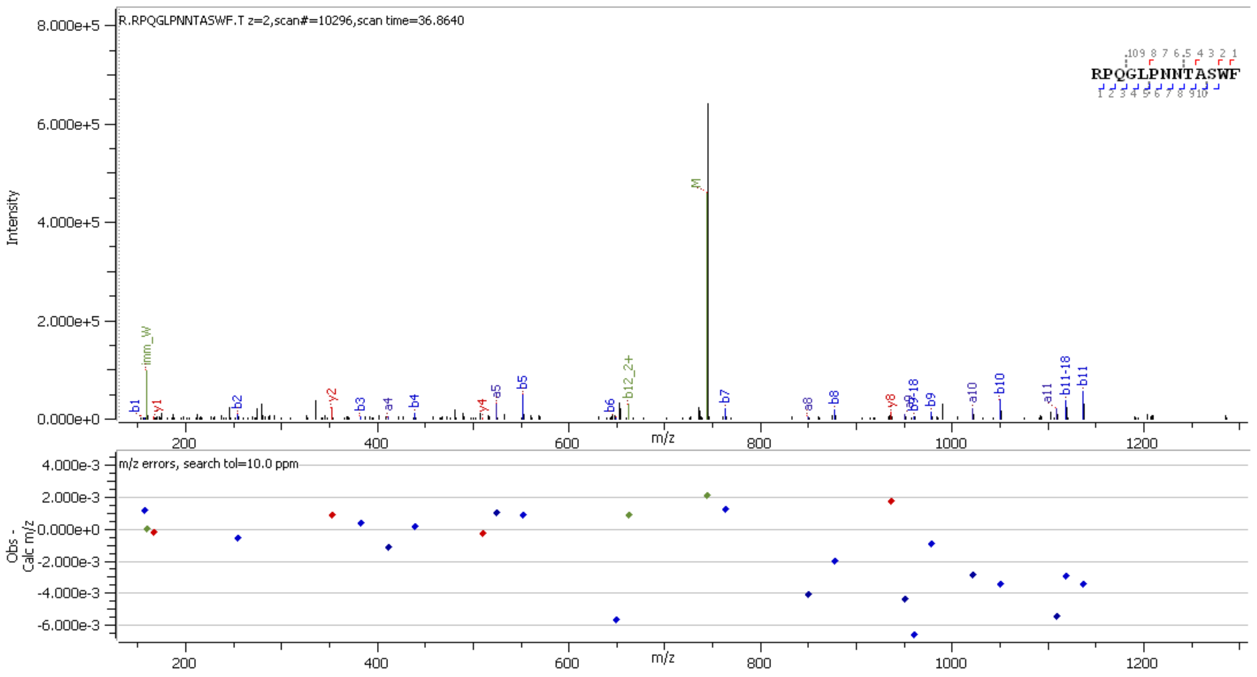


**Supp. Figure S6** HCD MS2 spectrum of unglycosylated peptide $^{41}$RPQGLPNNTASWF$^{53}$

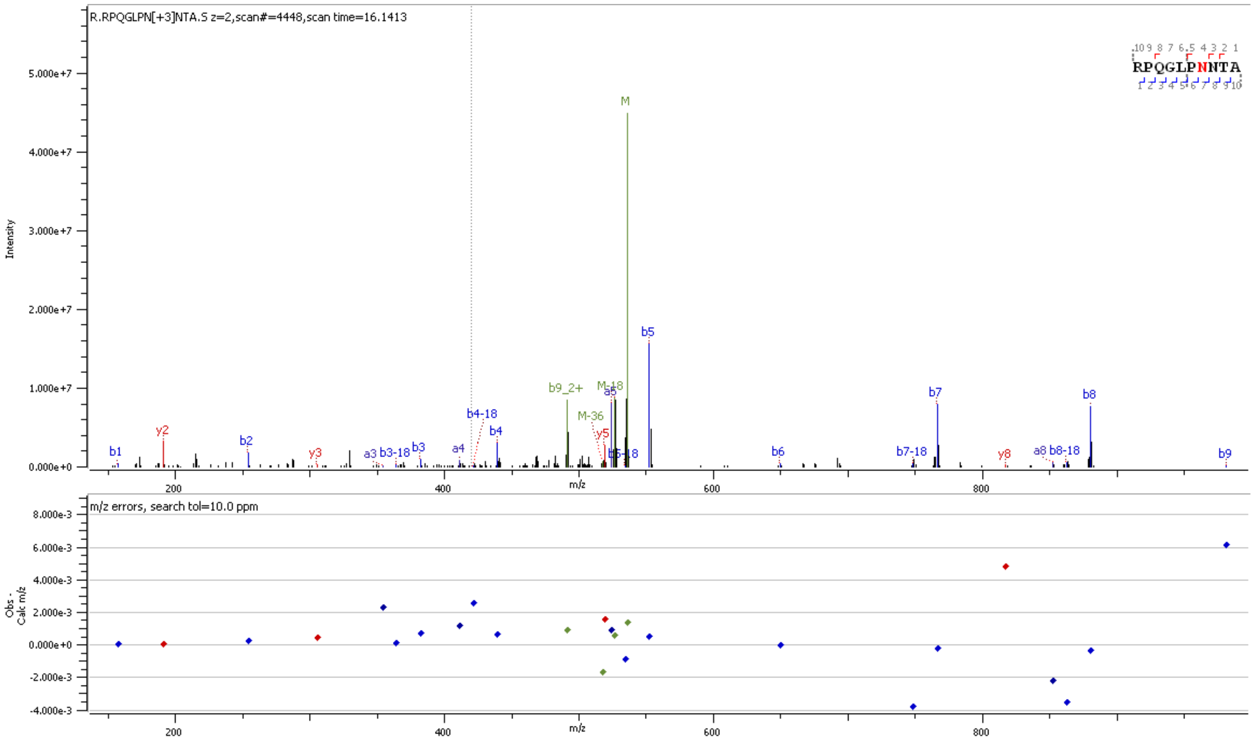


**Supp. Figure S7** HCD MS2 spectrum of peptide [41]RPQGLPNNTA[50] showing presence of [18]O-Asp at position 47

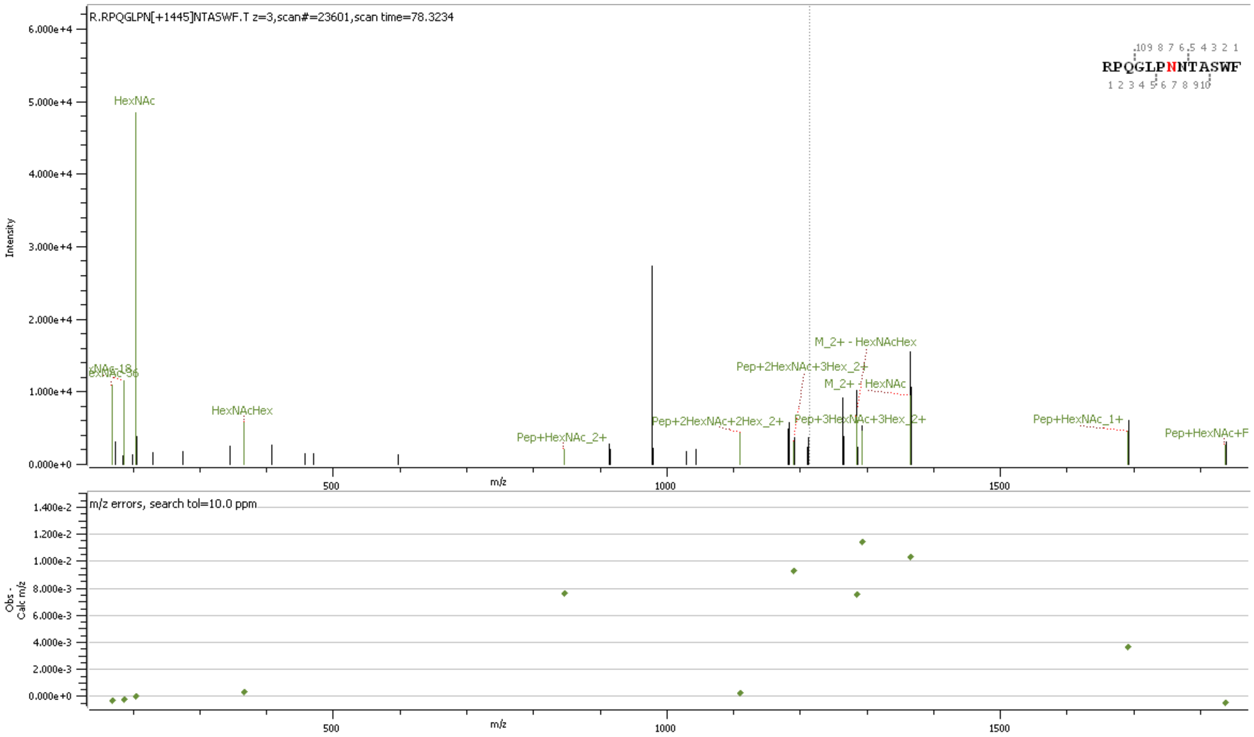


**Supp. Figure S8** HCD MS2 spectrum of N-glycopeptide [41]RPQGLPNNTASWF[53] containing HexNAc(4)Hex(3)Fuc(1)

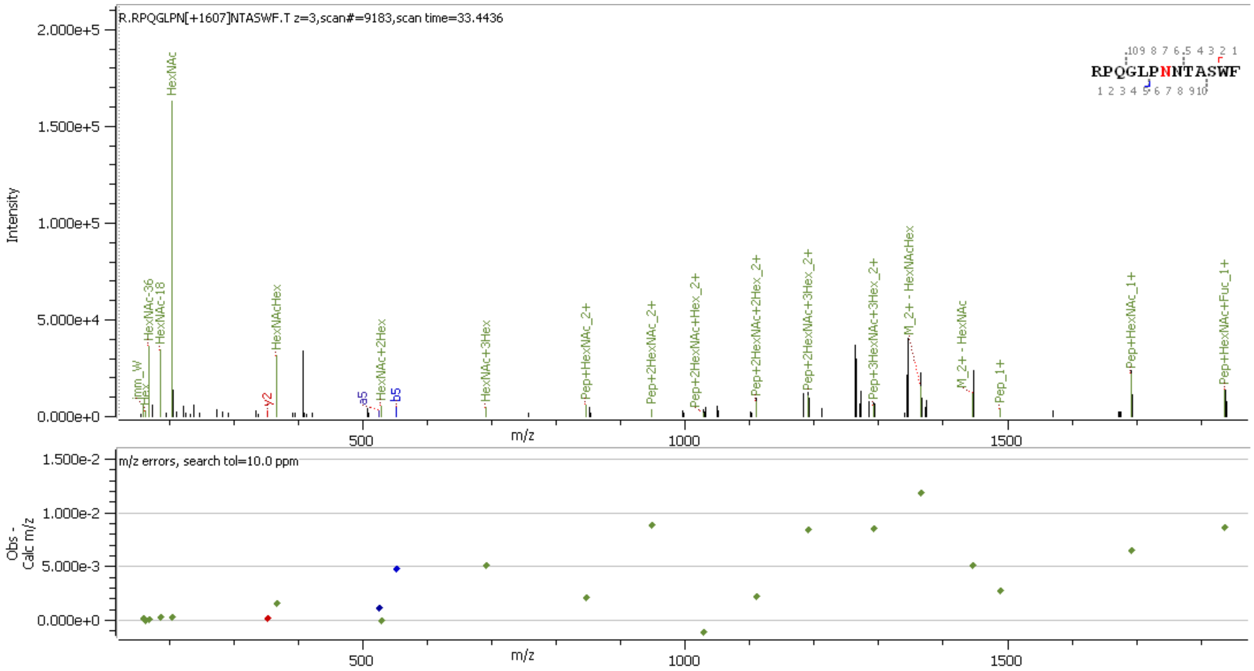


**Supp. Figure S9** HCD MS2 spectrum of N-glycopeptide [41]RPQGLPNNTASWF[53] containing HexNAc(4)Hex(4)Fuc(1)

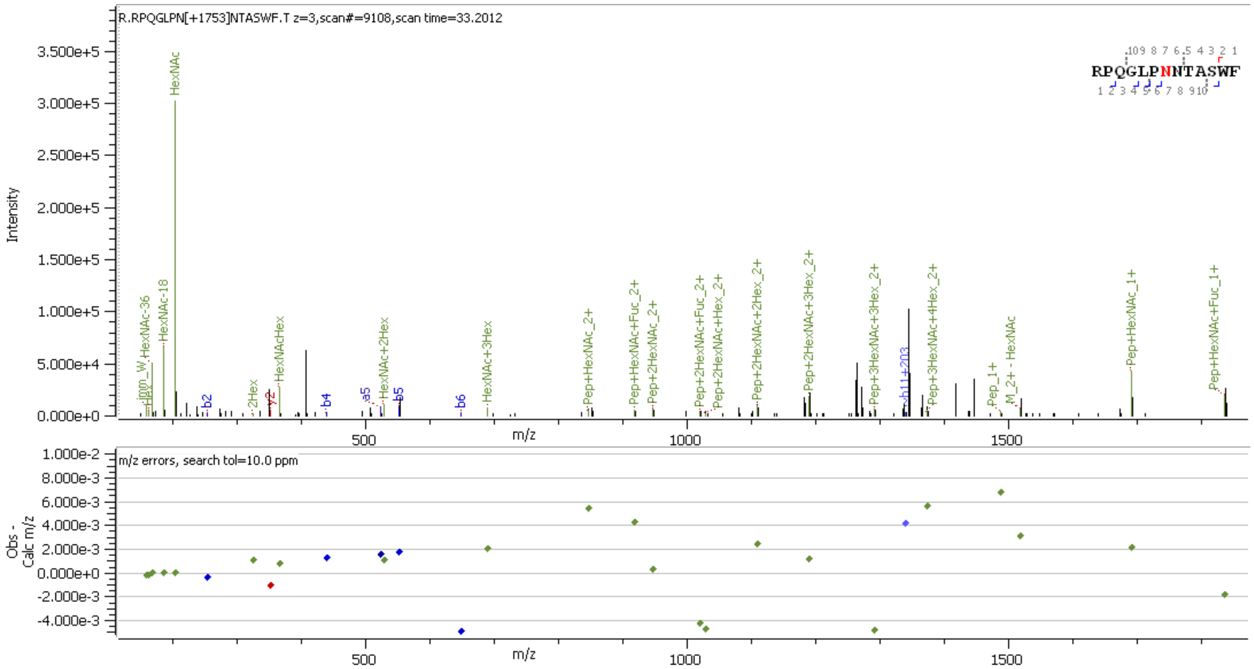


**Supp. Figure S10** HCD MS2 spectrum of N-glycopeptide [41]RPQGLPNNTASWF[53] containing HexNAc(4)Hex(4)Fuc(2)

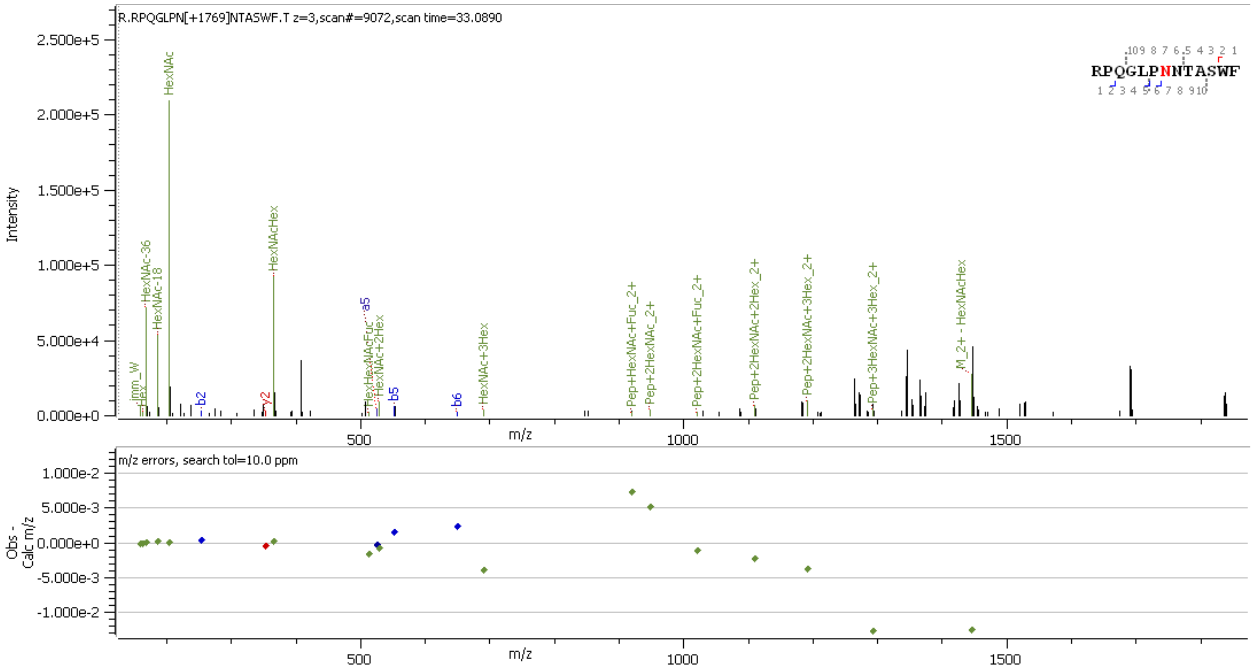


**Supp. Figure S11** HCD MS2 spectrum of N-glycopeptide [41]RPQGLPNNTASWF[53] containing HexNAc(4)Hex(5)Fuc(1)

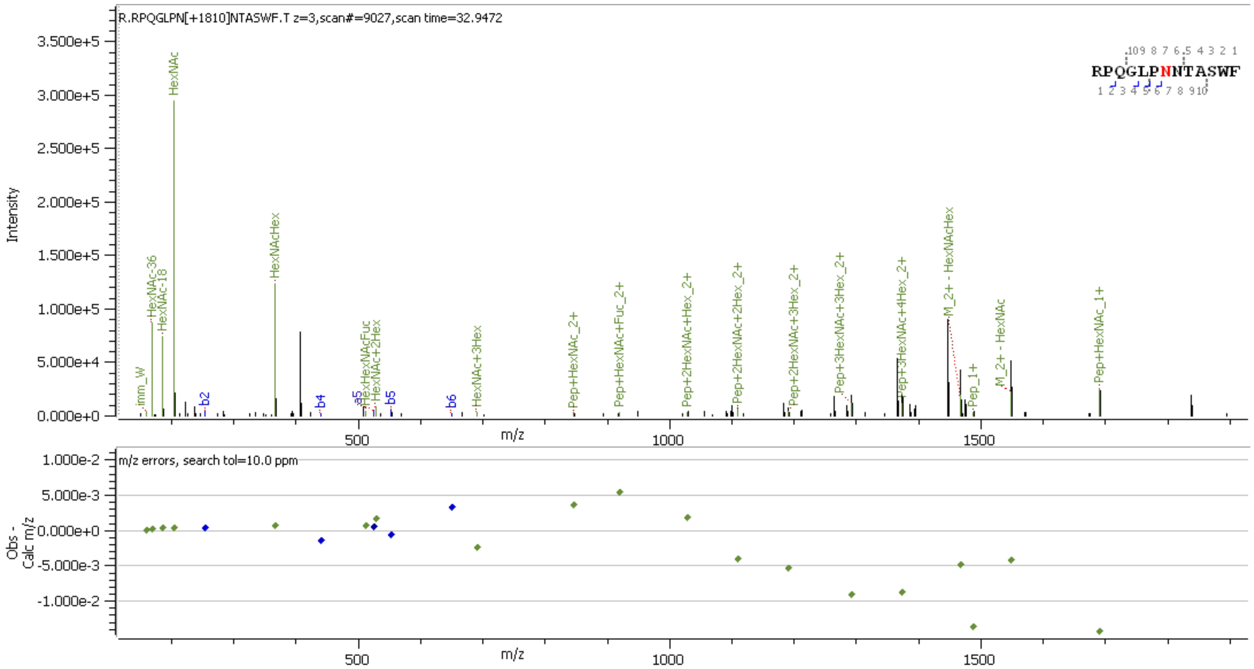


**Supp. Figure S12** HCD MS2 spectrum of N-glycopeptide [41]RPQGLPNNTASWF[53] containing HexNAc(5)Hex(4)Fuc(1)

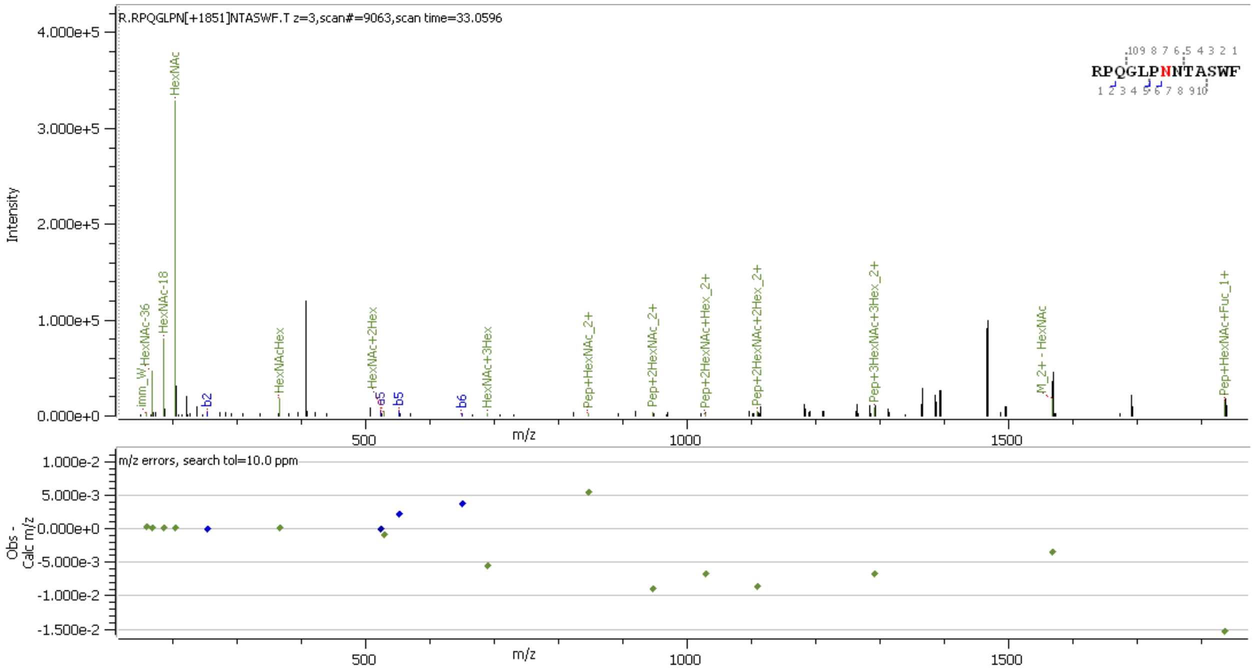


**Supp. Figure S13** HCD MS2 spectrum of N-glycopeptide [41]RPQGLPNNTASWF[53] containing HexNAc(6)Hex(3)Fuc(1)

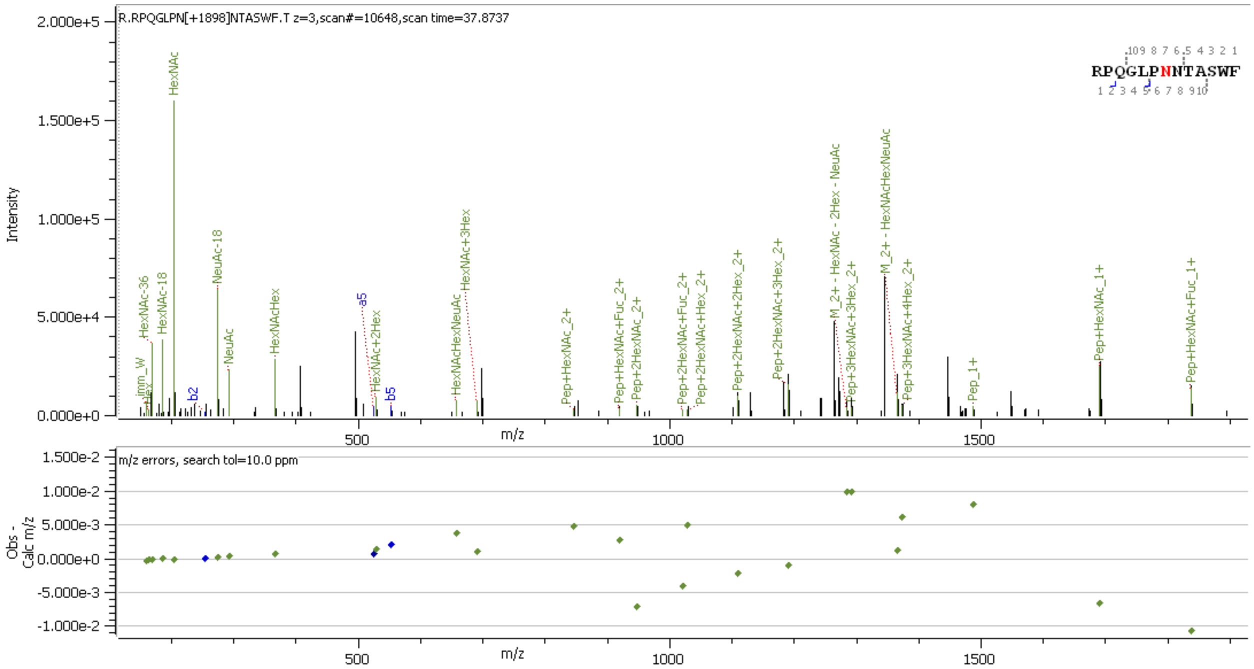


**Supp. Figure S14** HCD MS2 spectrum of N-glycopeptide [41]RPQGLPNNTASWF[53] containing HexNAc(4)Hex(4)Fuc(1)NeuAc(1)

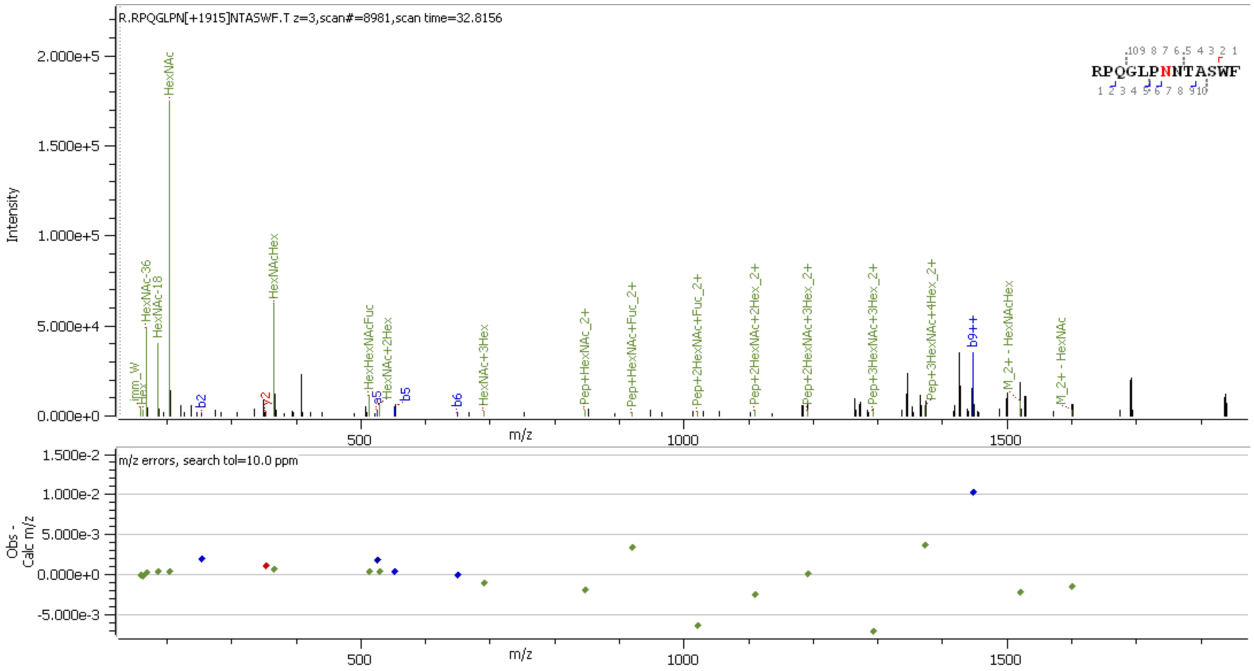


**Supp. Figure S15** HCD MS2 spectrum of N-glycopeptide [41]RPQGLPNNTASWF[53] containing HexNAc(4)Hex(5)Fuc(2)

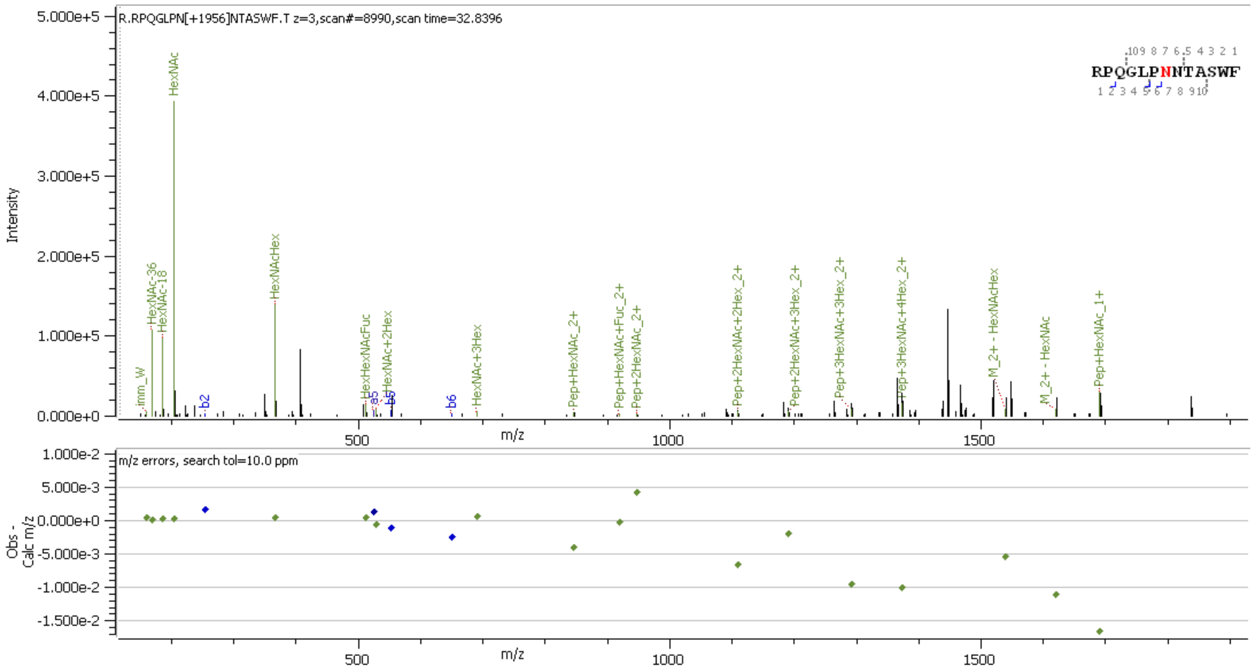


**Supp. Figure S16** HCD MS2 spectrum of N-glycopeptide [41]RPQGLPNNTASWF[53] containing HexNAc(5)Hex(4)Fuc(2)

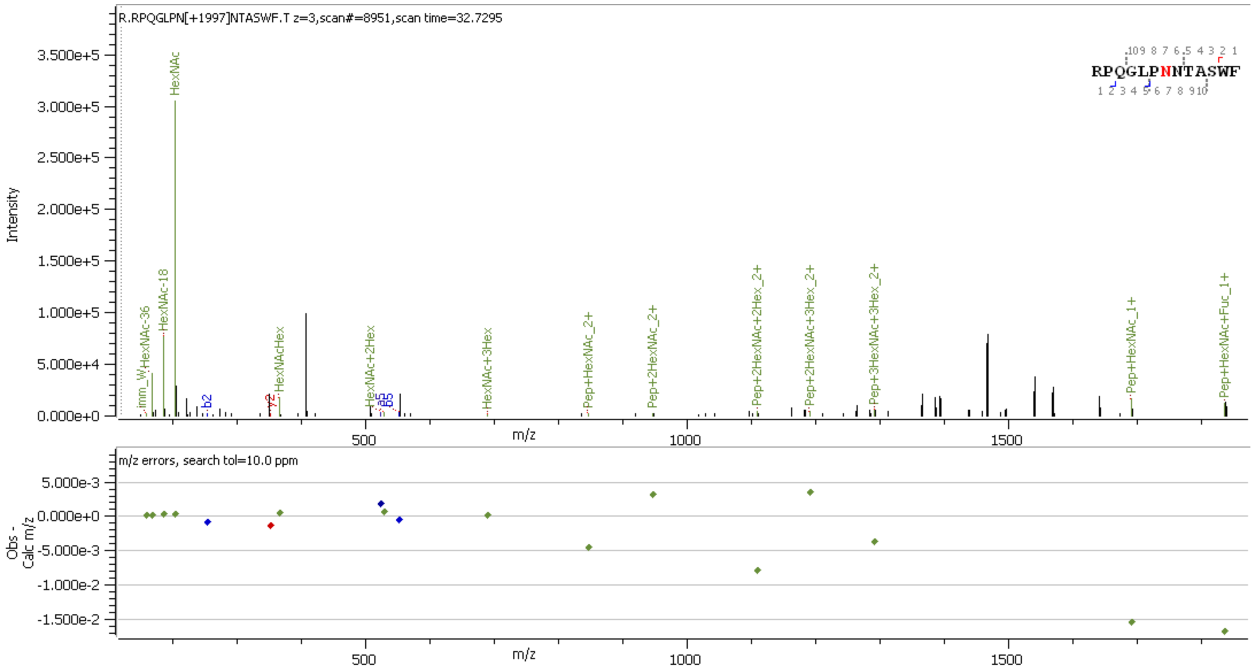


**Supp. Figure S17** HCD MS2 spectrum of N-glycopeptide [41]RPQGLPNNTASWF[53] containing HexNAc(6)Hex(3)Fuc(2)

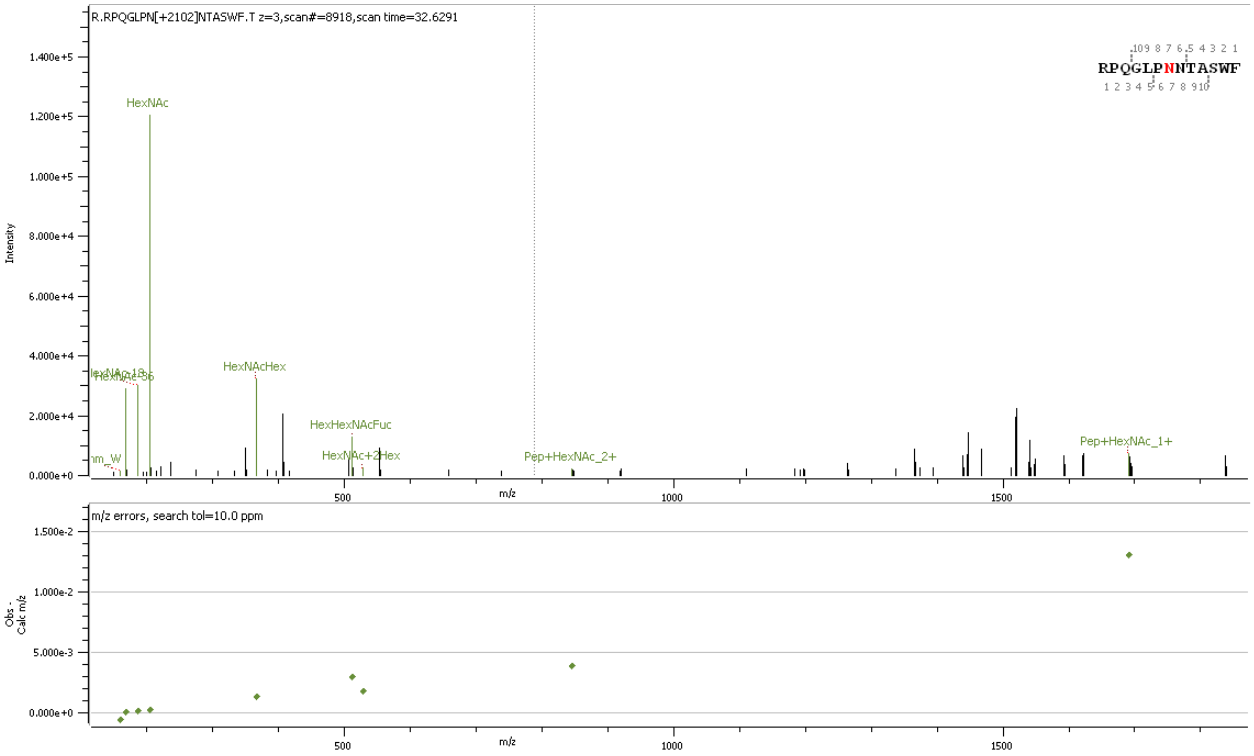


**Supp. Figure S18** HCD MS2 spectrum of N-glycopeptide [41]RPQGLPNNTASWF[53] containing HexNAc(5)Hex(4)Fuc(3)

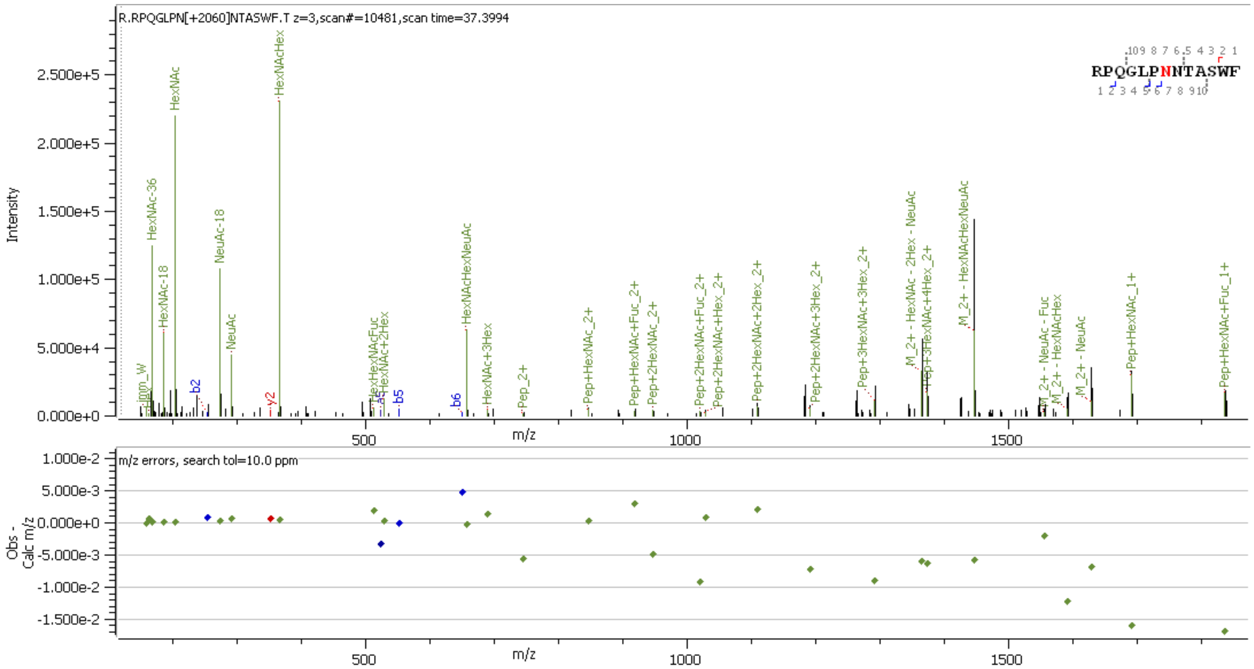


**Supp. Figure S19** HCD MS2 spectrum of N-glycopeptide [41]RPQGLPNNTASWF[53] containing HexNAc(4)Hex(5)Fuc(1)NeuAc(1)

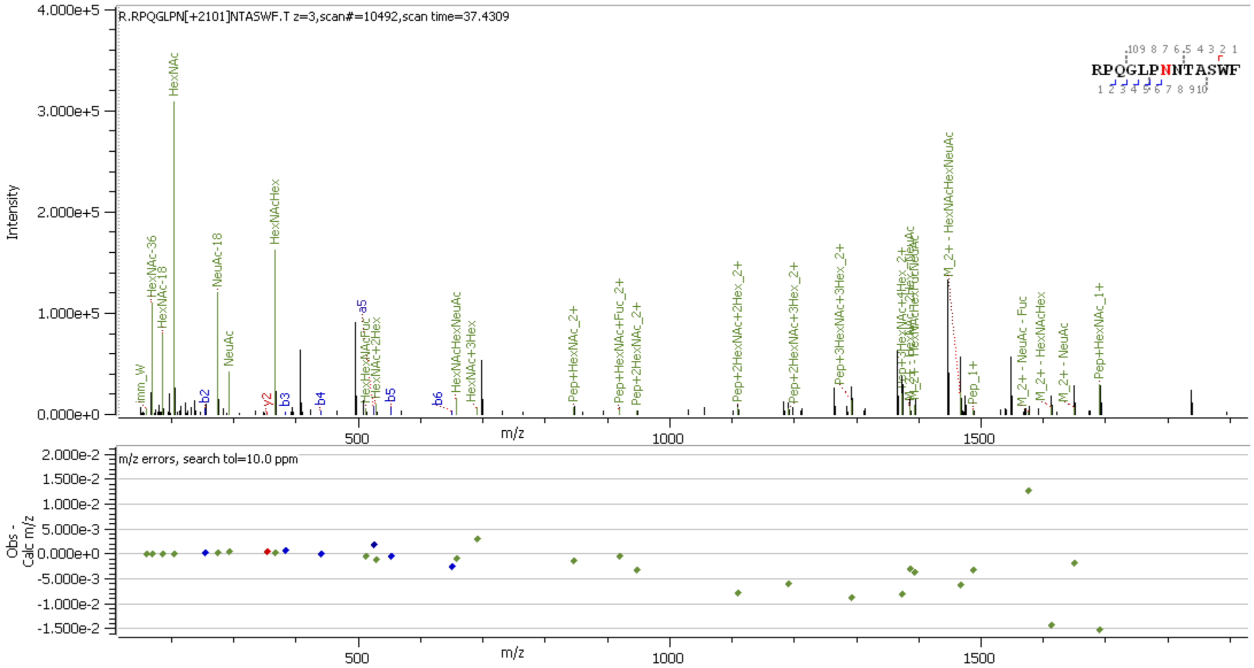


**Supp. Figure S20** HCD MS2 spectrum of N-glycopeptide [41]RPQGLPNNTASWF[53] containing HexNAc(5)Hex(4)Fuc(1)NeuAc(1)

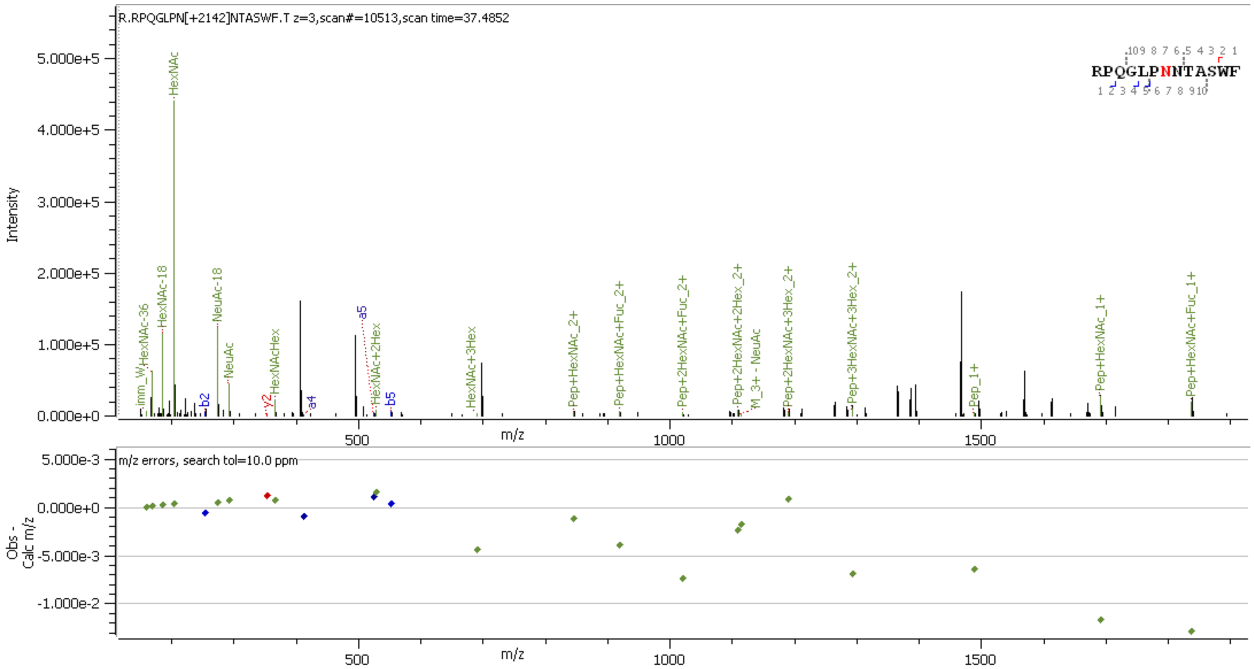


**Supp. Figure S21** HCD MS2 spectrum of N-glycopeptide [41]RPQGLPNNTASWF[53] containing HexNAc(6)Hex(3)Fuc(1)NeuAc(1)

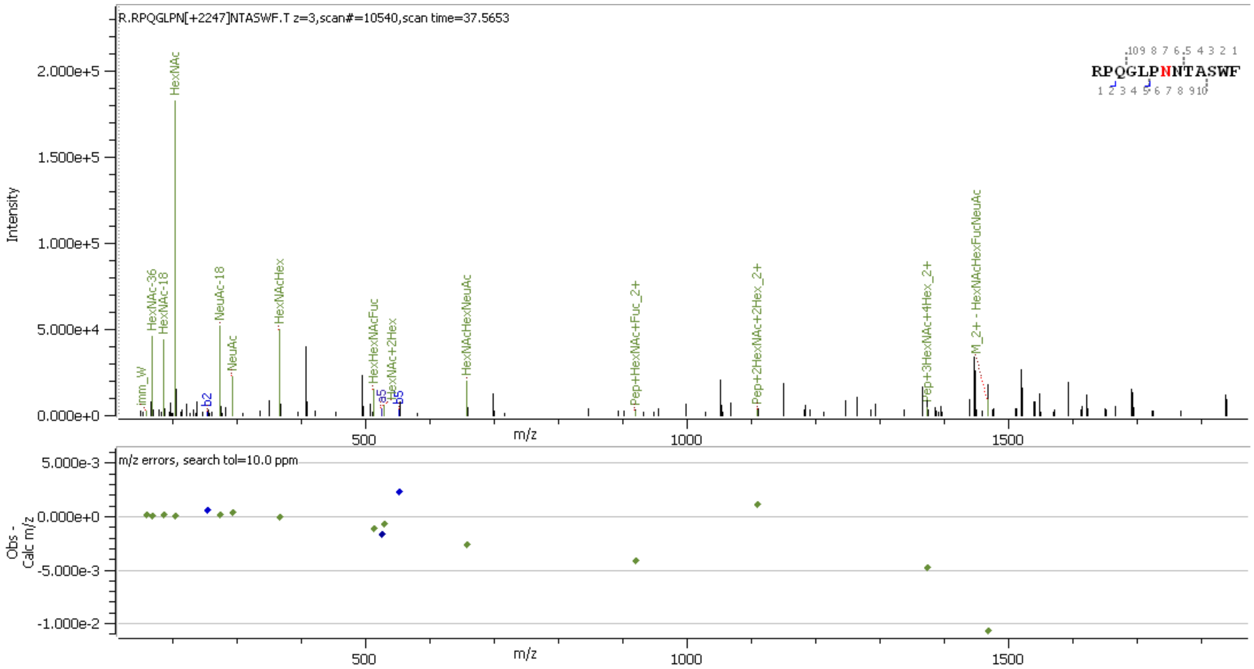


**Supp. Figure S22** HCD MS2 spectrum of N-glycopeptide [41]RPQGLPNNTASWF[53] containing HexNAc(5)Hex(4)Fuc(2)NeuAc(1)

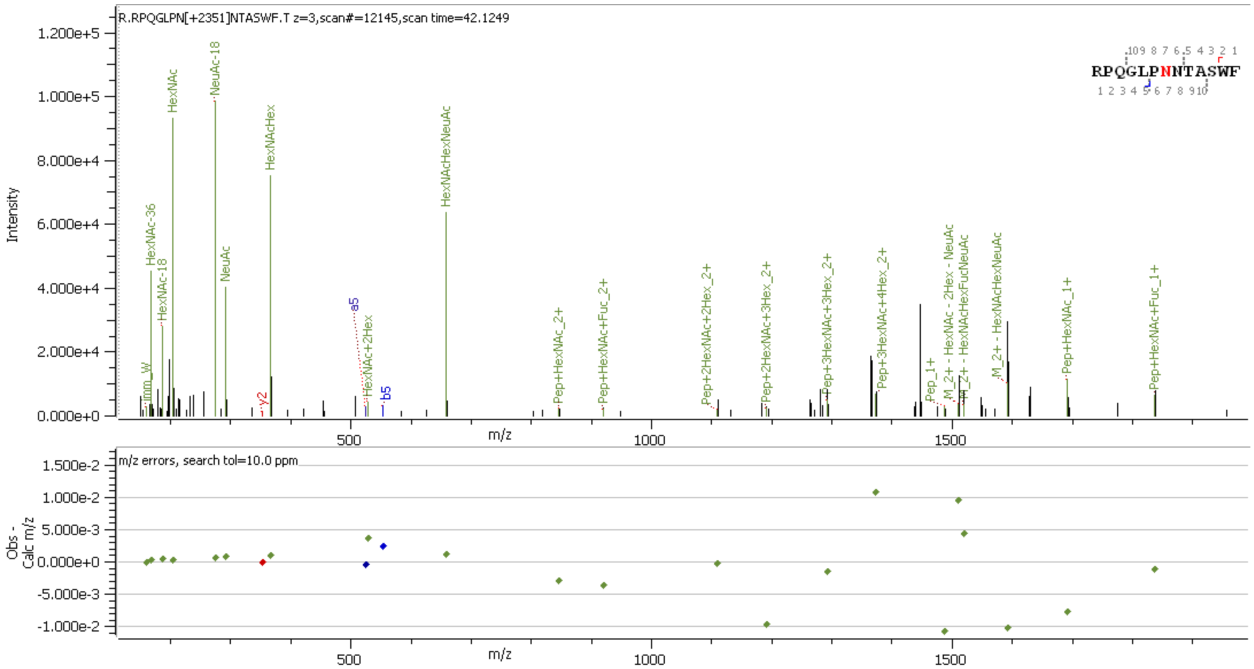


**Supp. Figure S23** HCD MS2 spectrum of N-glycopeptide [41]RPQGLPNNTASWF[53] containing HexNAc(4)Hex(5)Fuc(1)NeuAc(2)

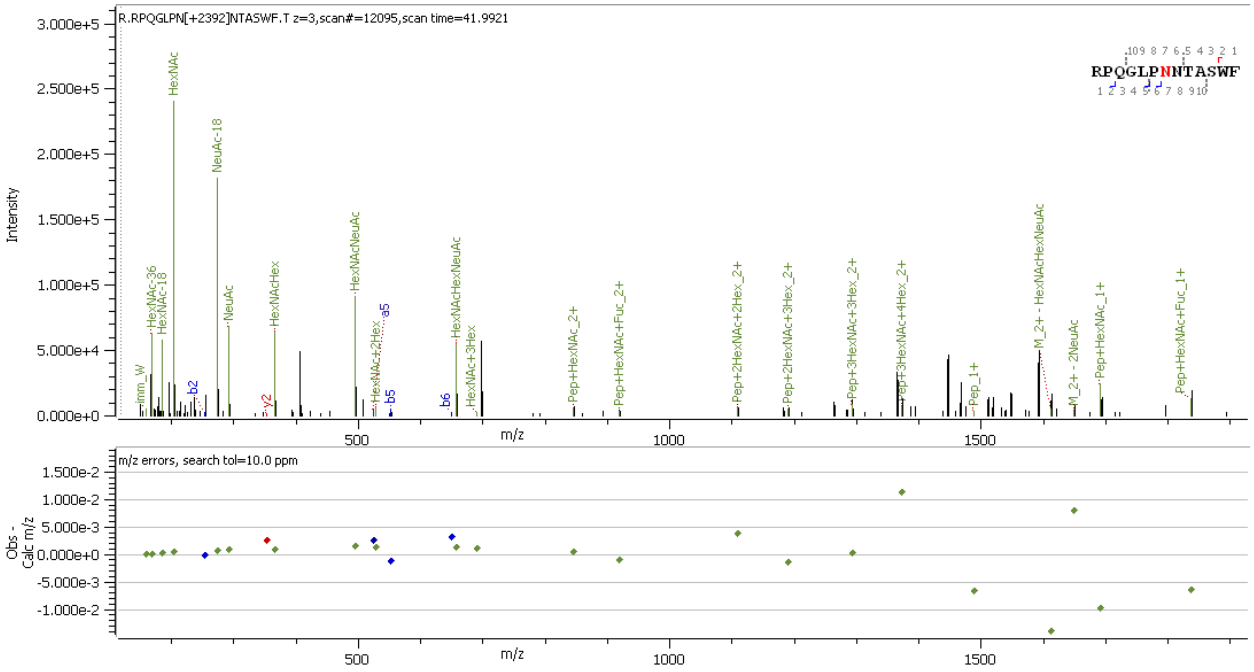


**Supp. Figure S24** HCD MS2 spectrum of N-glycopeptide [41]RPQGLPNNTASWF[53] containing HexNAc(5)Hex(4)Fuc(1)NeuAc(2)

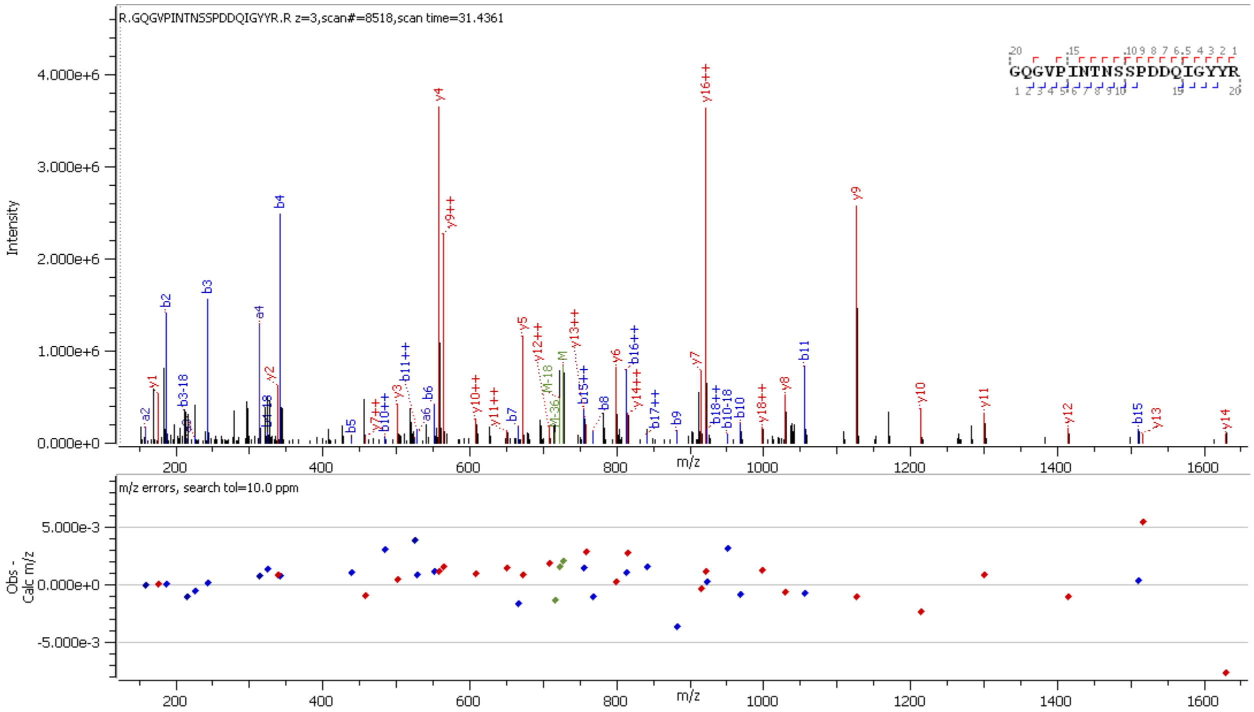


**Supp. Figure S25** HCD MS2 spectrum of unglycosylated peptide [69]GQGVPINTNSSPDDQIGYYR[88]

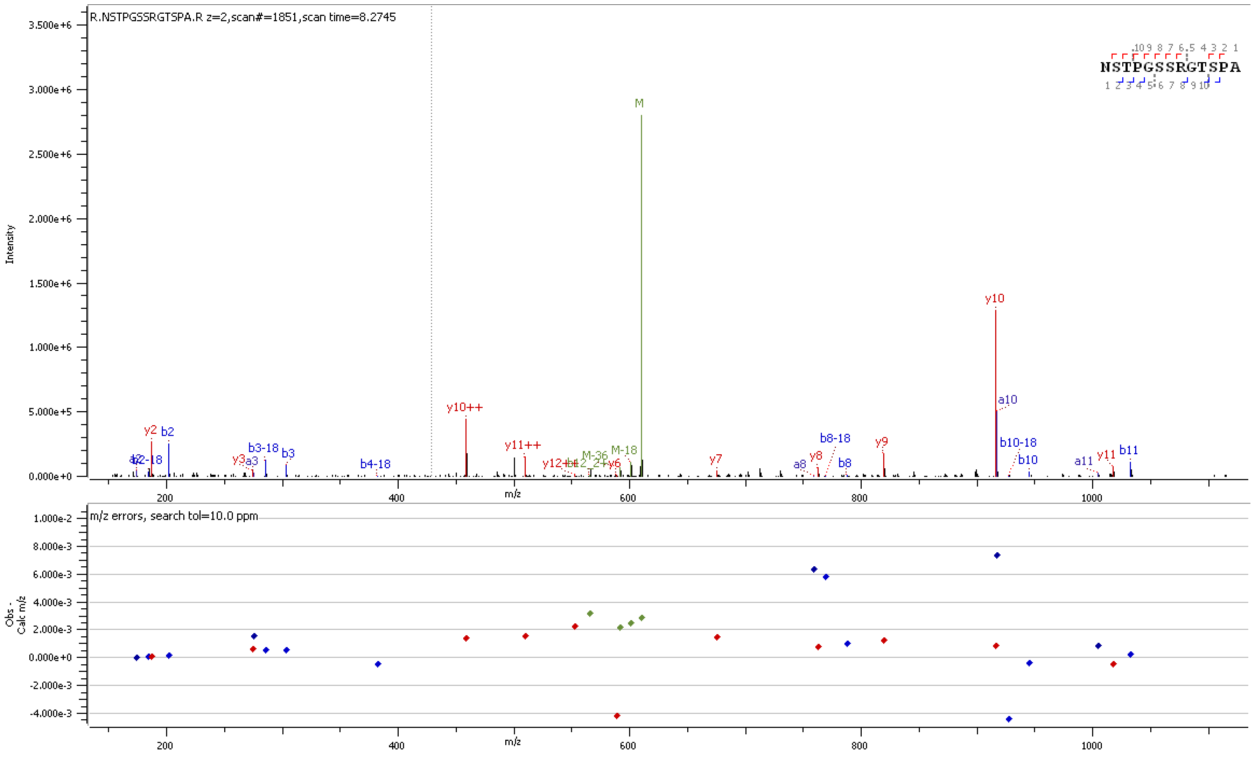


**Supp. Figure S26** HCD MS2 spectrum of peptide 196NSTPGSSRGTSPA208 showing absence of 18O-Asp at position 196

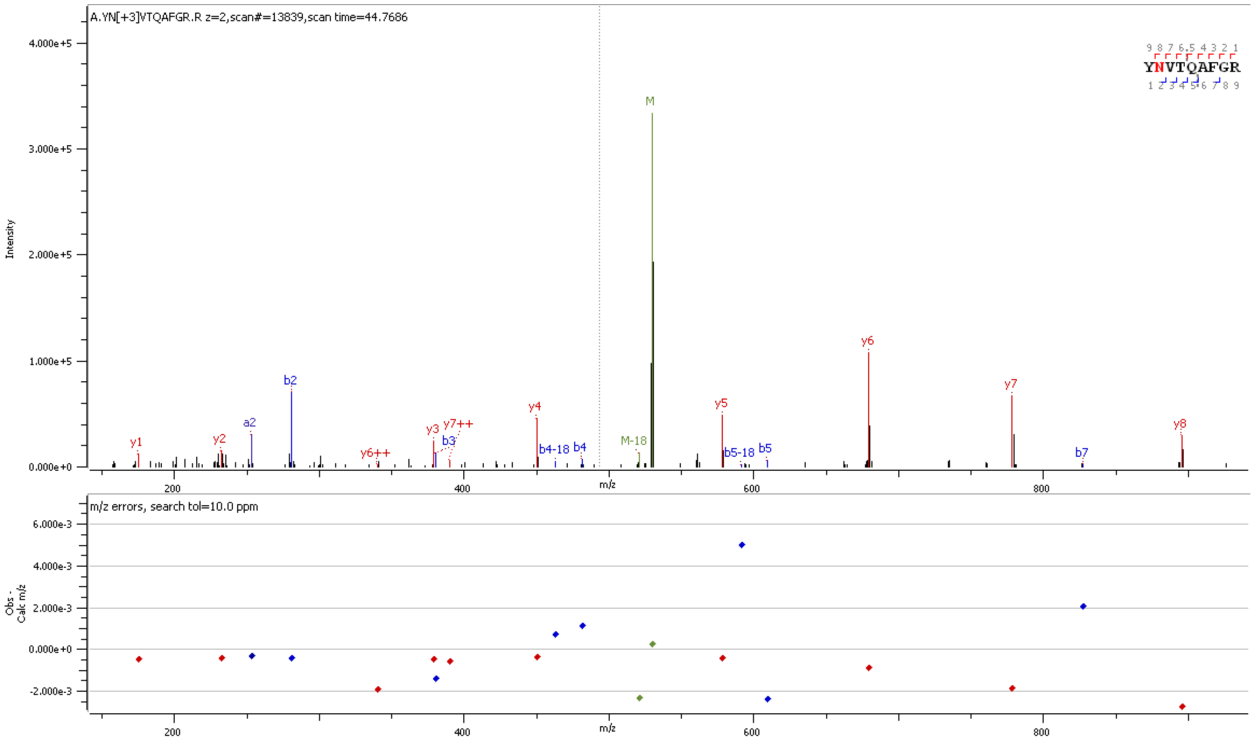


**Supp. Figure S27** HCD MS2 spectrum of peptide $^{268}$YNVTQAFGR$^{276}$ showing presence of $^{18}$O-Asp at position 269

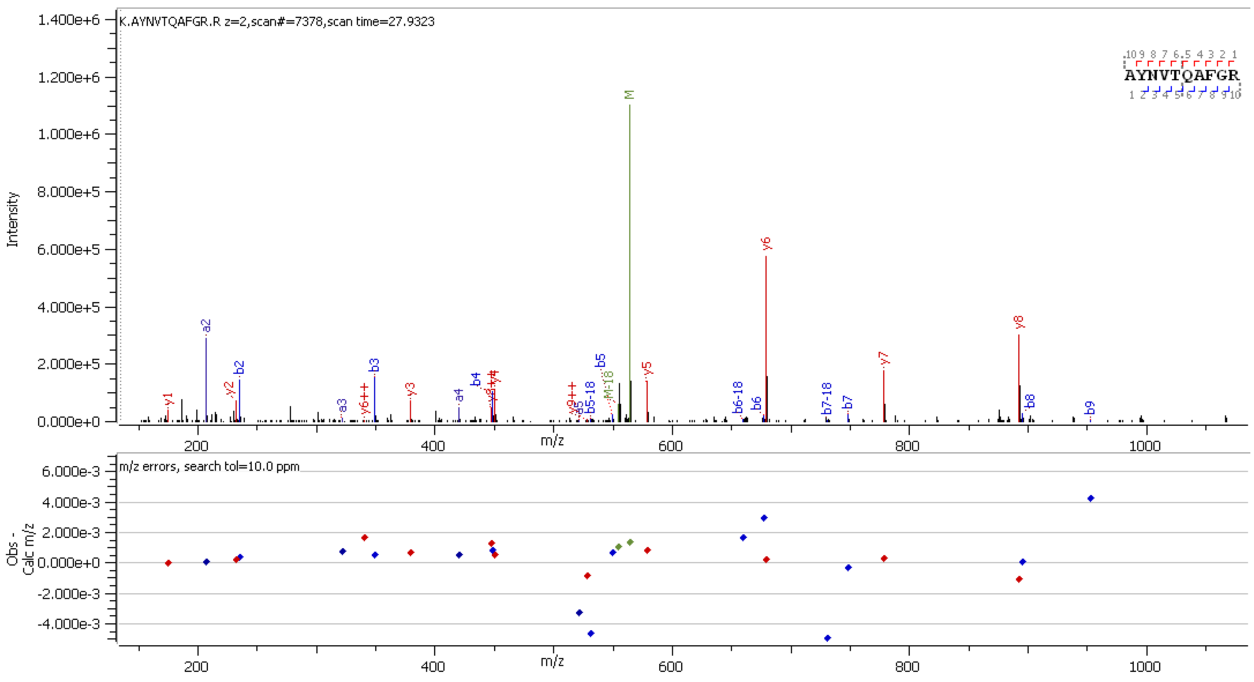


**Supp. Figure S28** HCD MS2 spectrum of unglycosylated peptide $^{267}$AYNVTQAFGR$^{276}$

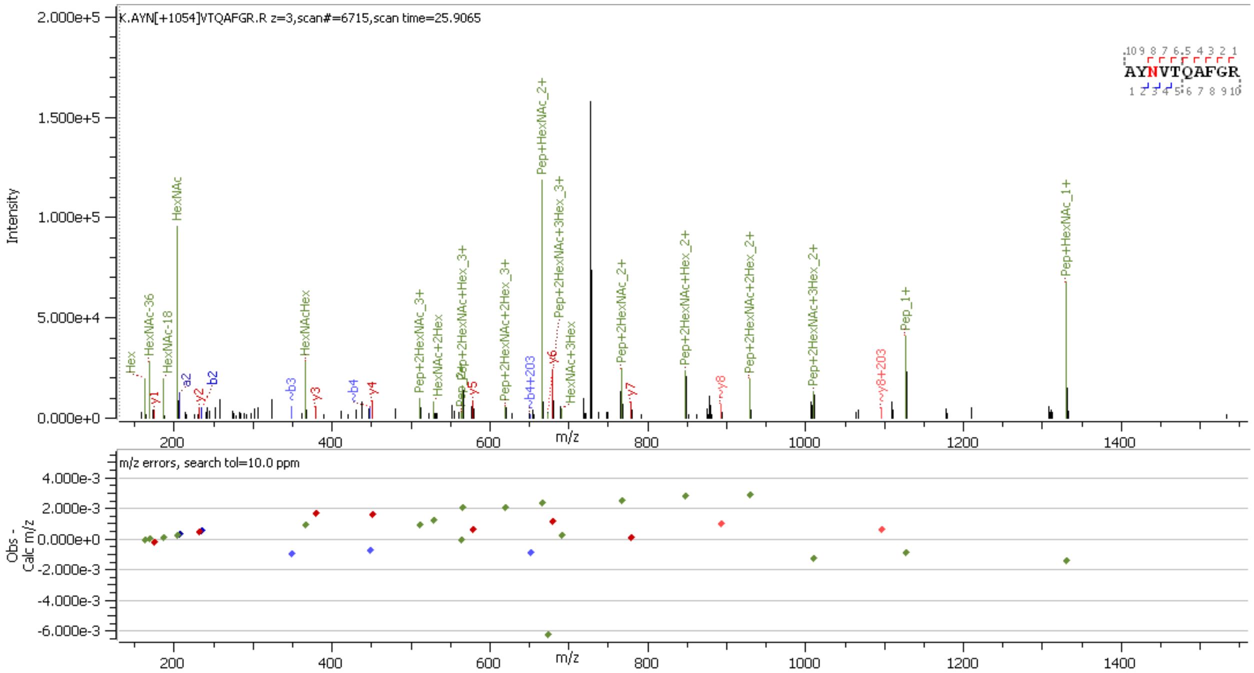


**Supp. Figure S29** HCD MS2 spectrum of N-glycopeptide [267]AYNVTQAFGR[276] containing HexNAc(2)Hex(4)

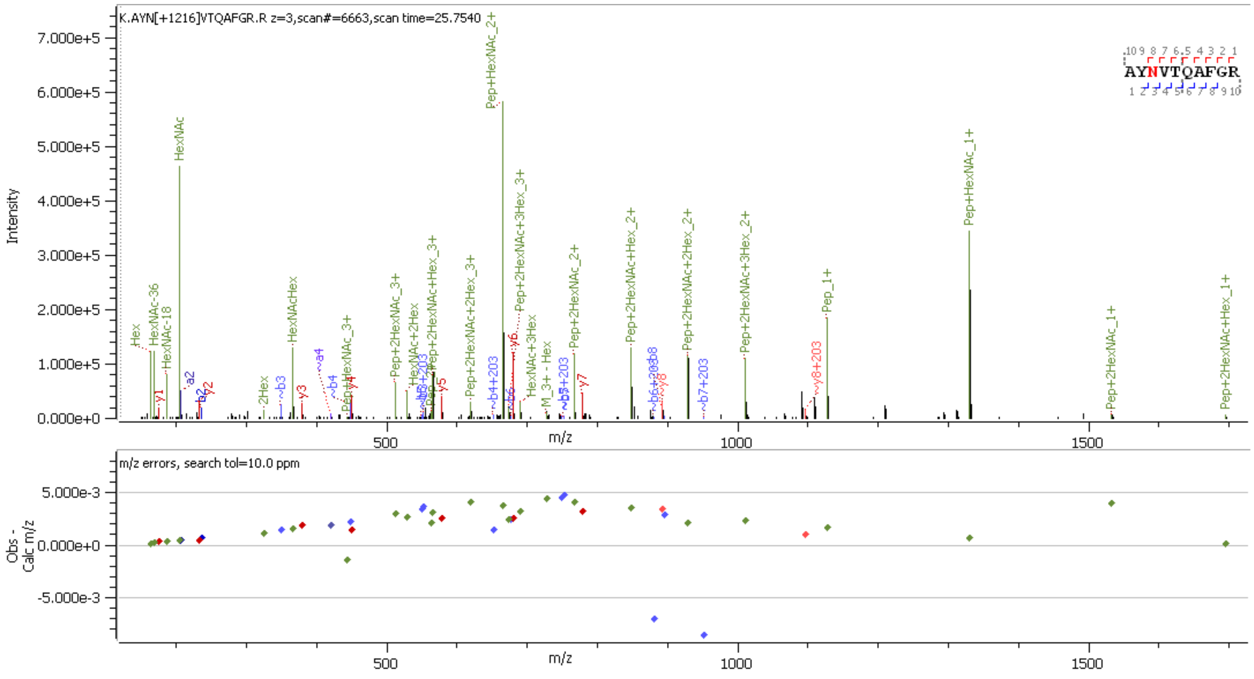


**Supp. Figure S30** HCD MS2 spectrum of N-glycopeptide [267]AYNVTQAFGR[276] containing HexNAc(2)Hex(5)

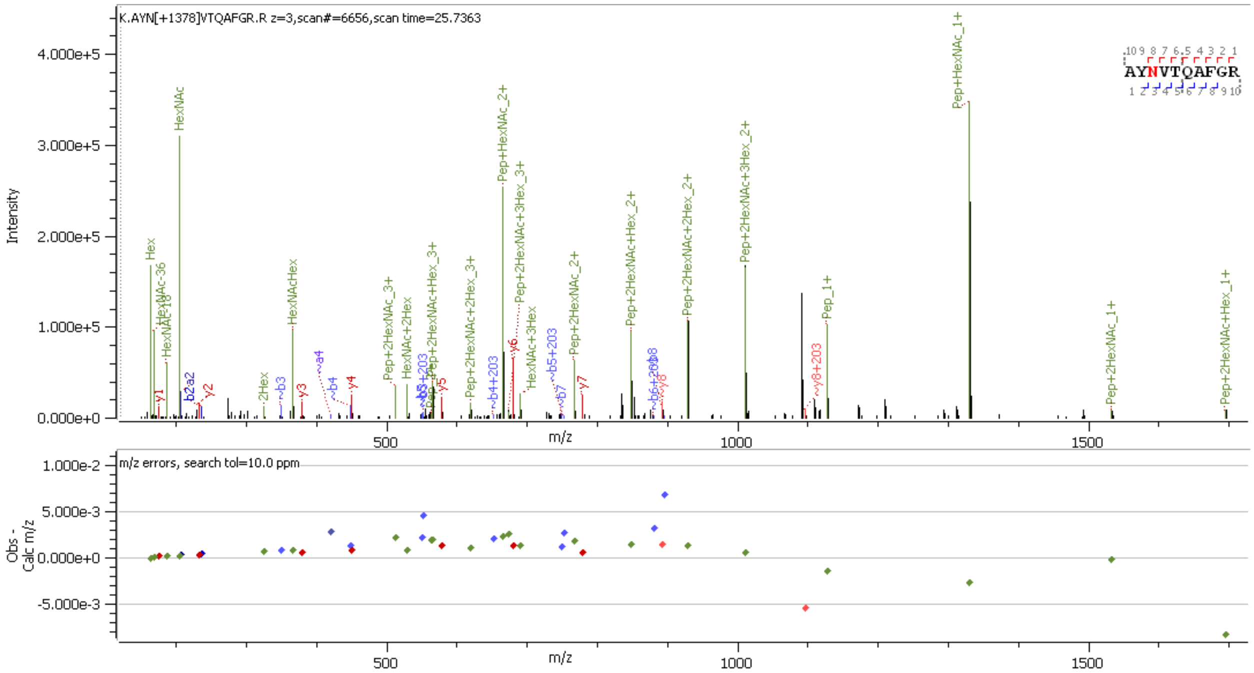


**Supp. Figure S31** HCD MS2 spectrum of N-glycopeptide [267]AYNVTQAFGR[276] containing HexNAc(2)Hex(6)

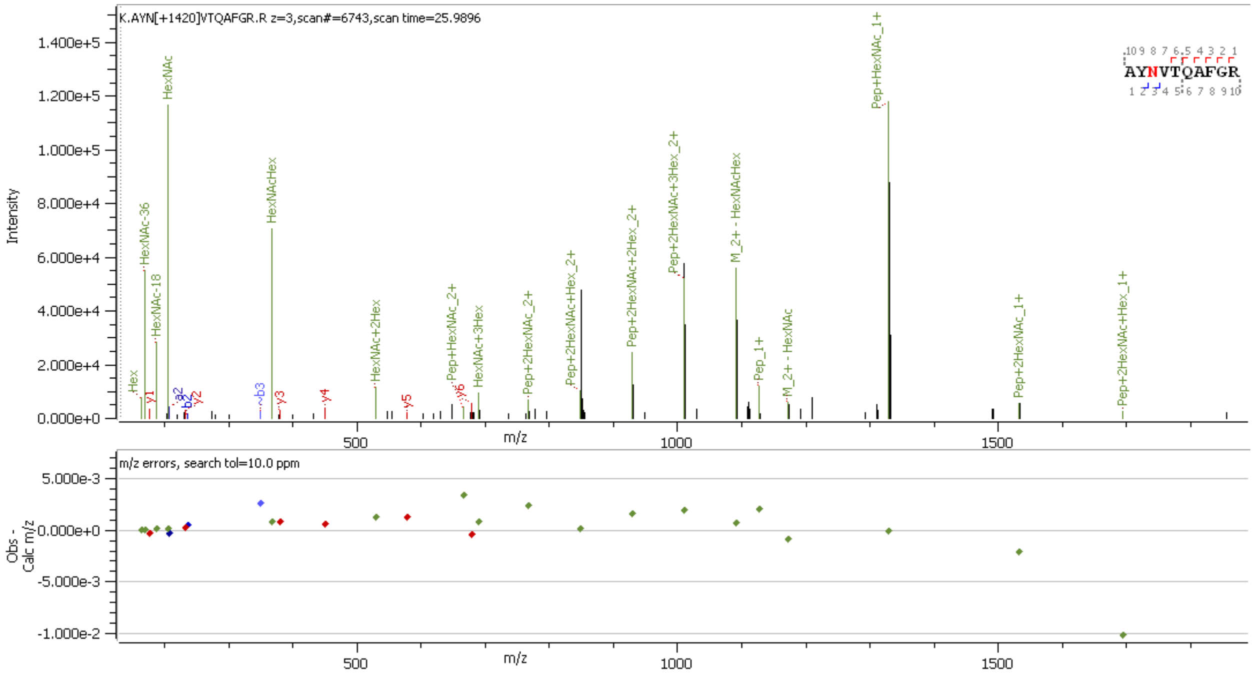


**Supp. Figure S32** HCD MS2 spectrum of N-glycopeptide [267]AYNVTQAFGR[276] containing HexNAc(3)Hex(5)

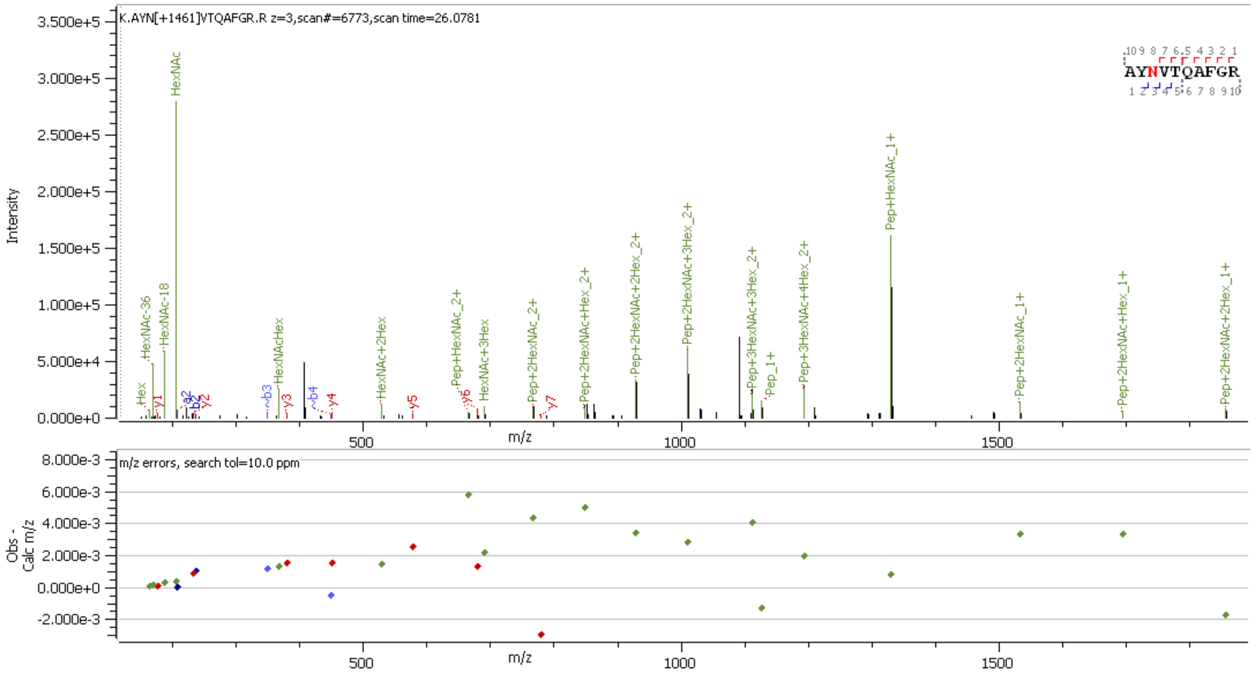


**Supp. Figure S33** HCD MS2 spectrum of N-glycopeptide [267]AYNVTQAFGR[276] containing HexNAc(4)Hex(4)

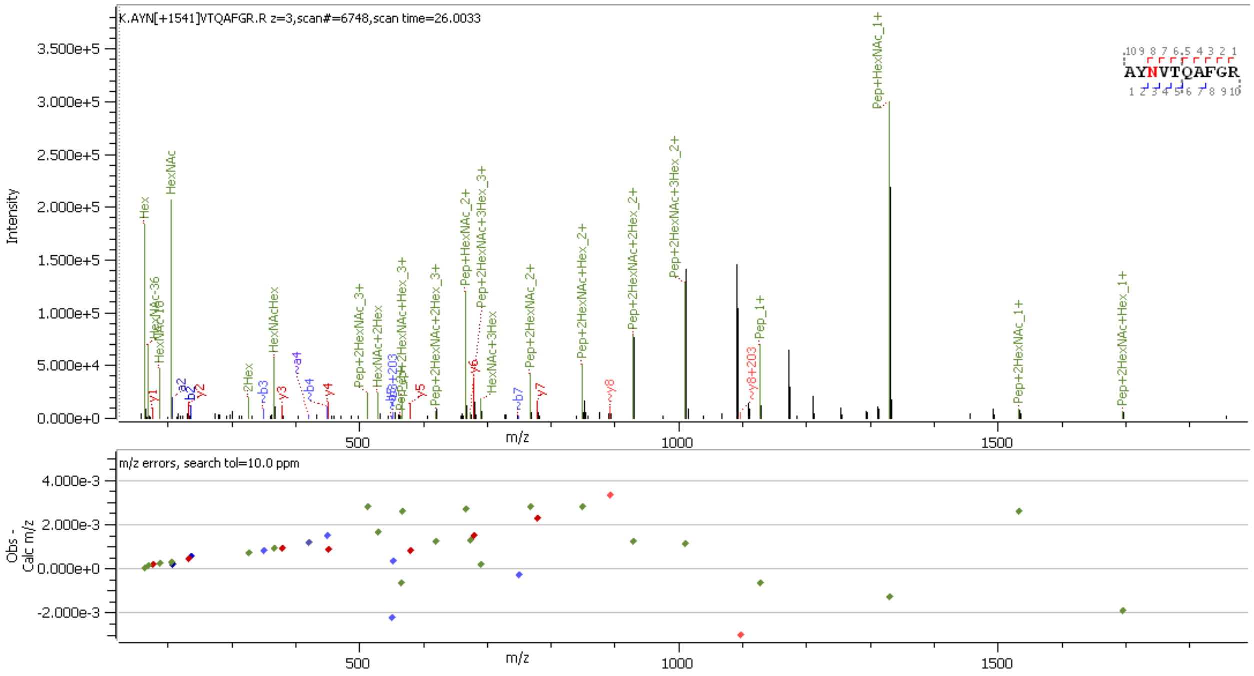


**Supp. Figure S34** HCD MS2 spectrum of N-glycopeptide [267]AYNVTQAFGR[276] containing HexNAc(2)Hex(7)

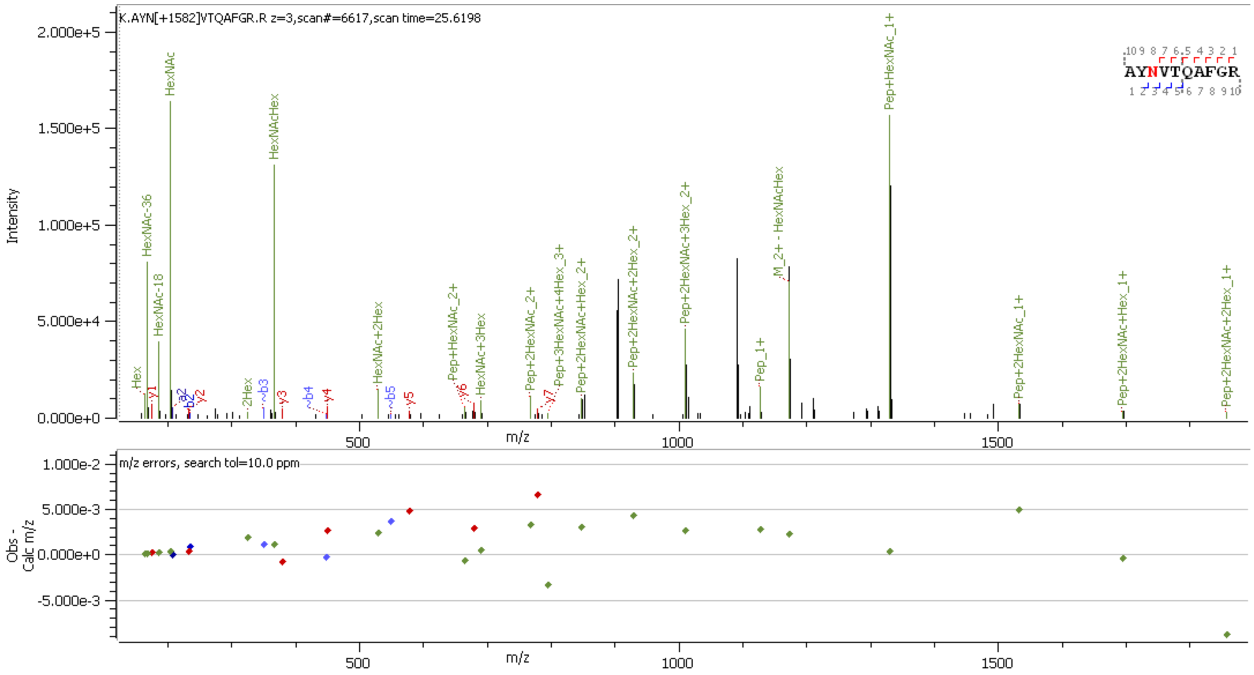


**Supp. Figure S35** HCD MS2 spectrum of N-glycopeptide [267]AYNVTQAFGR[276] containing HexNAc(3)Hex(6)

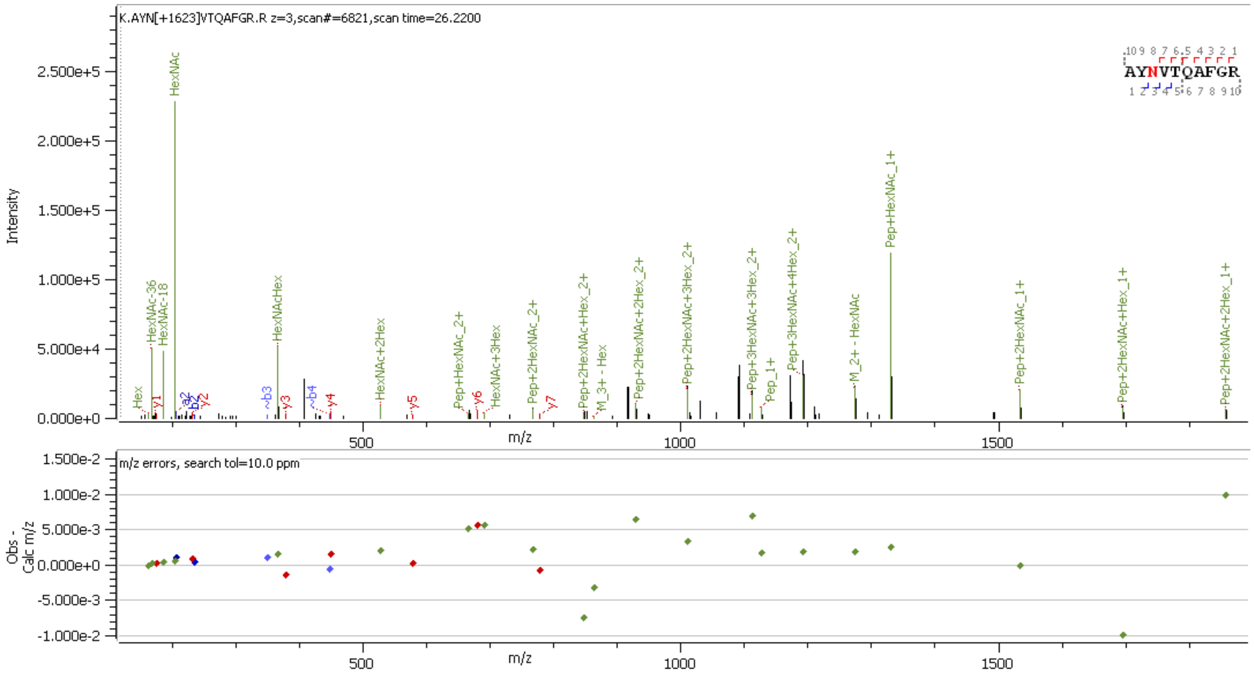


**Supp. Figure S36** HCD MS2 spectrum of N-glycopeptide [267]AYNVTQAFGR[276] containing HexNAc(4)Hex(5)

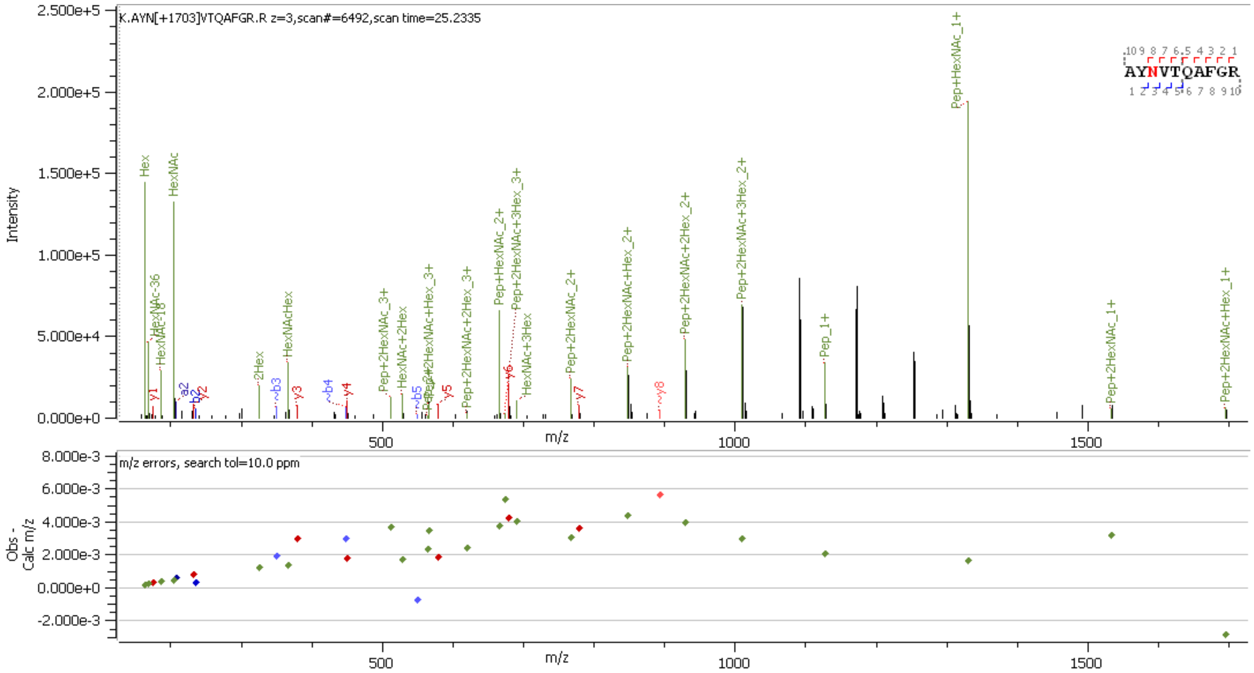


**Supp. Figure S37** HCD MS2 spectrum of N-glycopeptide [267]AYNVTQAFGR[276] containing HexNAc(2)Hex(8)

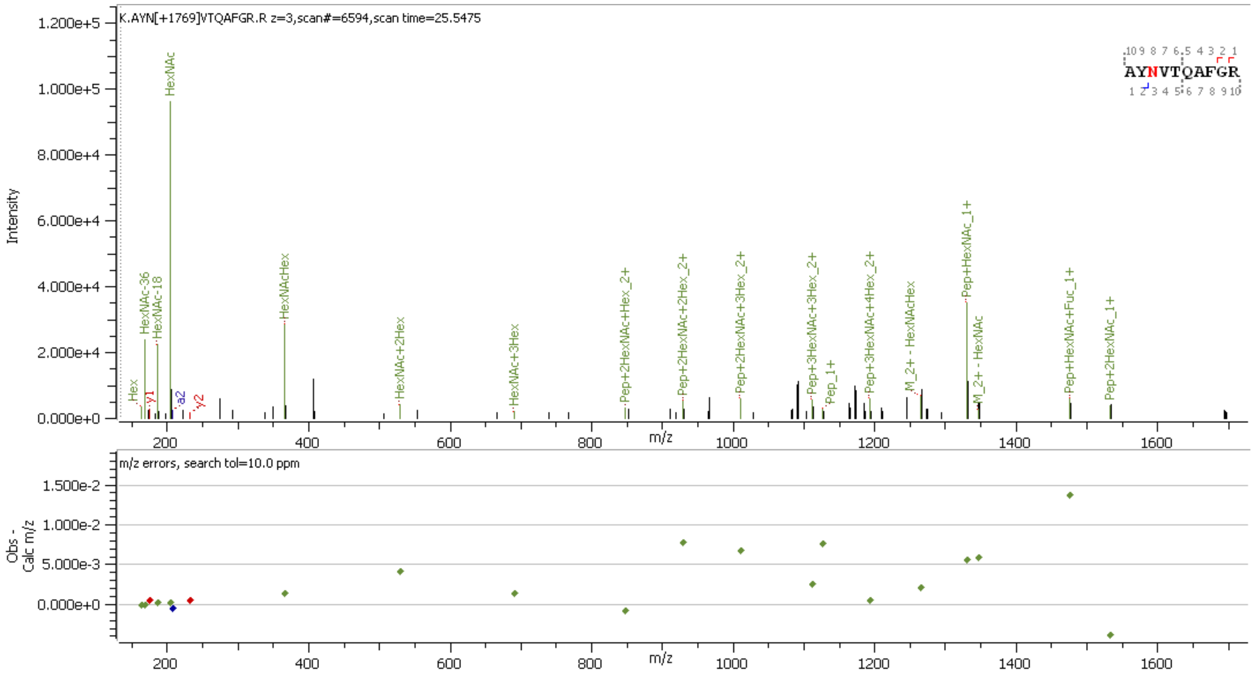


**Supp. Figure S38** HCD MS2 spectrum of N-glycopeptide [267]AYNVTQAFGR[276] containing HexNAc(4)Hex(5)Fuc(1)

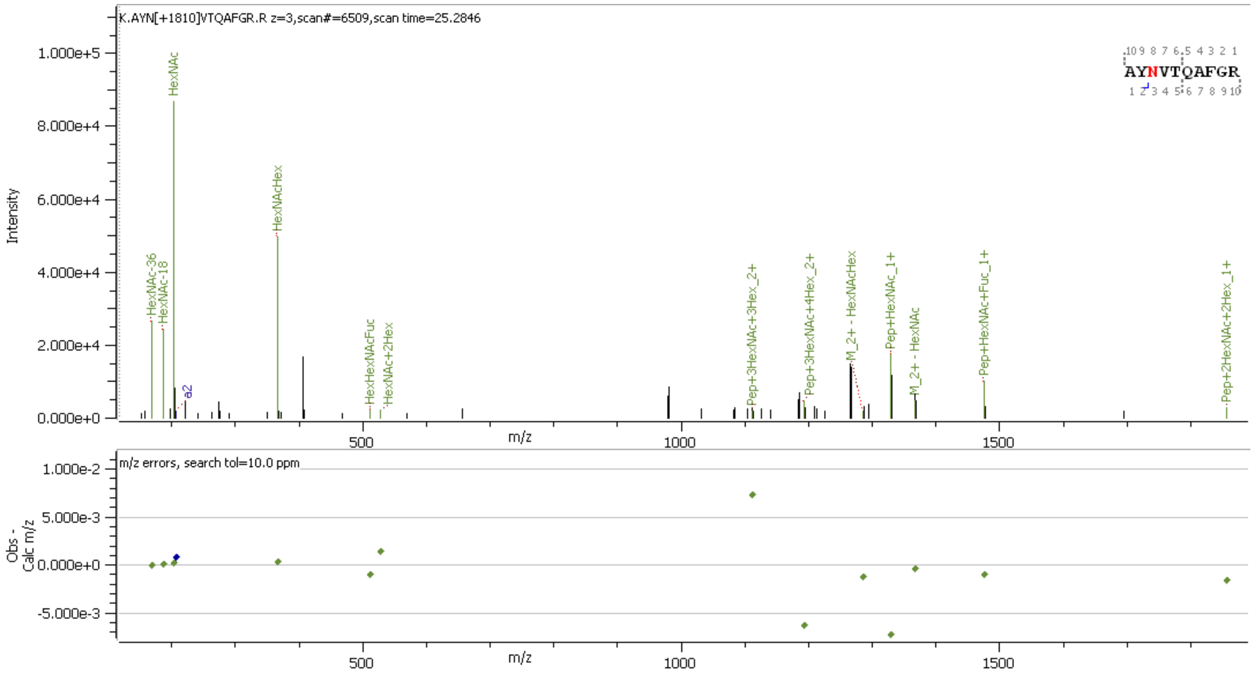


**Supp. Figure S39** HCD MS2 spectrum of N-glycopeptide [267]AYNVTQAFGR[276] containing HexNAc(5)Hex(4)Fuc(1)

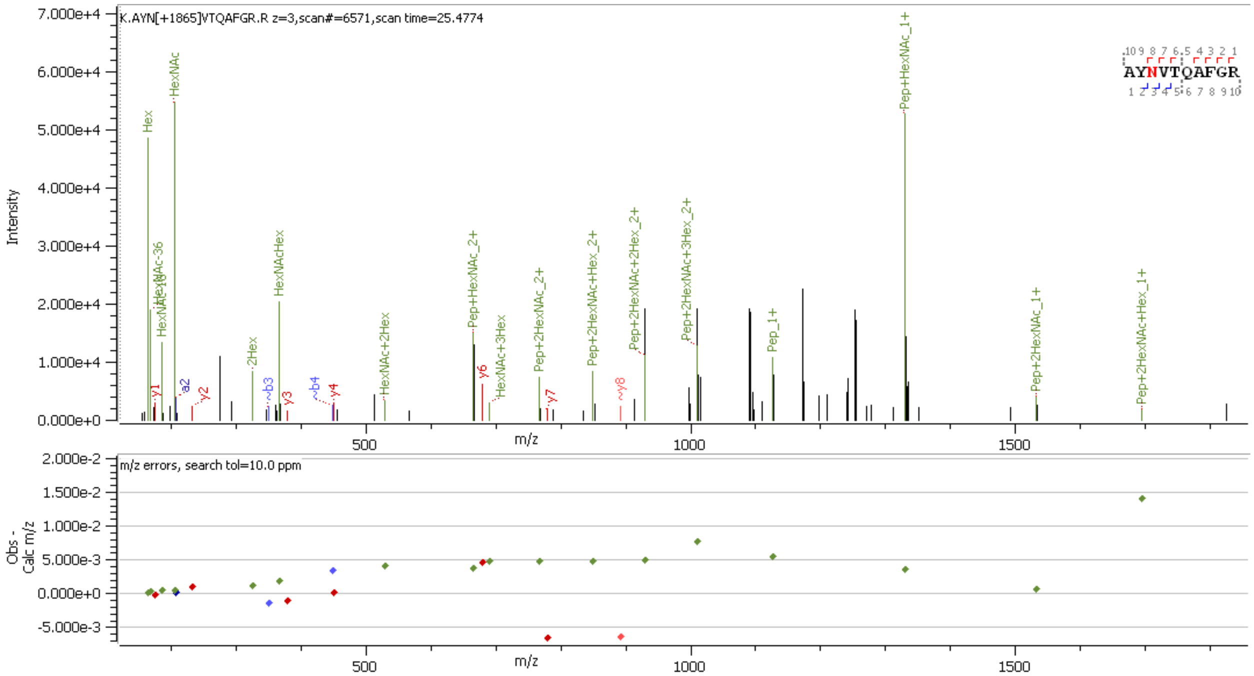


**Supp. Figure S40** HCD MS2 spectrum of N-glycopeptide [267]AYNVTQAFGR[276] containing HexNAc(2)Hex(9)

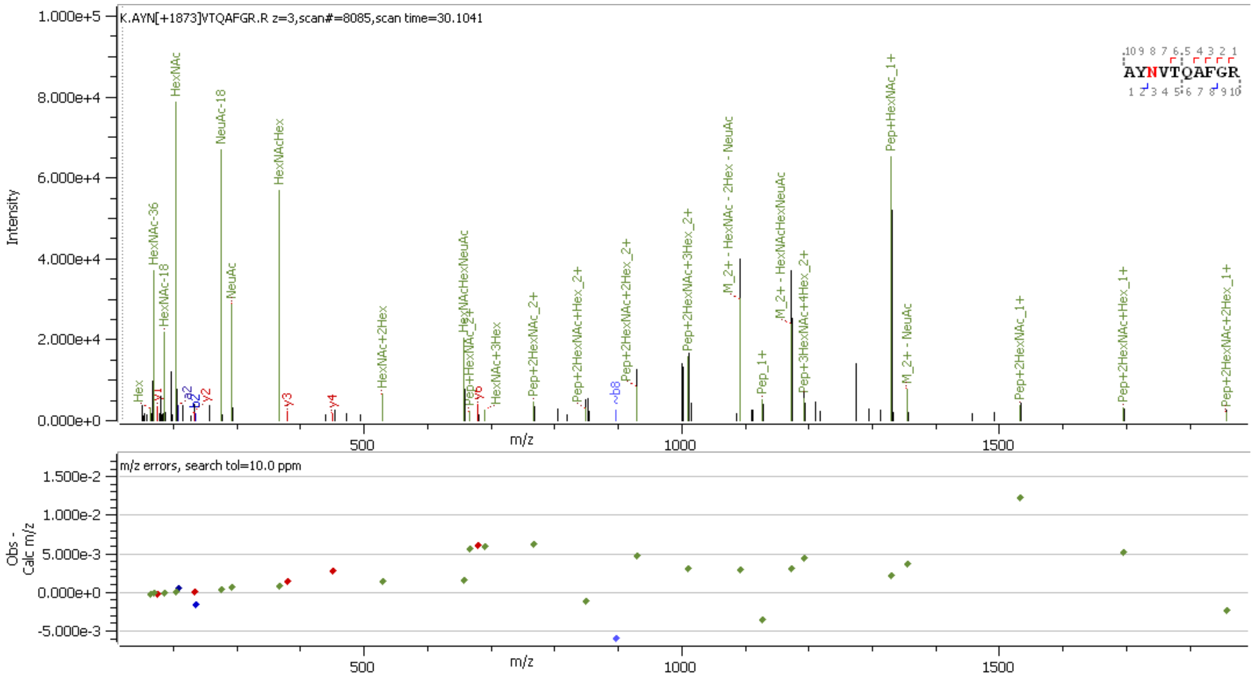


**Supp. Figure S41** HCD MS2 spectrum of N-glycopeptide [267]AYNVTQAFGR[276] containing HexNAc(3)Hex(6)NeuAc(1)

**O-glycopeptide Analysis**

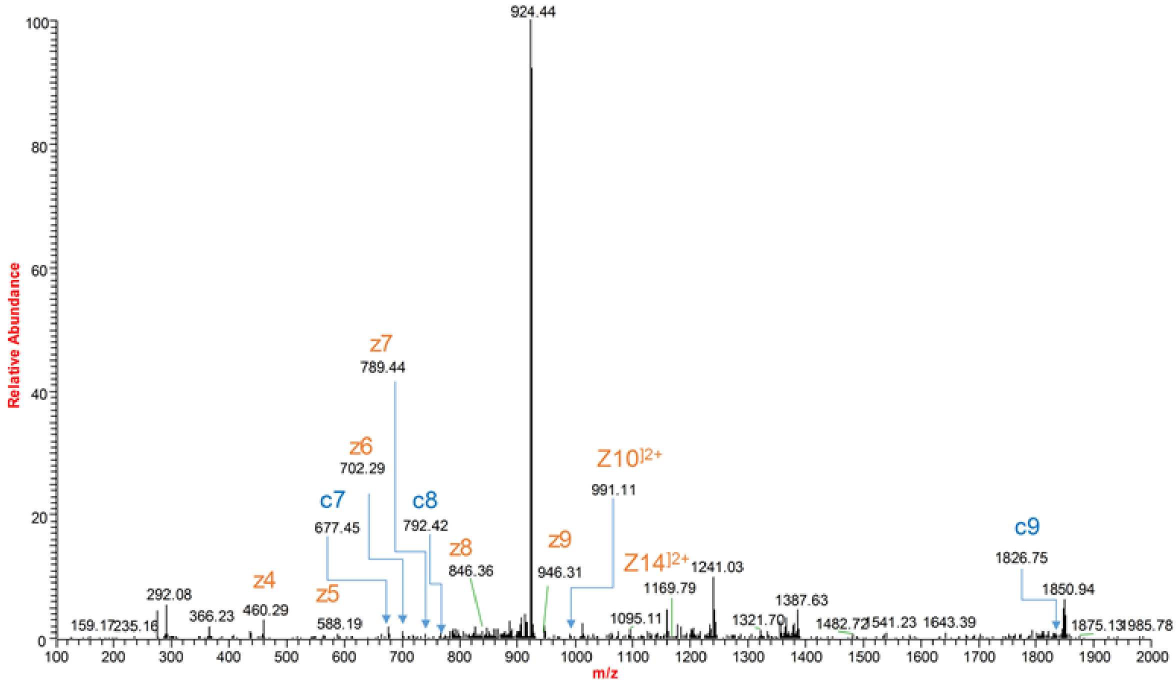


**Supp. Figure S42** ETD MS2 spectrum of O-glycopeptide [15]ITFGGPSDSTGSNQNGER[32] containing HexNAc(1)Hex(1)NeuAc(2) at site S23

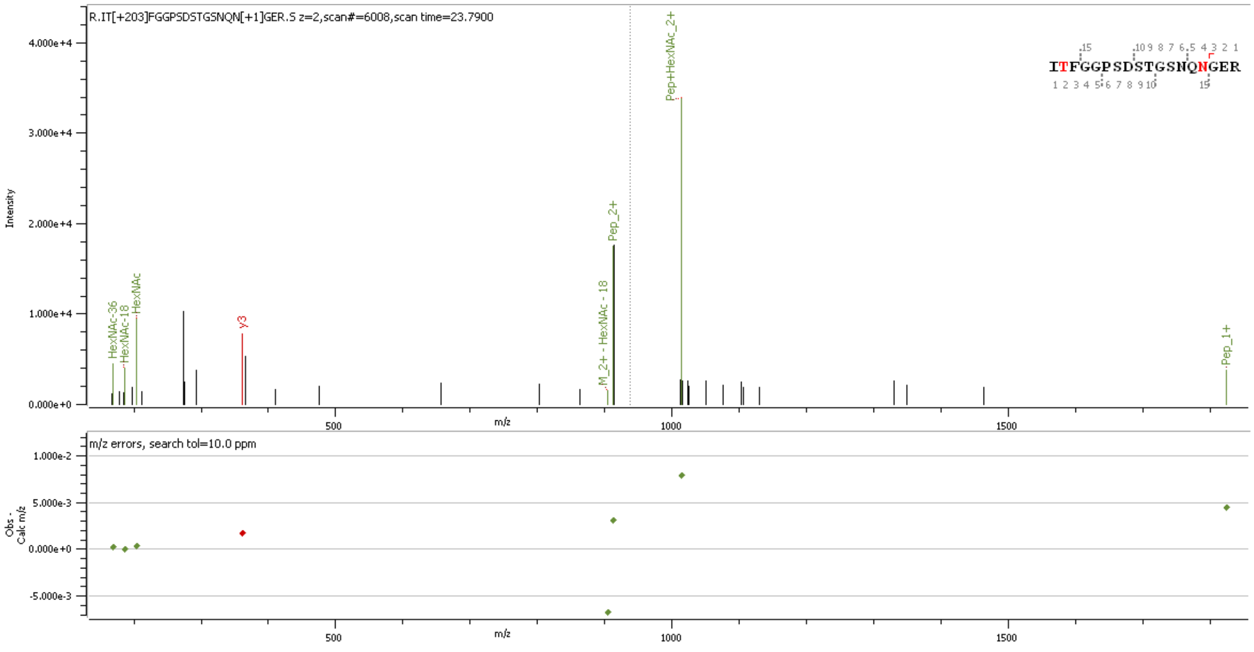


**Supp. Figure S43** HCD MS2 spectrum of O-glycopeptide [15]ITFGGPSDSTGSNQNGER[32] containing HexNAc(1) at site S23

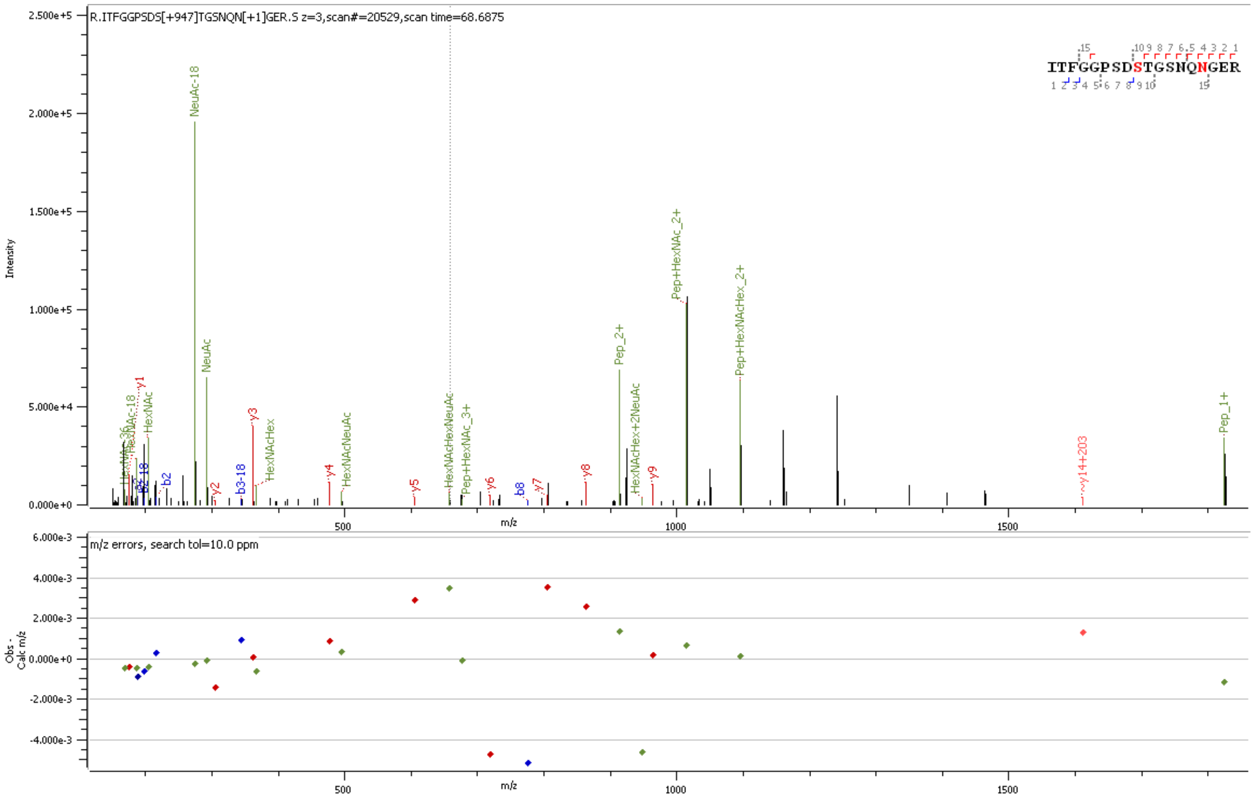

**Supp. Figure S44** HCD MS2 spectrum of O-glycopeptide [15]ITFGGPSDSTGSNQNGER[32] containing HexNAc(1)Hex(1)NeuAc(2) at site S23

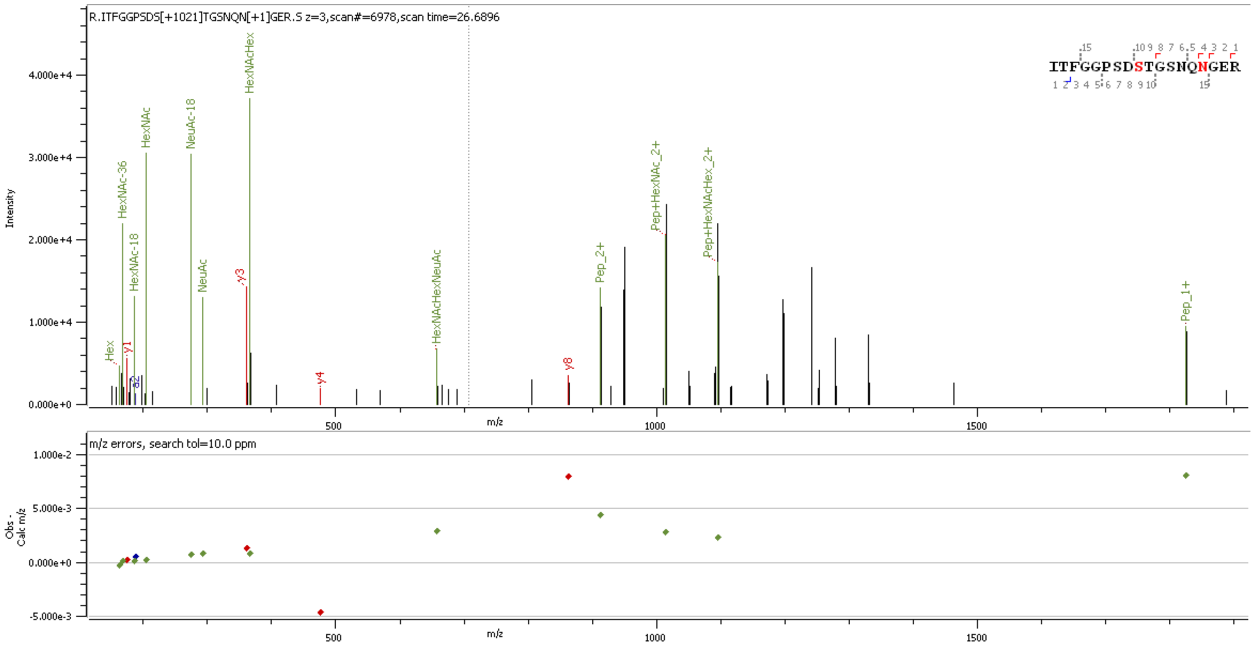

**Supp. Figure S45** HCD MS2 spectrum of O-glycopeptide [15]ITFGGPSDSTGSNQNGER[32] containing HexNAc(2)Hex(2)NeuAc(1) at site S23

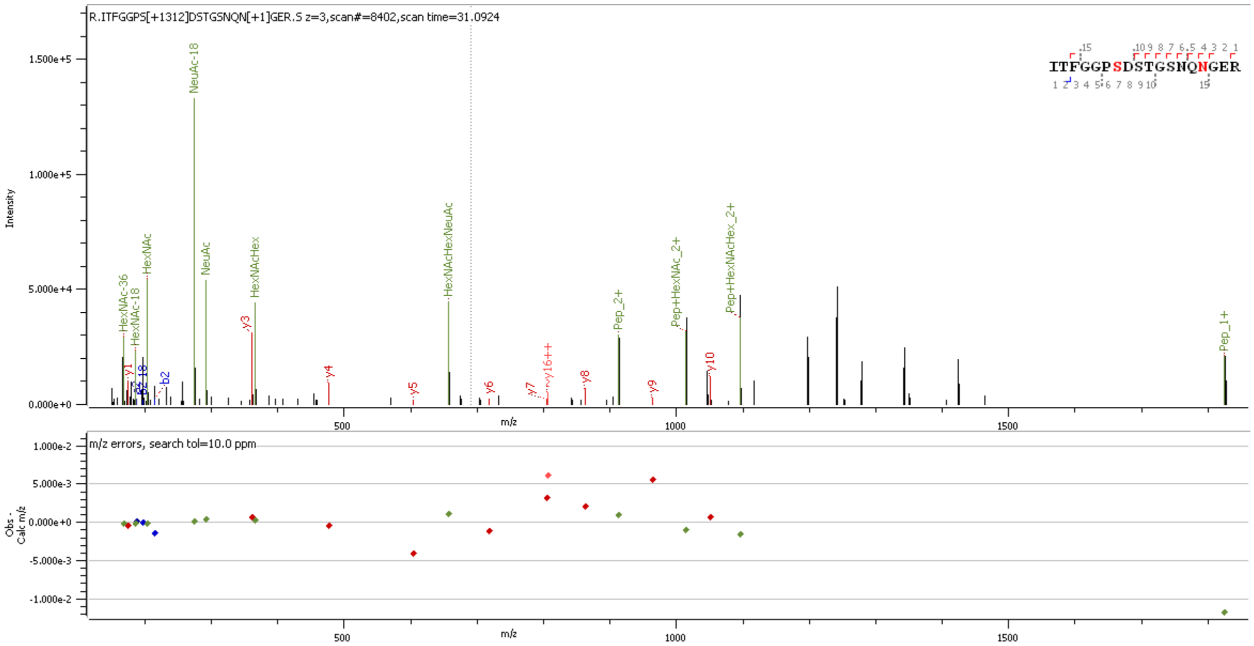


**Supp. Figure S46** HCD MS2 spectrum of O-glycopeptide [15]ITFGGPSDSTGSNQNGER[32] containing HexNAc(2)Hex(2)NeuAc(2) at site S23

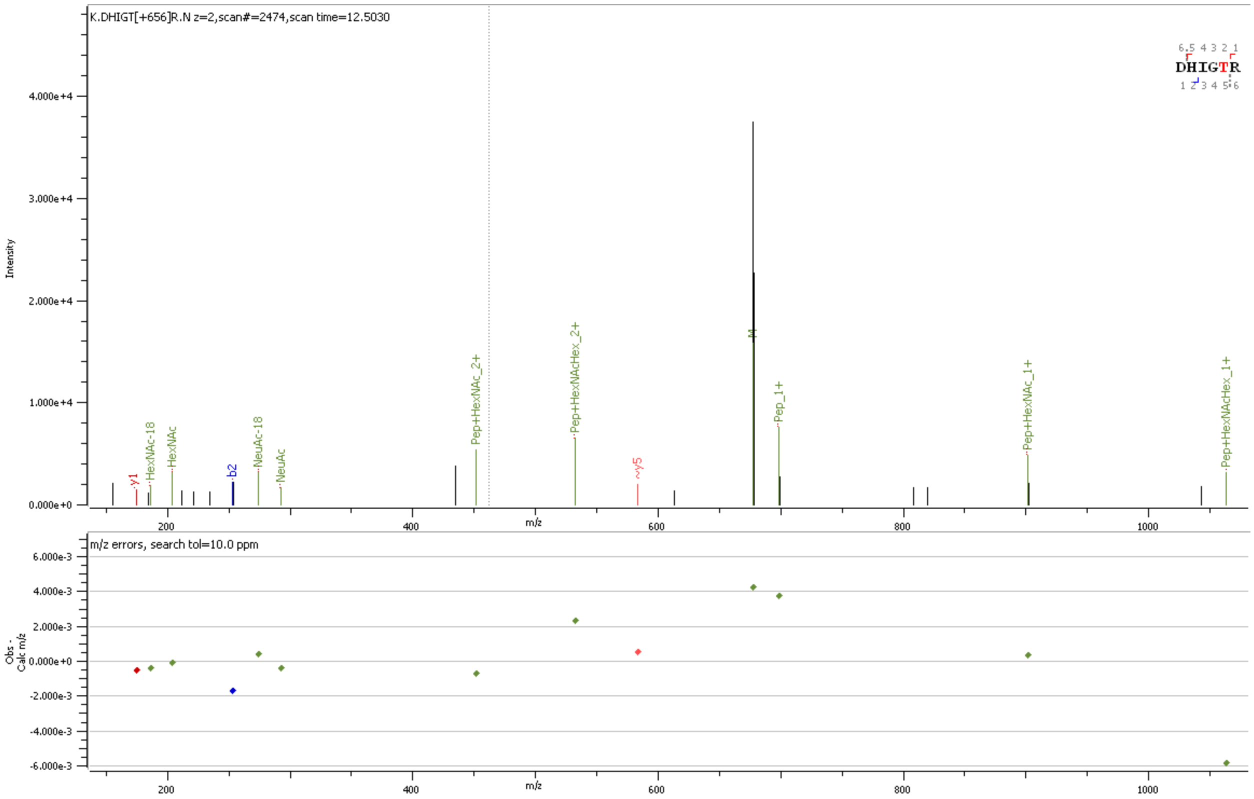


**Supp. Figure S47** HCD MS2 spectrum of O-glycopeptide [144]DHIGTR[149]containing HexNAc(1)Hex(1)NeuAc(1) at site T148

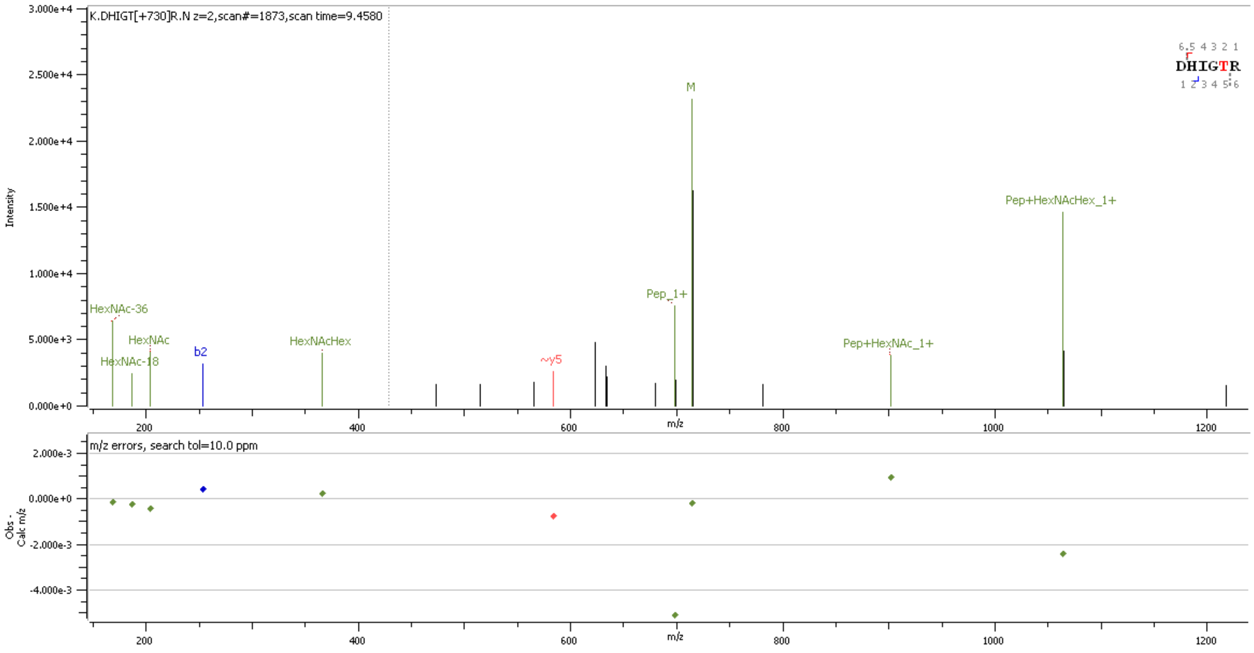


**Supp. Figure S48** HCD MS2 spectrum of O-glycopeptide [144]DHIGTR[149]containing HexNAc(2)Hex(2) at site T148

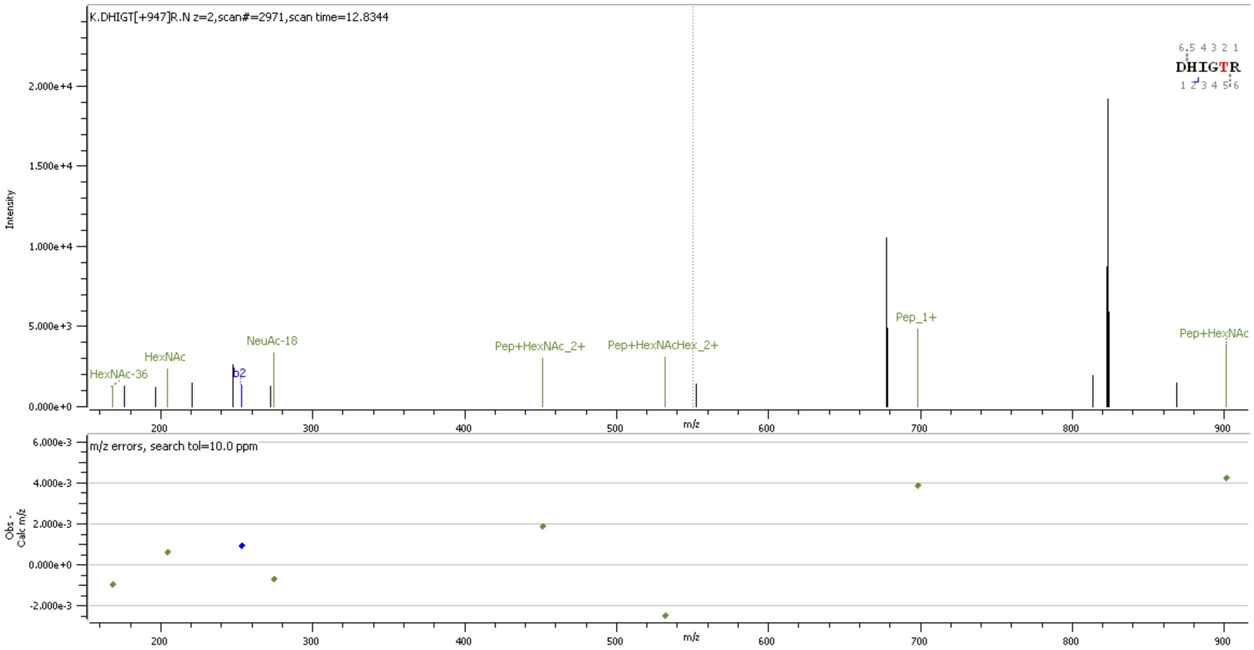


**Supp. Figure S49** HCD MS2 spectrum of O-glycopeptide [144]DHIGTR[149]containing HexNAc(1)Hex(1)NeuAc(2) at site T148

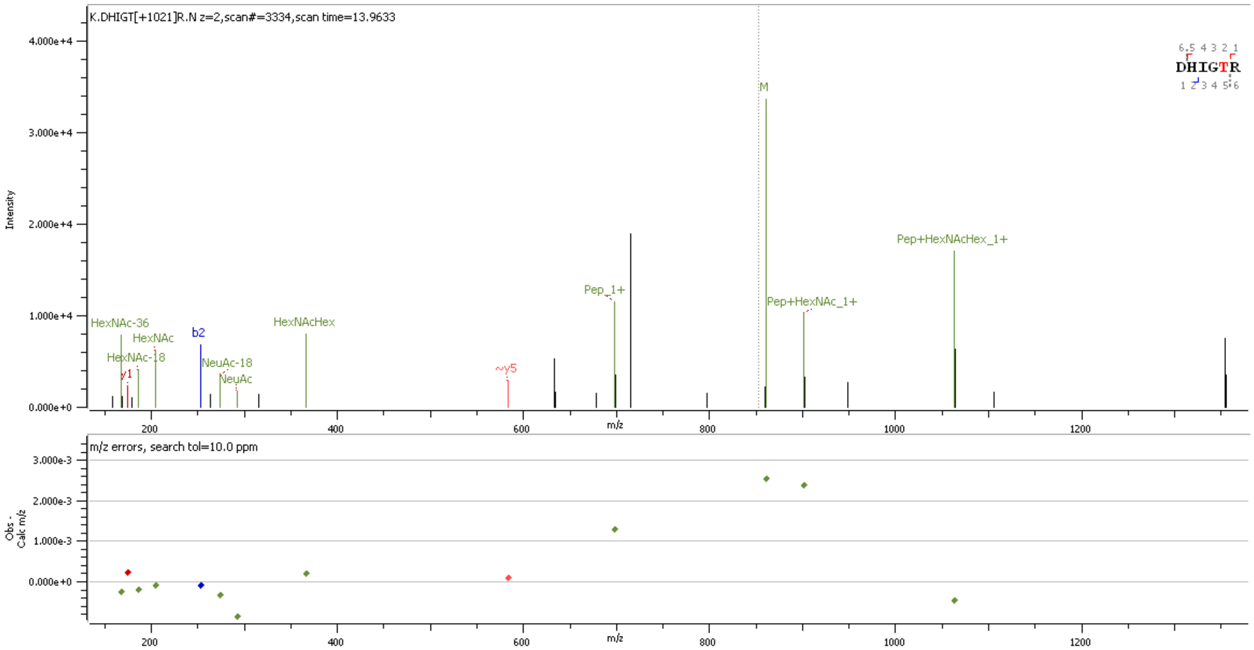


**Supp. Figure S50** HCD MS2 spectrum of O-glycopeptide [144]DHIGTR[149]containing HexNAc(2)Hex(2)NeuAc(1) at site T148

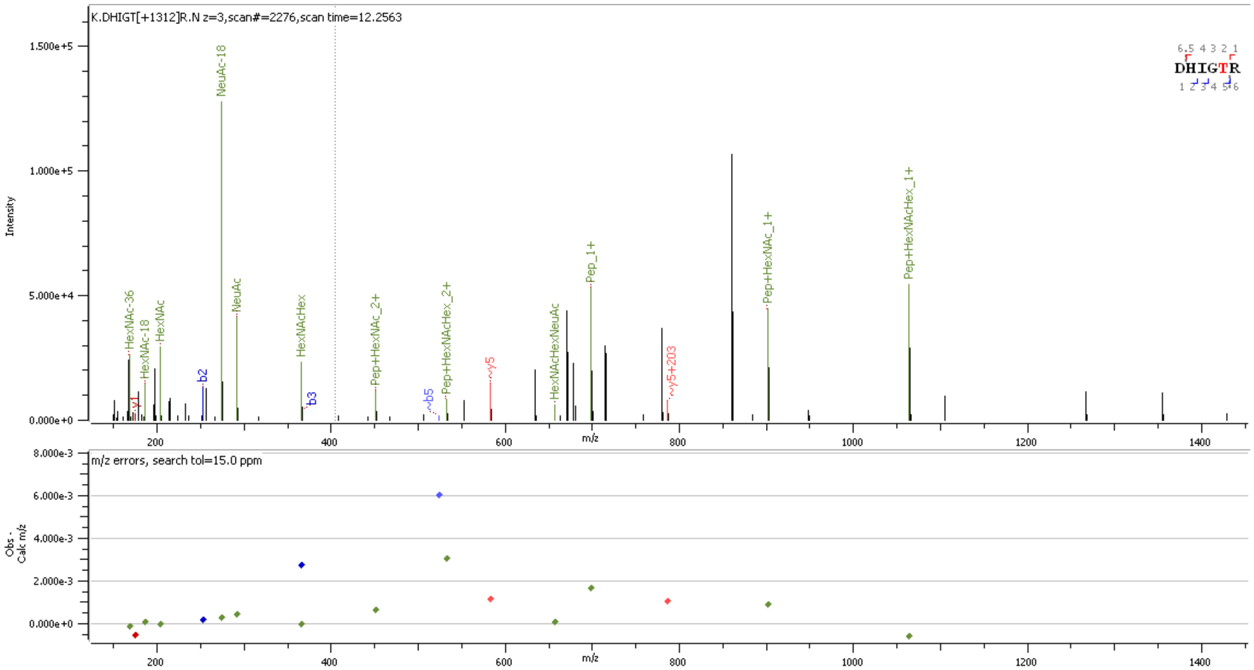


**Supp. Figure S51** HCD MS2 spectrum of O-glycopeptide [144]DHIGTR[149]containing HexNAc(2)Hex(2)NeuAc(2) at site T148

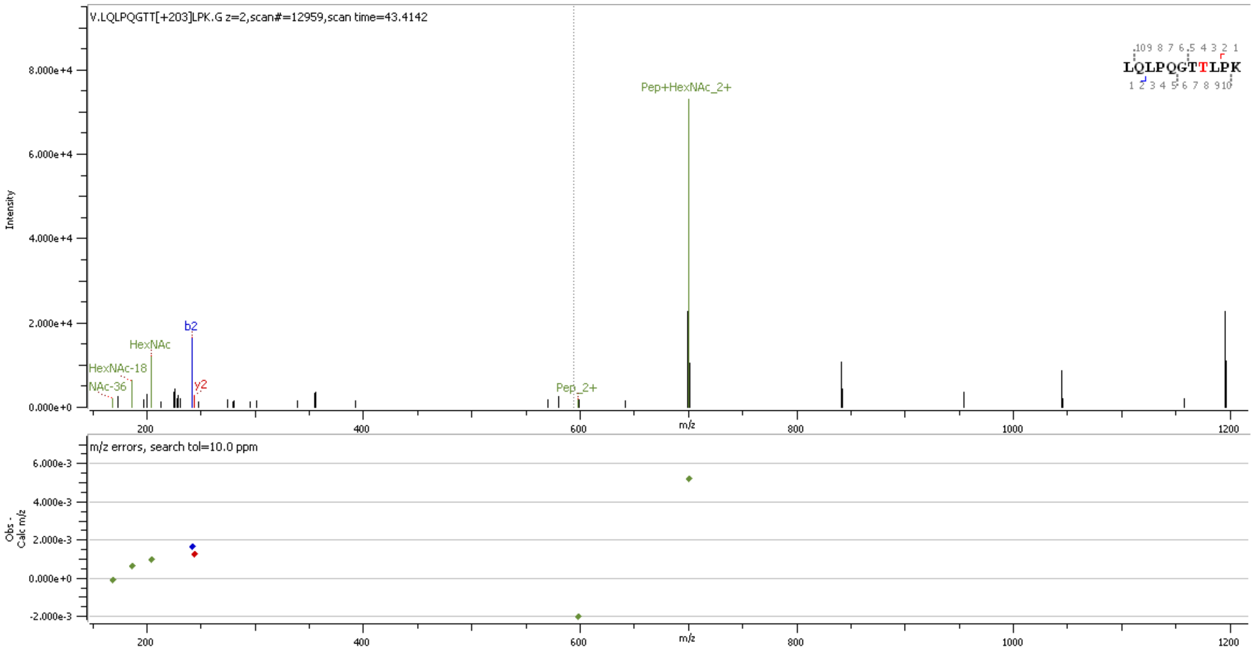


**Supp. Figure S52** HCD MS2 spectrum of O-glycopeptide [159]LQLPQGTTLPK[169]containing HexNAc(1) at site T165/T166

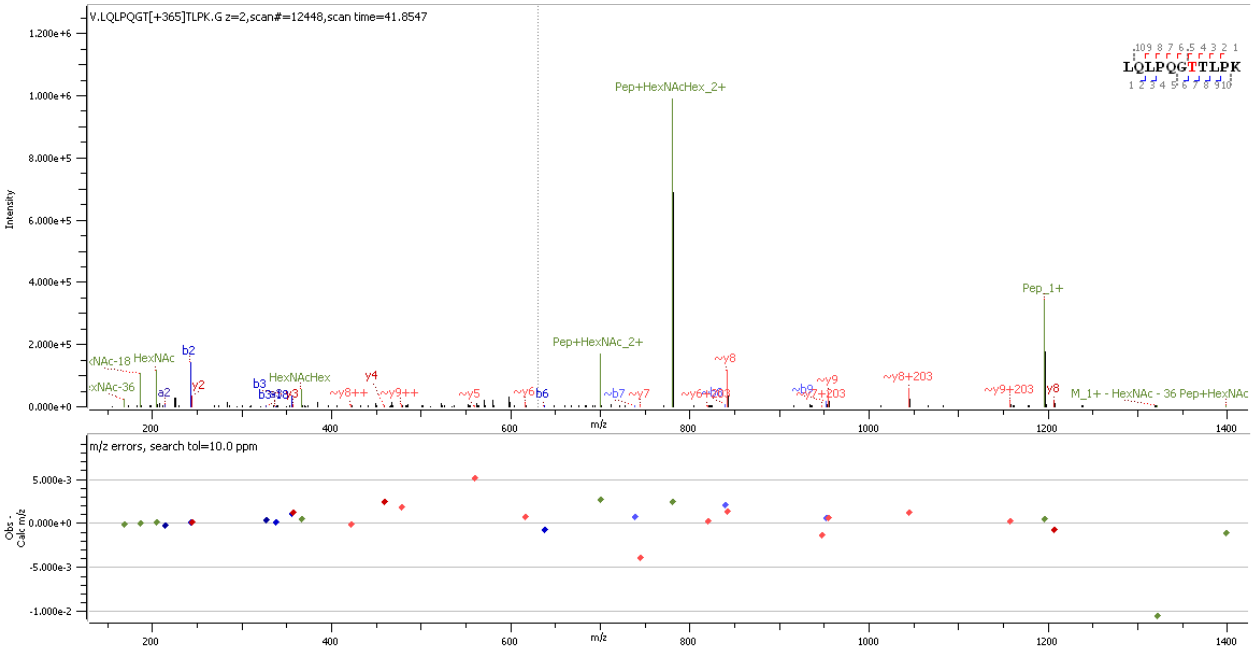


**Supp. Figure S53** HCD MS2 spectrum of O-glycopeptide [159]LQLPQGTTLPK[169]containing HexNAc(1) at site T165/T166

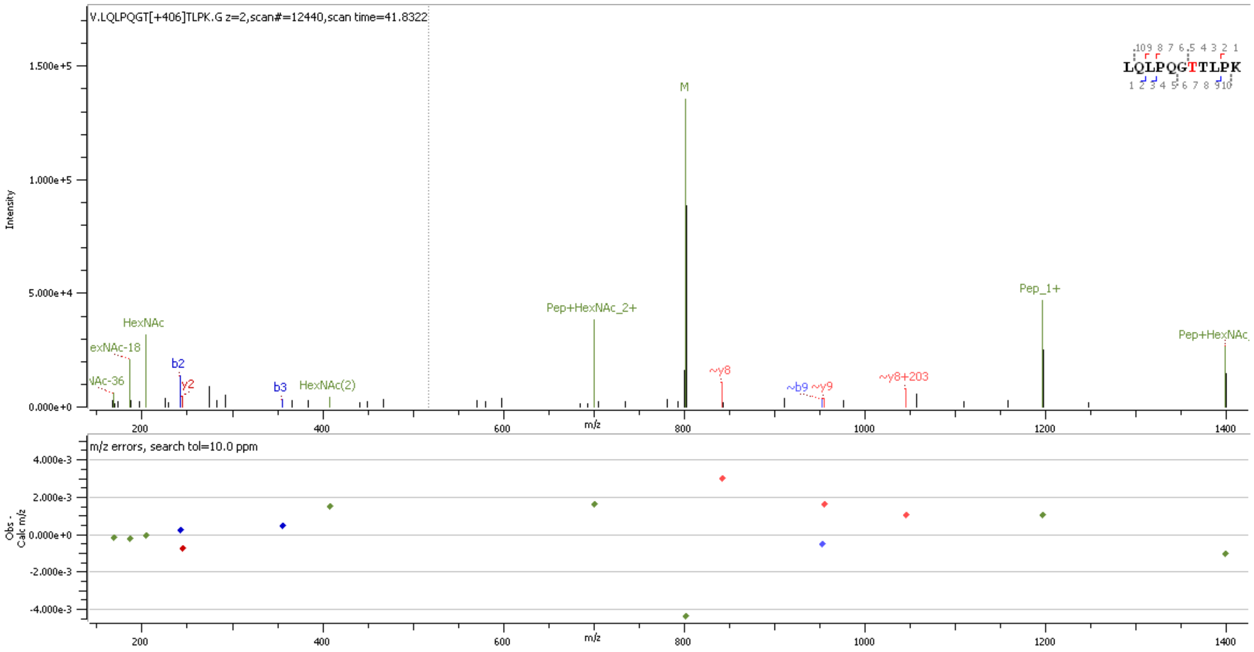


**Supp. Figure S54** HCD MS2 spectrum of O-glycopeptide [159]LQLPQGTTLPK[169]containing HexNAc(2) at site T165/T166

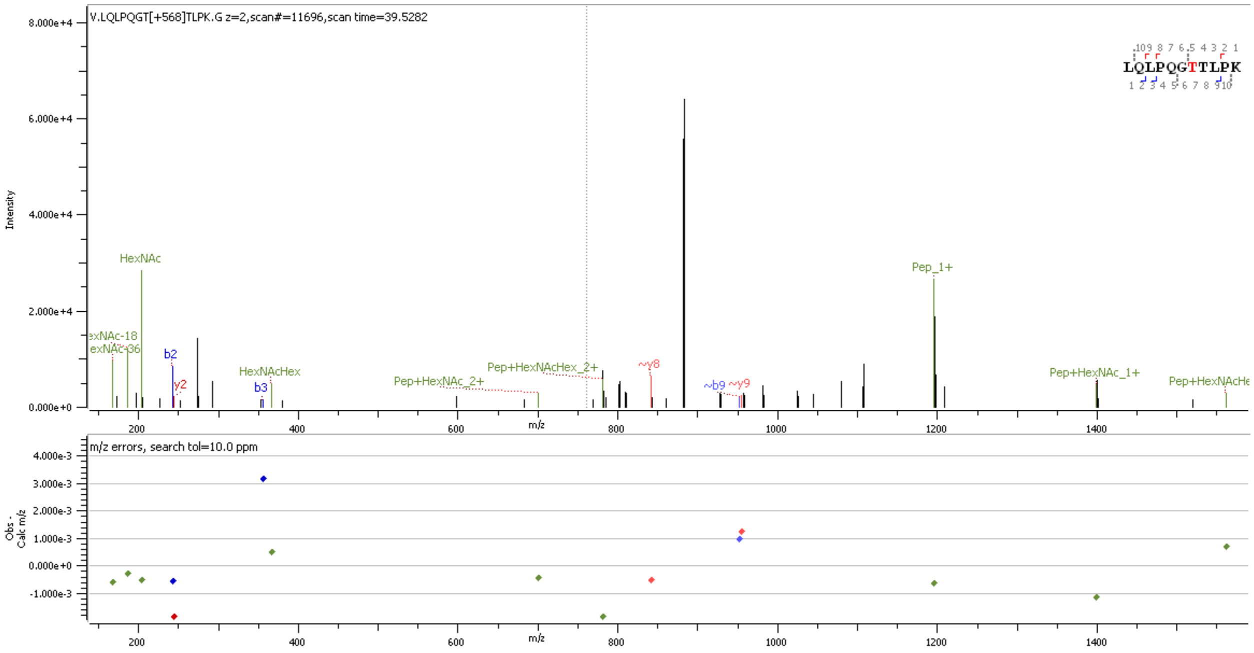


**Supp. Figure S55** HCD MS2 spectrum of O-glycopeptide [159]LQLPQGTTLPK[169]containing HexNAc(2)Hex(1) at site T165/T166

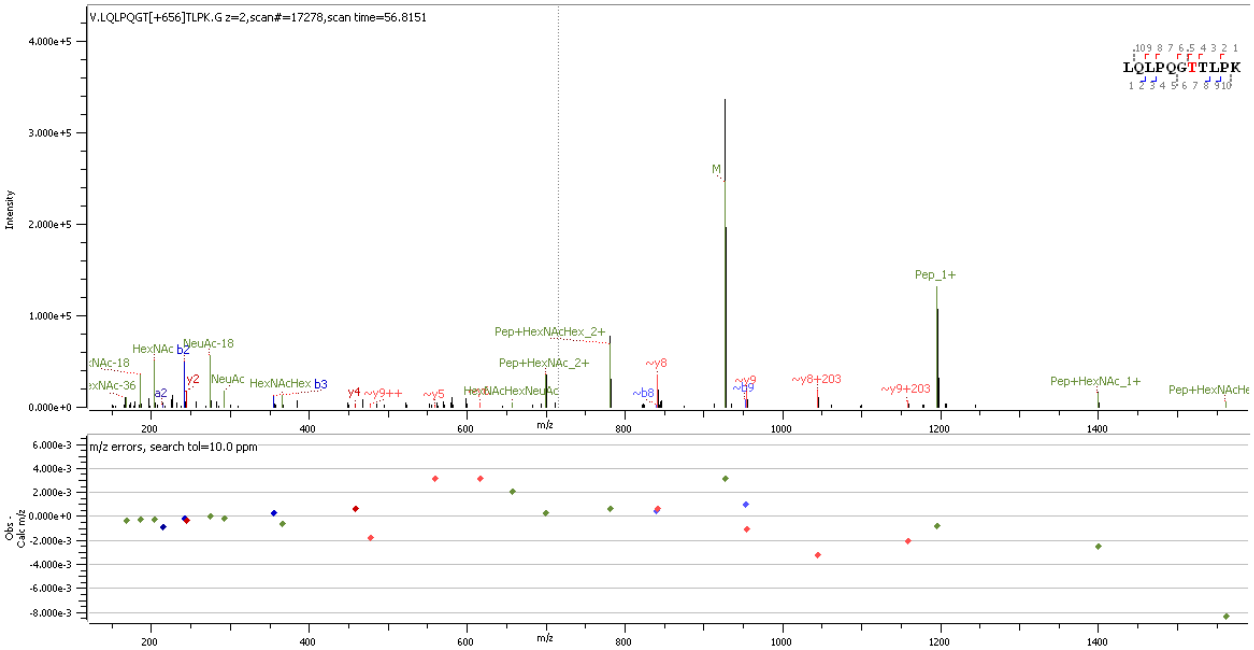

**Supp. Figure S56** HCD MS2 spectrum of O-glycopeptide [159]LQLPQGTTLPK[169]containing HexNAc(1)Hex(1)NeuAc(1) at site T165/T166

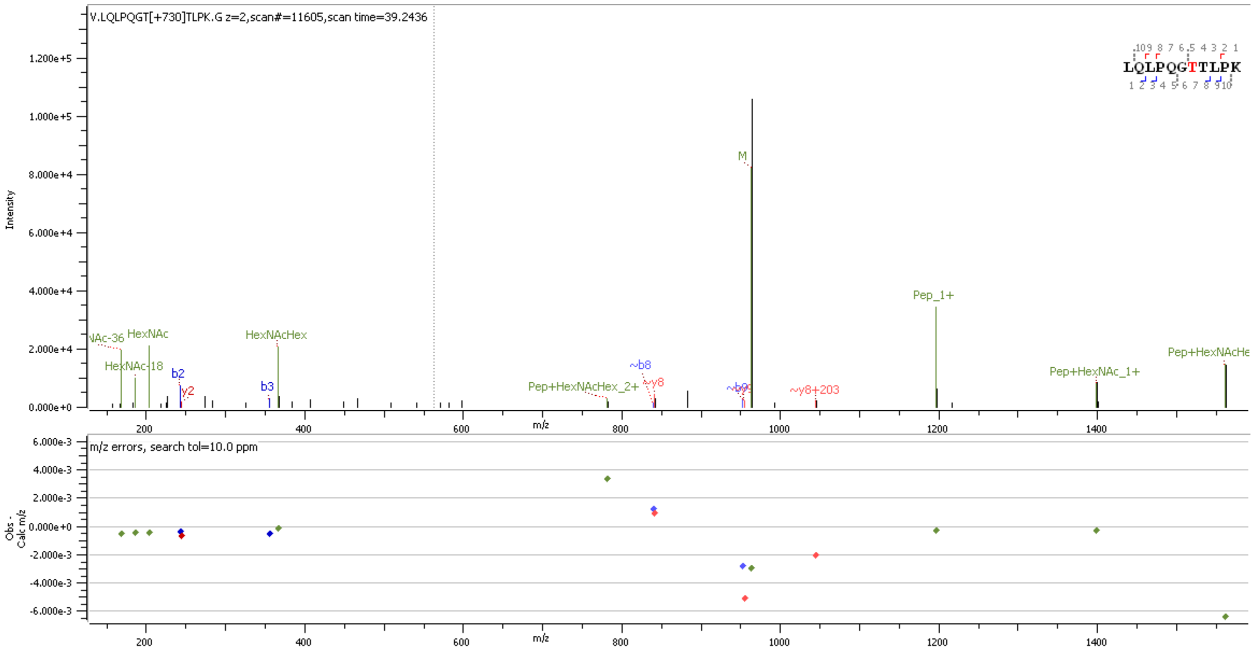

**Supp. Figure S57** HCD MS2 spectrum of O-glycopeptide [159]LQLPQGTTLPK[169]containing HexNAc(2)Hex(2) at site T165/T166

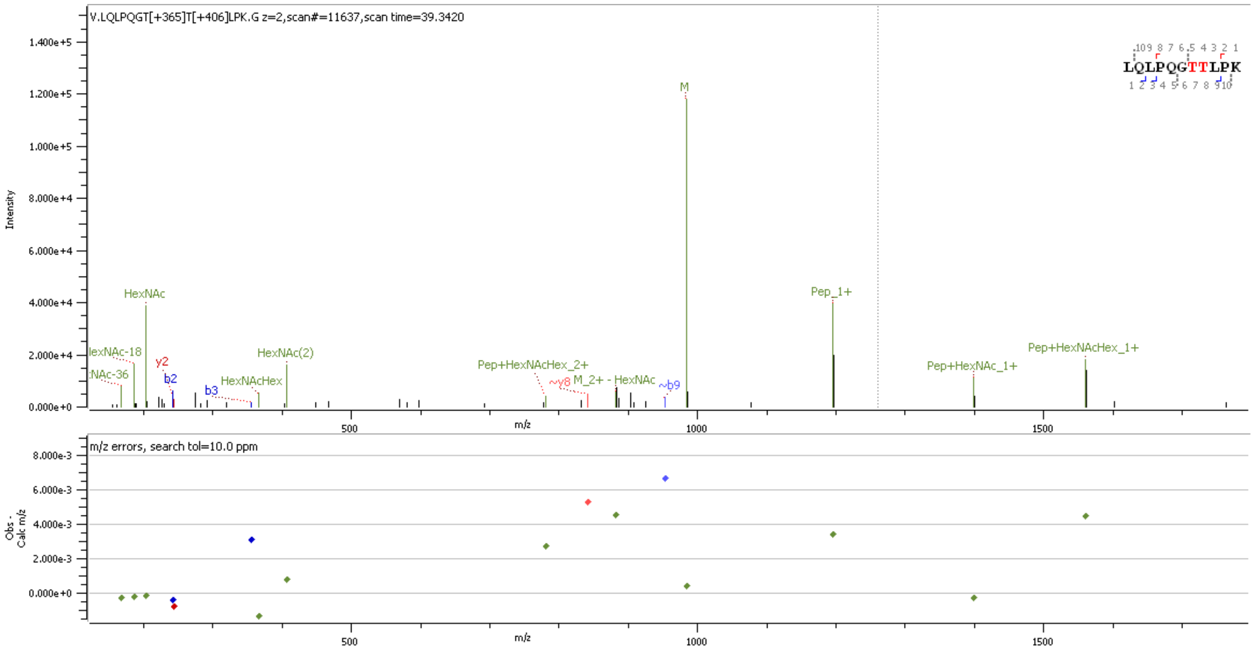


**Supp. Figure S58** HCD MS2 spectrum of O-glycopeptide [159]LQLPQGTTLPK[169]containing HexNAc(1)Hex(1); HexNAc(2) at sites T165 & T166

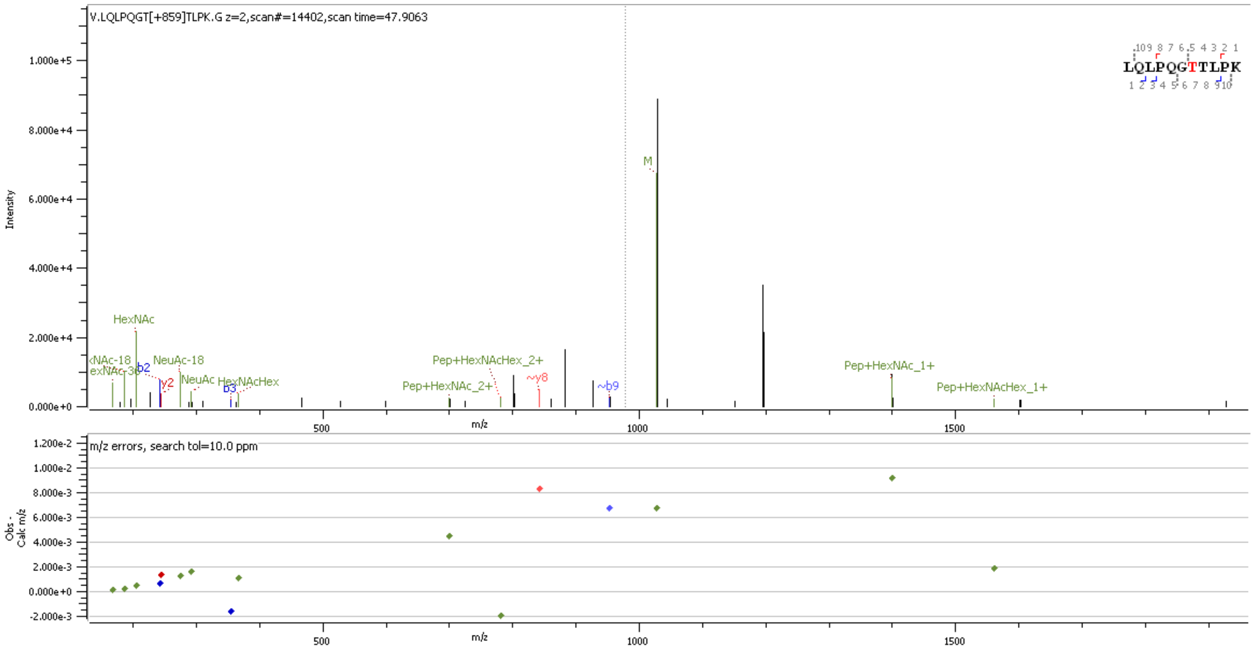


**Supp. Figure S59** HCD MS2 spectrum of O-glycopeptide [159]LQLPQGTTLPK[169]containing HexNAc(2)Hex(1)NeuAc(1) at site T165/T166

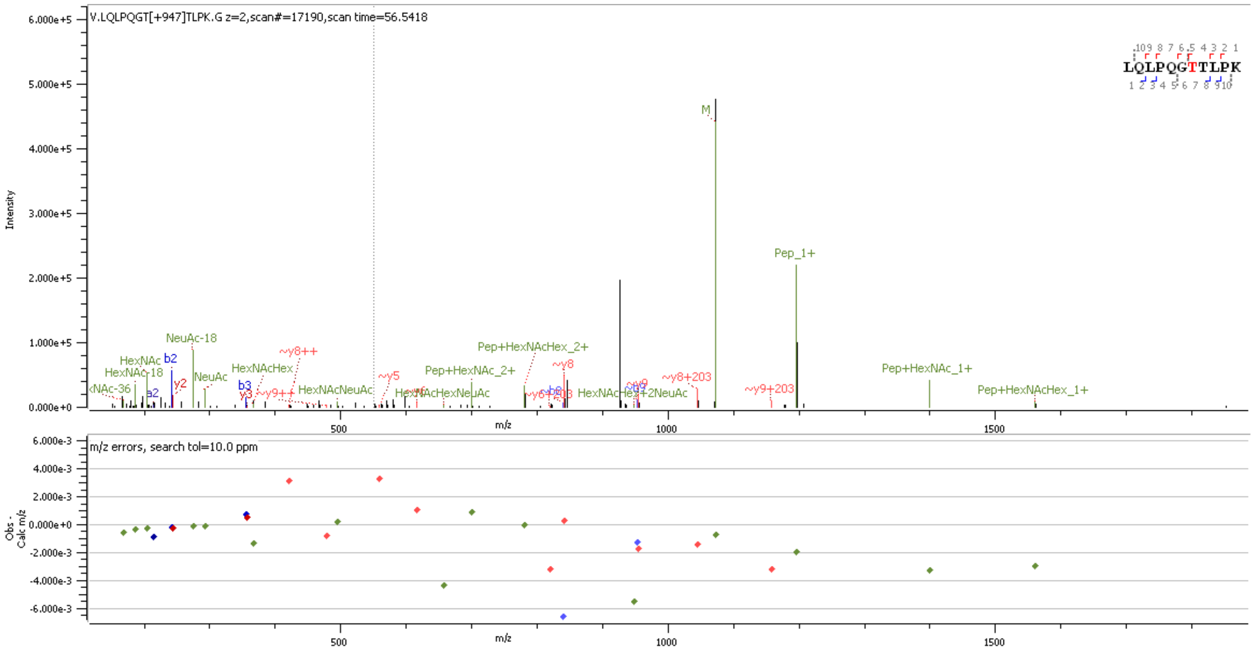


**Supp. Figure S60** HCD MS2 spectrum of O-glycopeptide [159]LQLPQGTTLPK[169]containing HexNAc(1)Hex(1)NeuAc(2) at site T165/T166

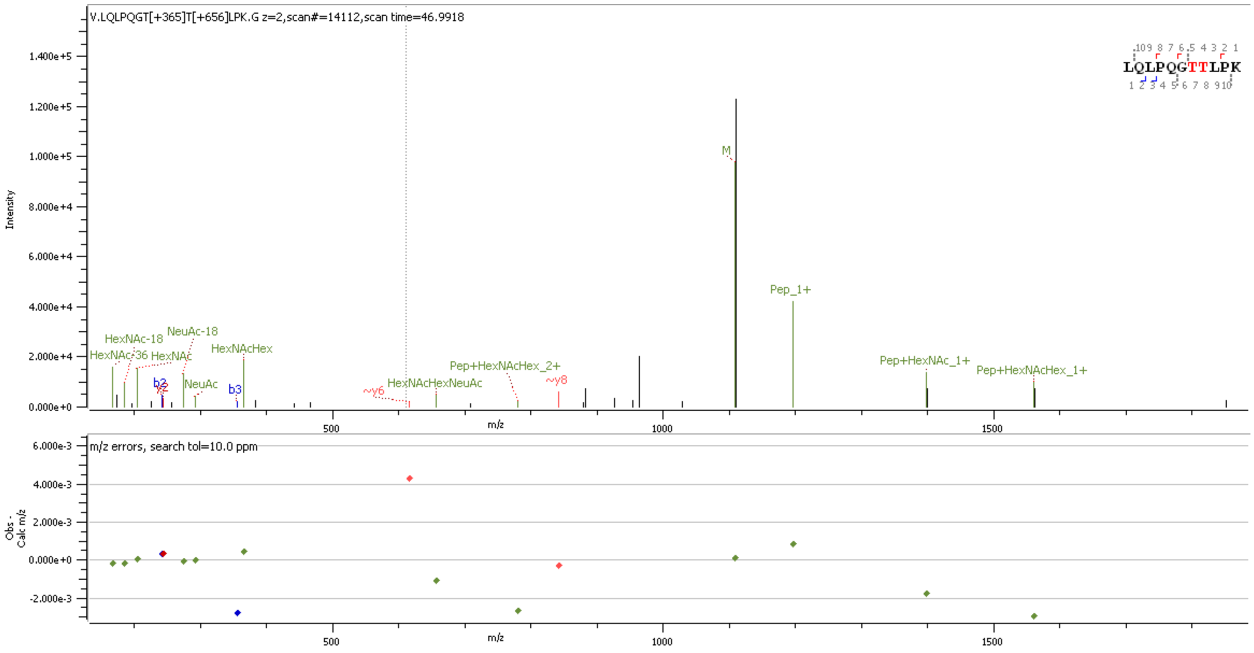


**Supp. Figure S61** HCD MS2 spectrum of O-glycopeptide [159]LQLPQGTTLPK[169]containing HexNAc(1)Hex(1); HexNAc(1)Hex(1)NeuAc(1) at sites T165 & T166

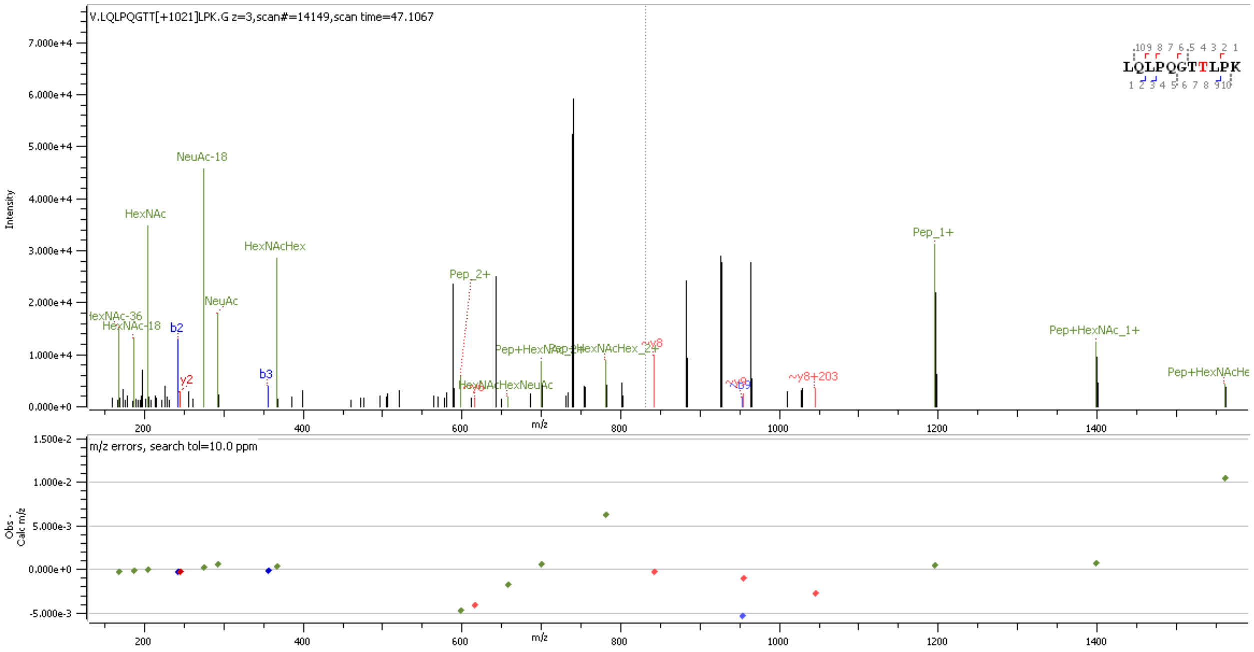


**Supp. Figure S62** HCD MS2 spectrum of O-glycopeptide [159]LQLPQGTTLPK[169]containing HexNAc(2)Hex(2)NeuAc(1) at site T165/T166

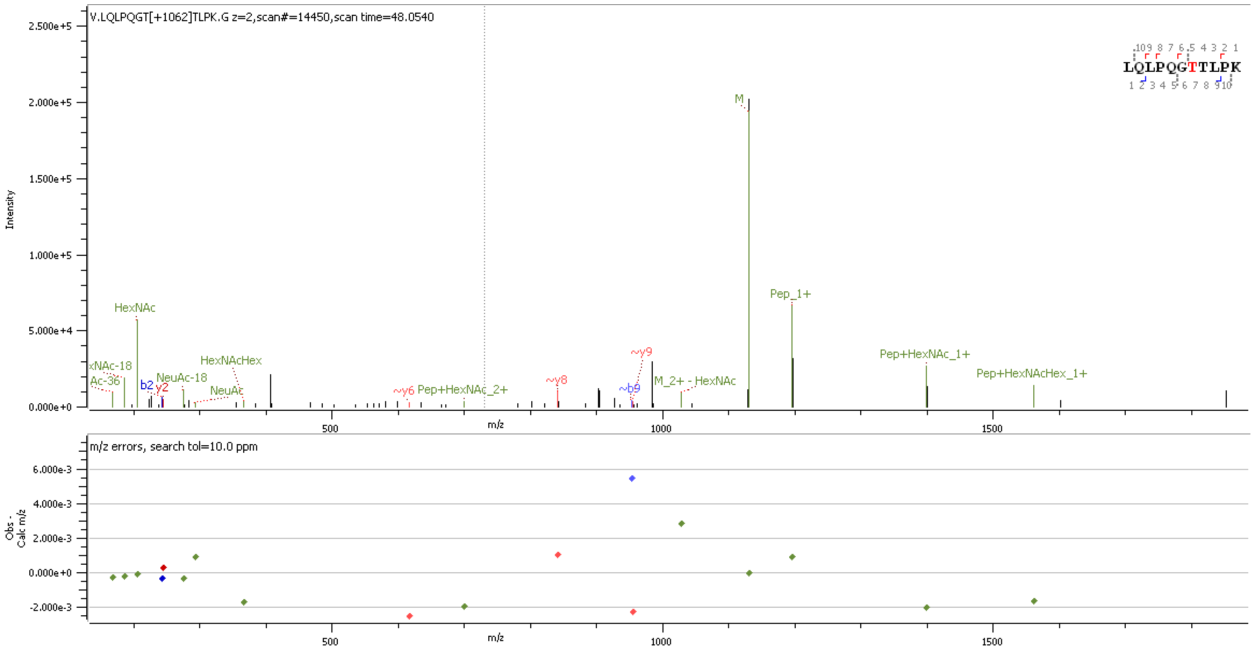


**Supp. Figure S63** HCD MS2 spectrum of O-glycopeptide [159]LQLPQGTTLPK[169]containing HexNAc(3)Hex(1)NeuAc(1) at site T165/T166

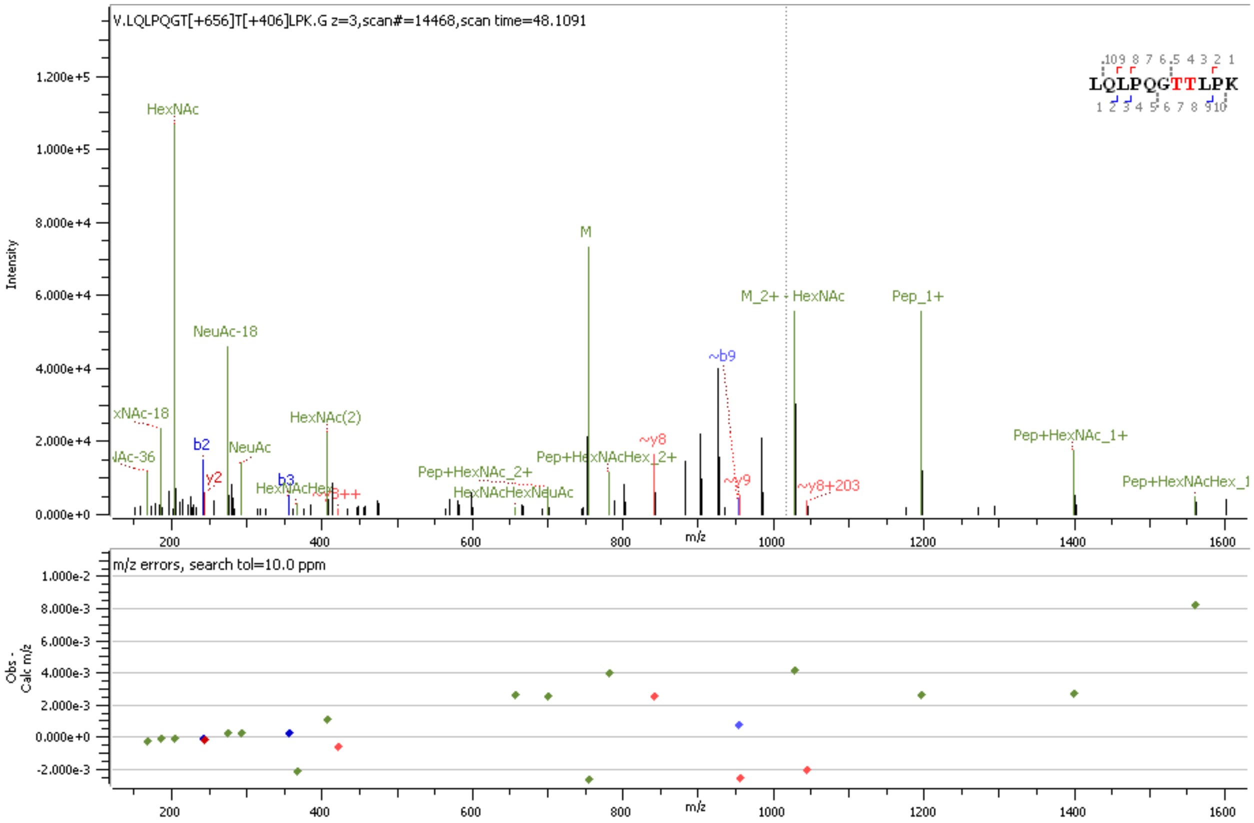


**Supp. Figure S64** HCD MS2 spectrum of O-glycopeptide [159]LQLPQGTTLPK[169]containing HexNAc(1)Hex(1)NeuAc(1); HexNAc(2) at sites T16 & T166

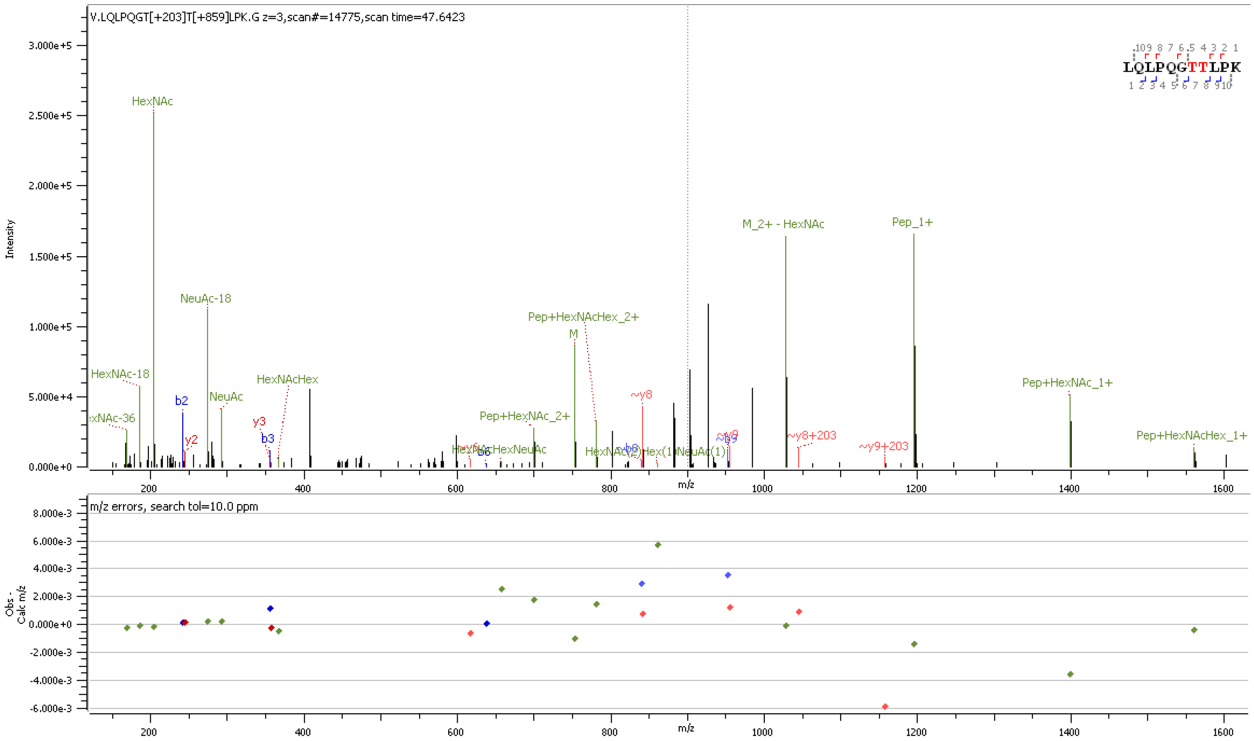


**Supp. Figure S65** HCD MS2 spectrum of O-glycopeptide [159]LQLPQGTTLPK[169]containing HexNAc(1); HexNAc(2)Hex(1)NeuAc(1) at sites T16 & T166

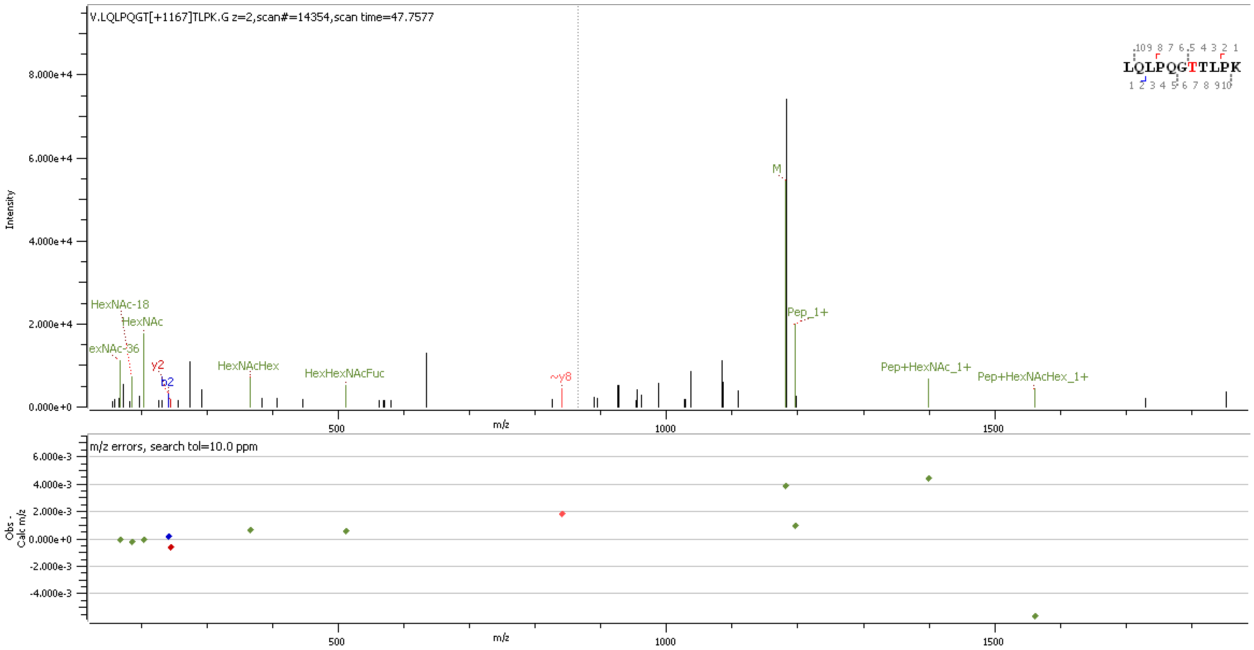


**Supp. Figure S66** HCD MS2 spectrum of O-glycopeptide [159]LQLPQGTTLPK[169]containing HexNAc(2)Hex(2)Fuc(1)NeuAc(1) at site T165/T166

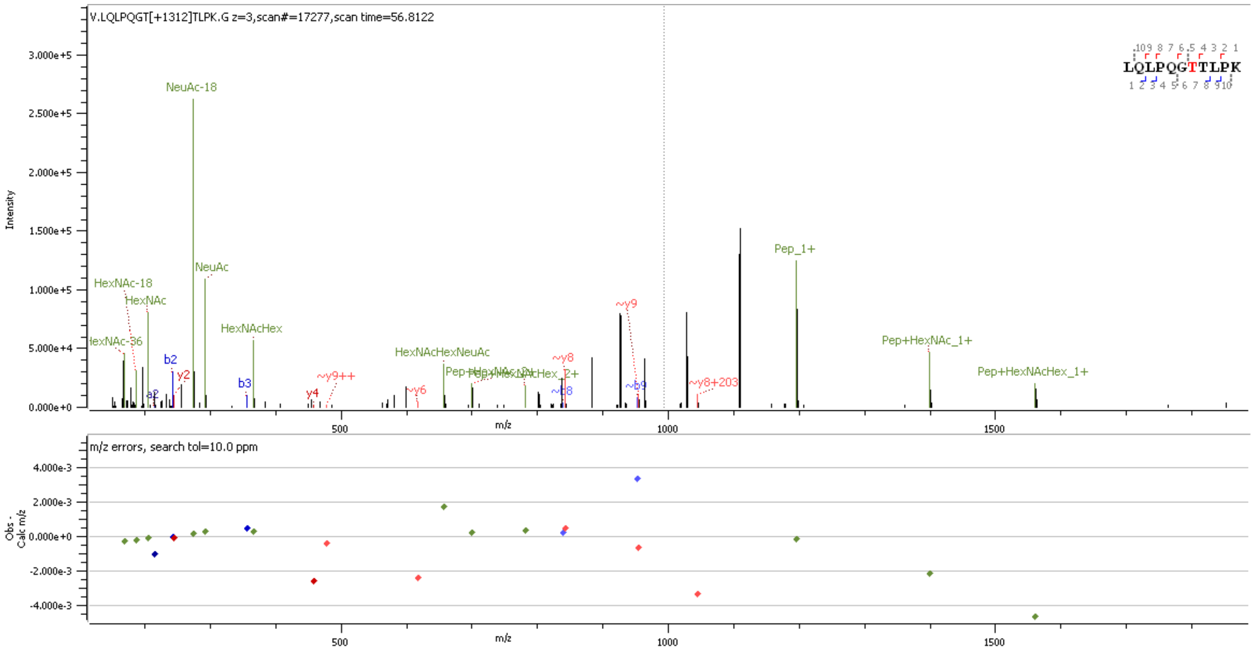


**Supp. Figure S67** HCD MS2 spectrum of O-glycopeptide [159]LQLPQGTTLPK[169]containing HexNAc(2)Hex(2)NeuAc(2) at site T165/T166

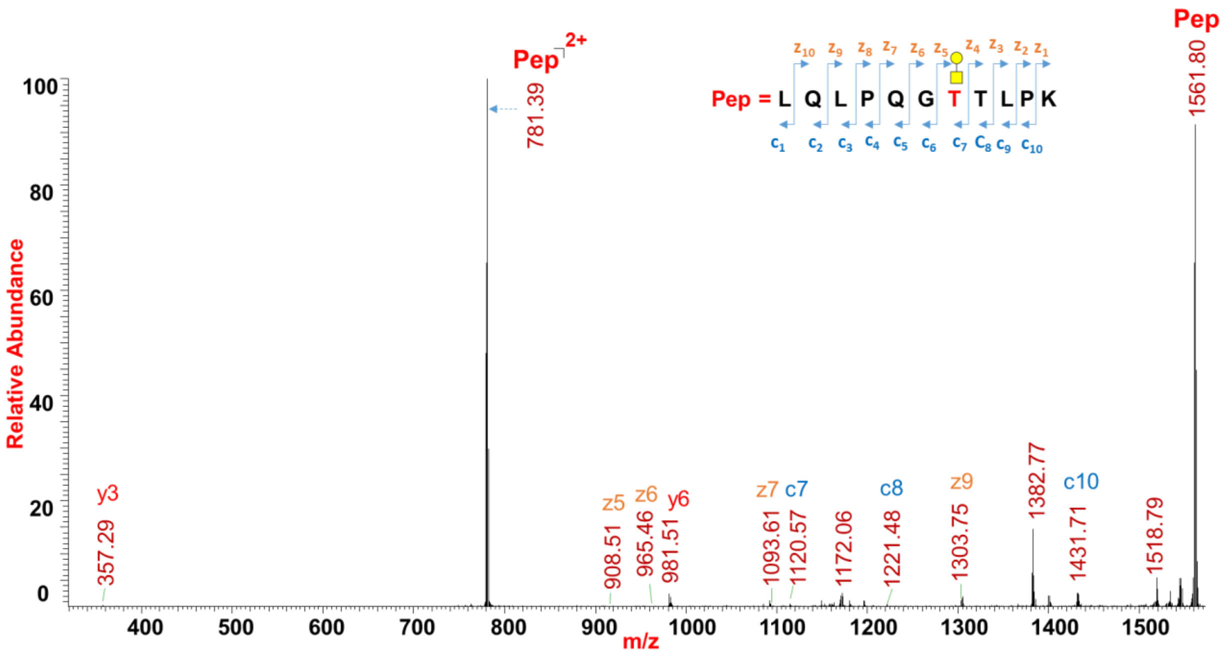


**Supp. Figure S68** ETD MS2 spectrum of O-glycopeptide [159]LQLPQGTTLPK[169]containing HexNAc(1)Hex(1) at site T165

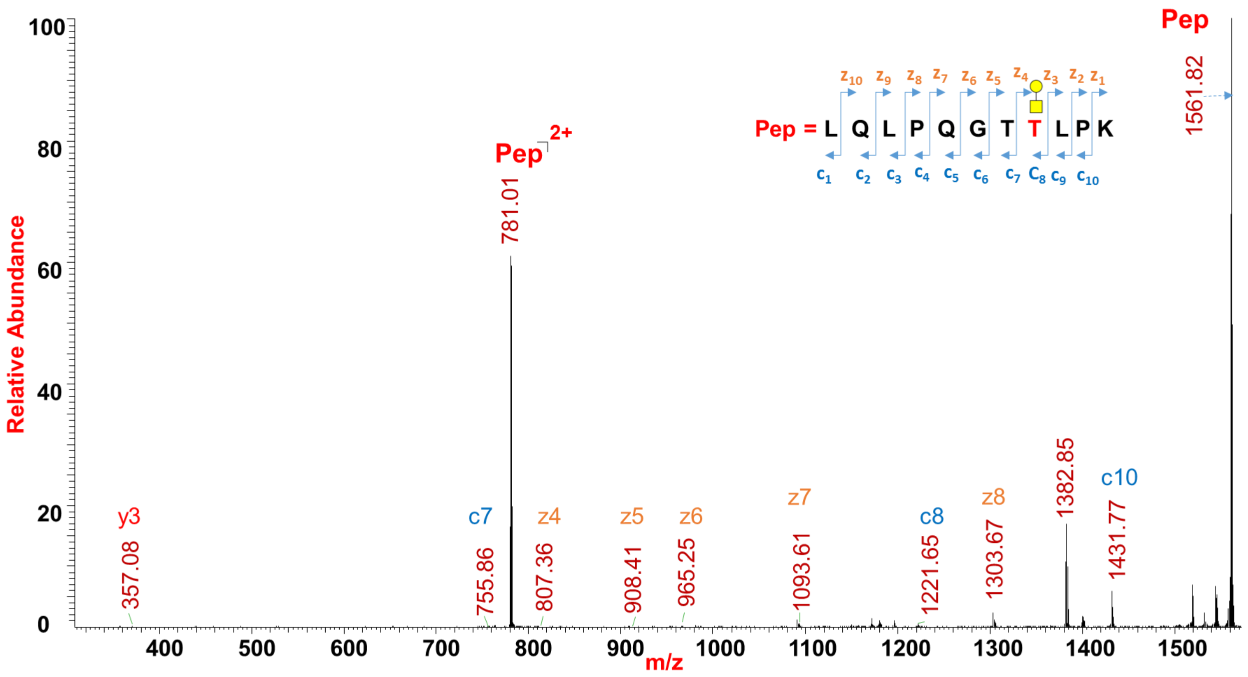


**Supp. Figure S69** ETD MS2 spectrum of O-glycopeptide [159]LQLPQGTTLPK[169]containing HexNAc(1)Hex(1) at site T166

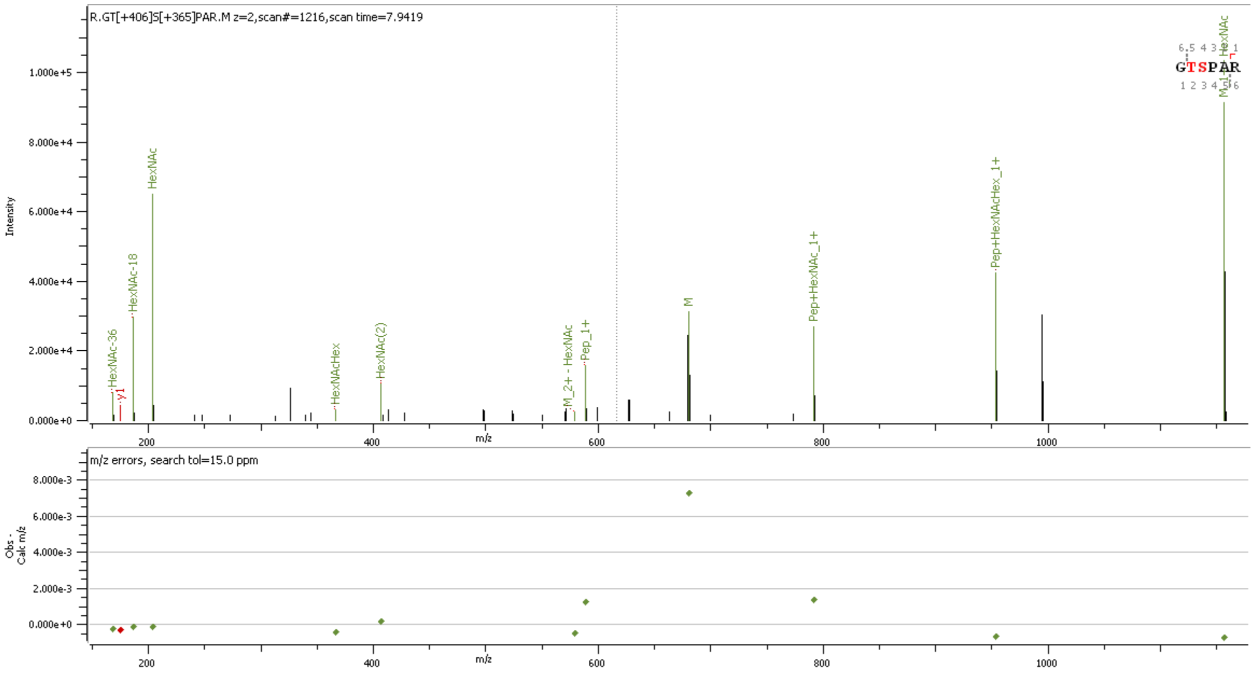


**Supp. Figure S70** HCD MS2 spectrum of O-glycopeptide [204]GTSPAR[209]containing HexNAc (2); HexNAc(1)Hex(1) at sites T205 & S206

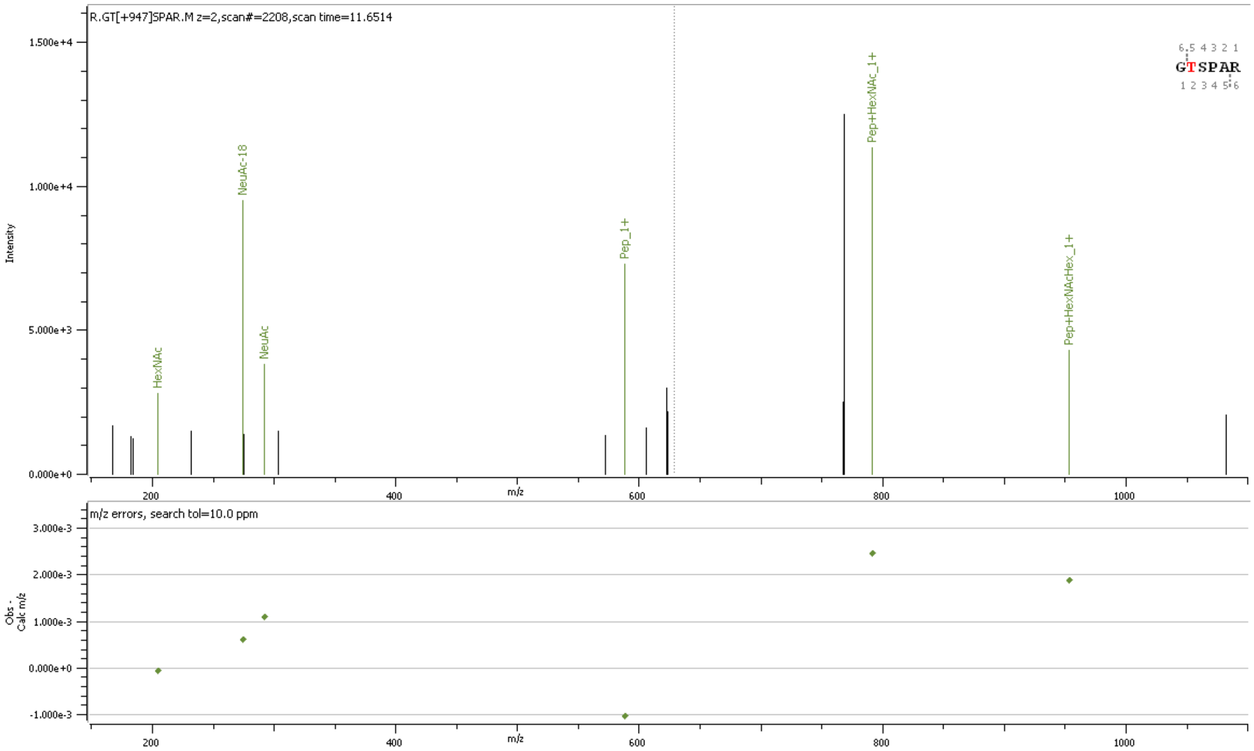


**Supp. Figure S71** HCD MS2 spectrum of O-glycopeptide [204]GTSPAR[209]containing HexNAc(1)Hex(1) NeuAc(2) at site T205/S206

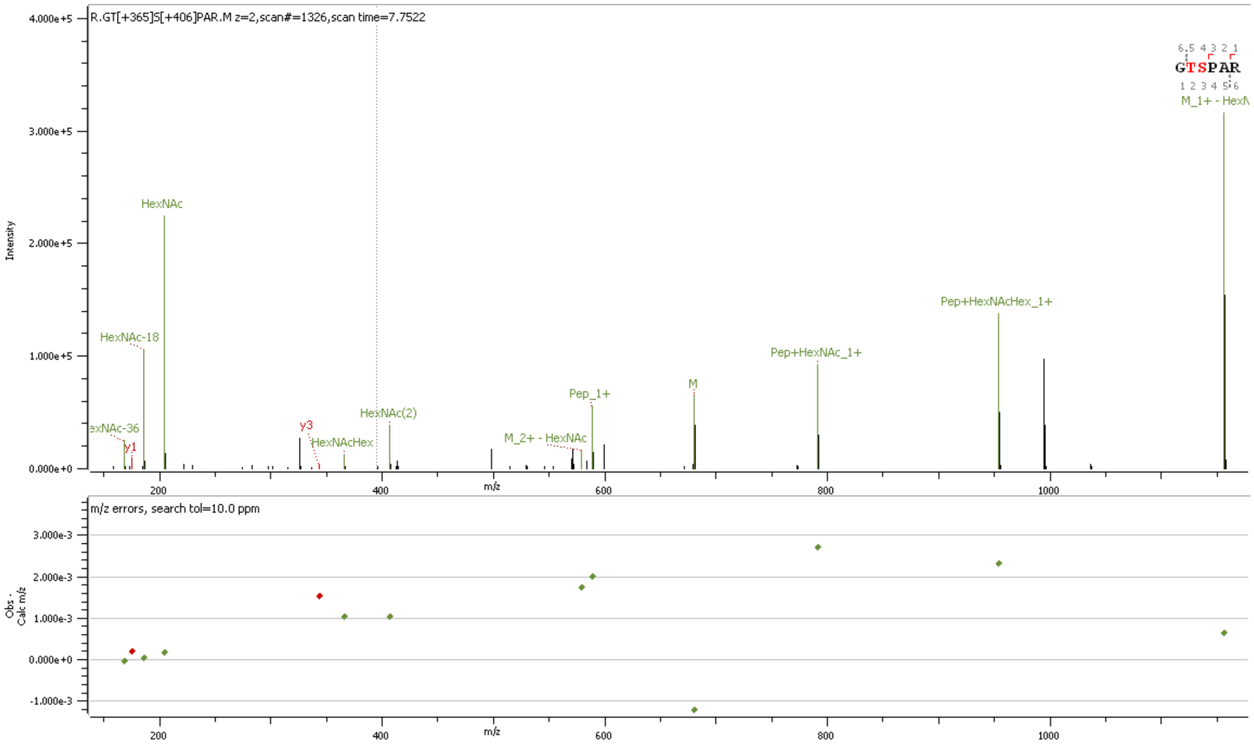


**Supp. Figure S72** HCD MS2 spectrum of O-glycopeptide [204]GTSPAR[209]containing HexNAc(1)Hex(1); HexNAc(2) at sites T205 & S206

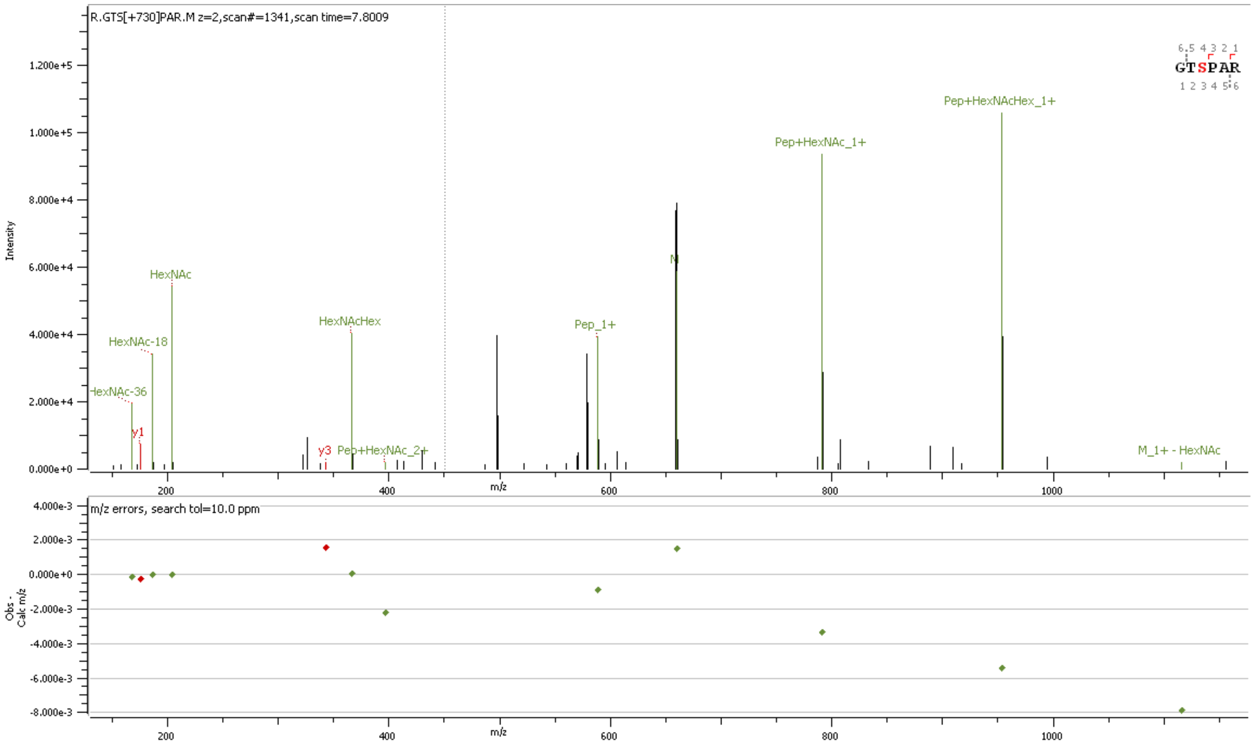


**Supp. Figure S73** HCD MS2 spectrum of O-glycopeptide [204]GTSPAR[209]containing HexNAc(2)Hex(2) at site T205/S206

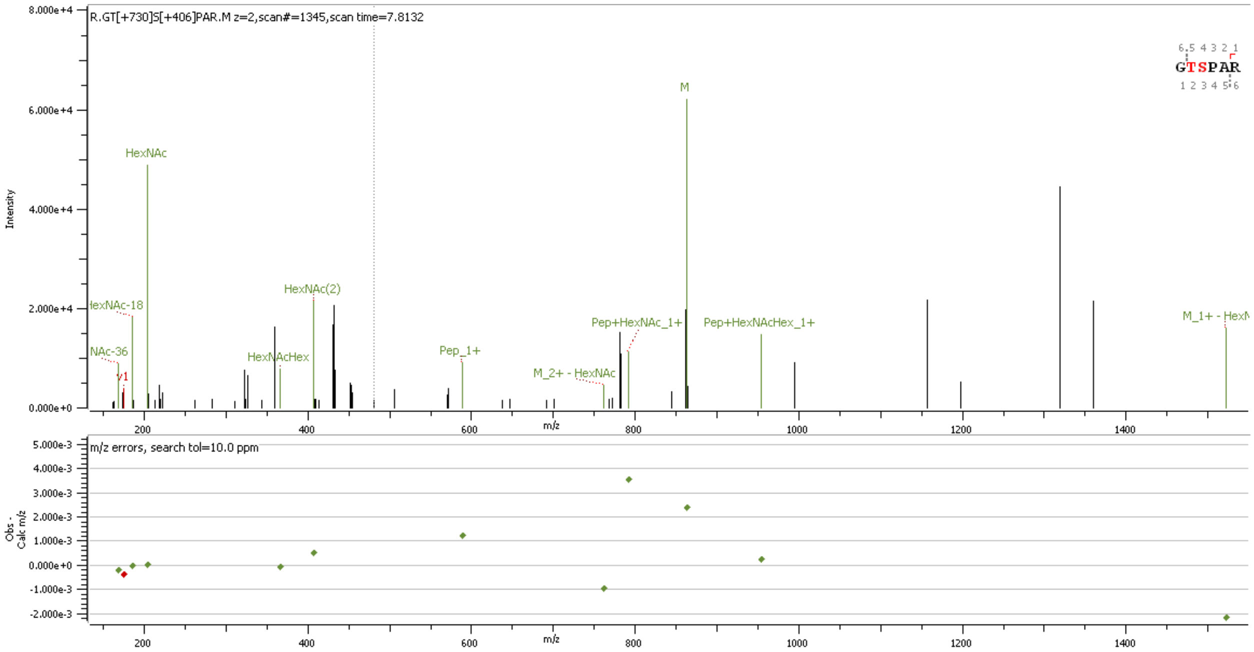


**Supp. Figure S74** HCD MS2 spectrum of O-glycopeptide [204]GTSPAR[209]containing HexNAc(2)Hex(2); HexNAc(2) at sites T205 & S206

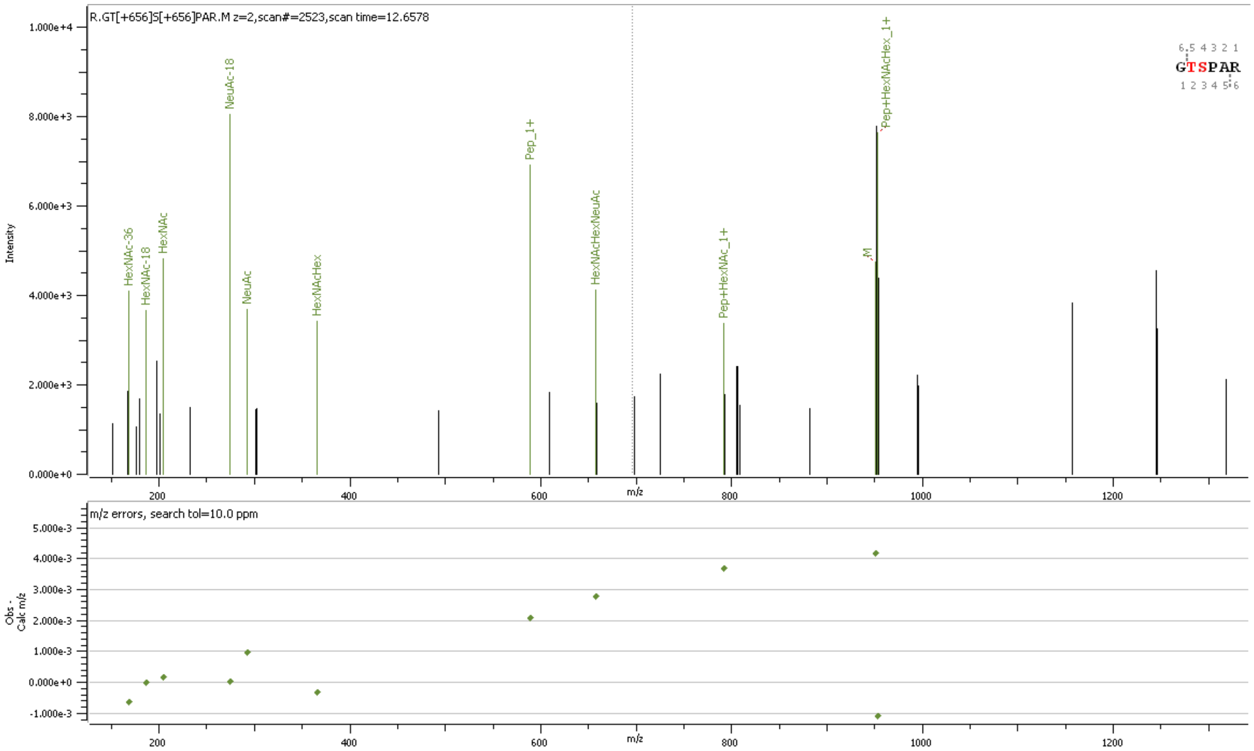


**Supp. Figure S75** HCD MS2 spectrum of O-glycopeptide [204]GTSPAR[209]containing HexNAc(1)Hex(1)NeuAc(1); HexNAc(1)Hex(1)NeuAc(1) at sites T205 & S206

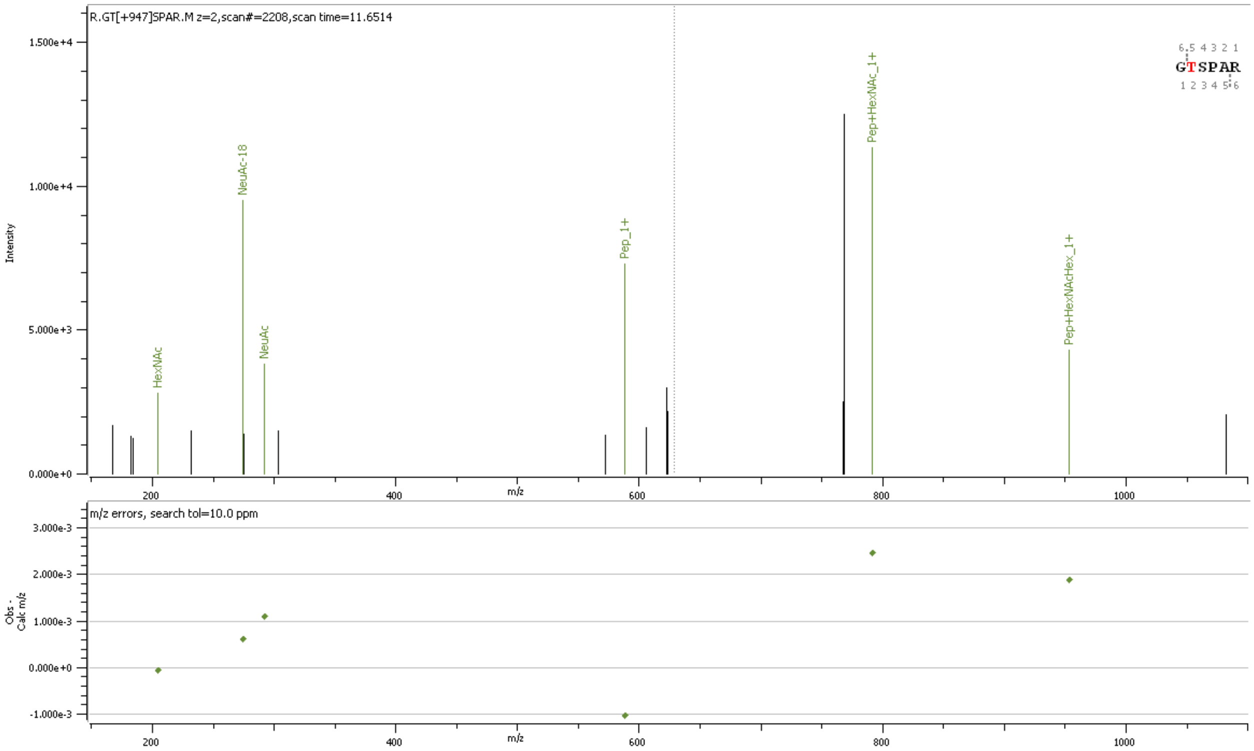


**Supp. Figure S76** HCD MS2 spectrum of O-glycopeptide [204]GTSPAR[209]containing HexNAc(1)Hex(1)NeuAc(2) at site T205/S206

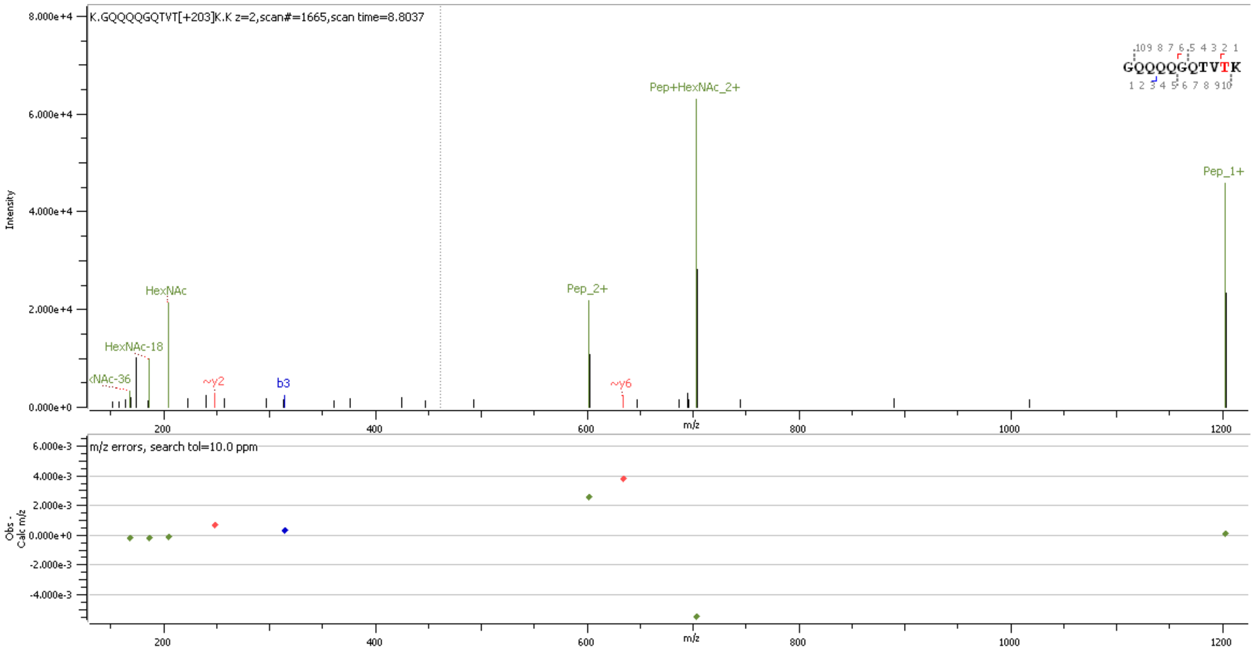


**Supp. Figure S77** HCD MS2 spectrum of O-glycopeptide [238]GQQQQGQTVTK[248]containing HexNAc(1) at site T245/T247

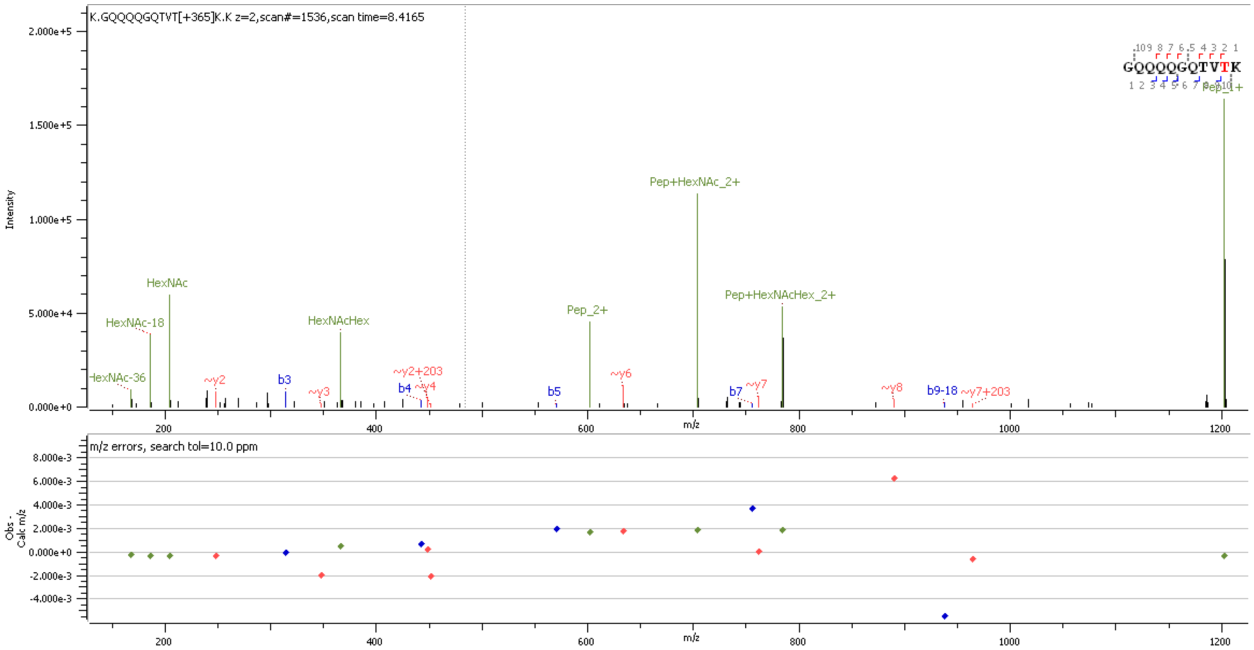


**Supp. Figure S78** HCD MS2 spectrum of O-glycopeptide [238]GQQQQGQTVTK[248]containing HexNAc(1)Hex(1) at site T245/T247

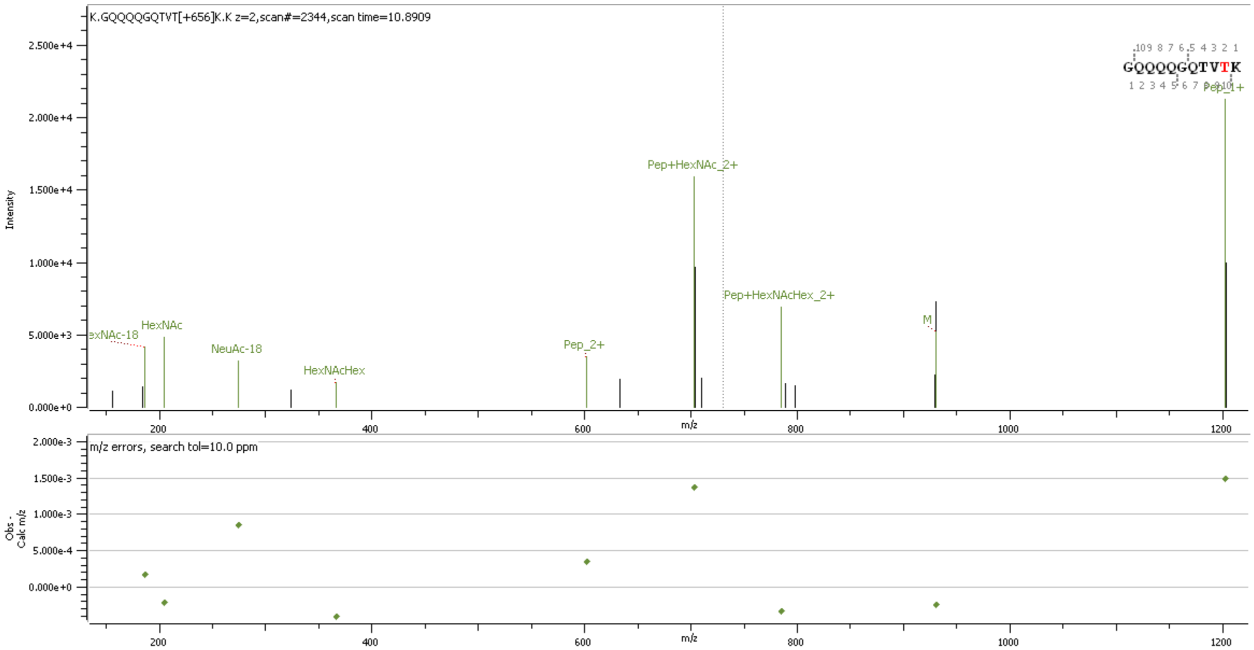


**Supp. Figure S79** HCD MS2 spectrum of O-glycopeptide [238]GQQQQGQTVTK[248]containing HexNAc(1)Hex(1)NeuAc(1) at site T245/T247

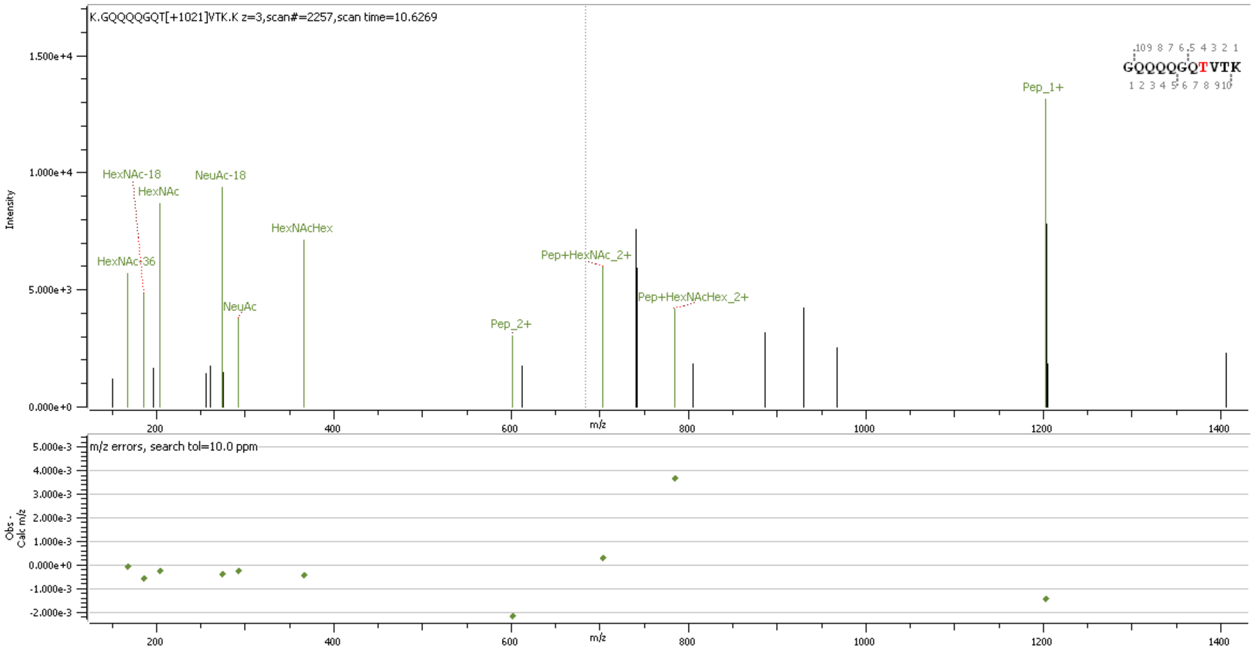


**Supp. Figure S80** HCD MS2 spectrum of O-glycopeptide [238]GQQQQGQTVTK[248]containing HexNAc(2)Hex(2)NeuAc(1) at site T245/T247

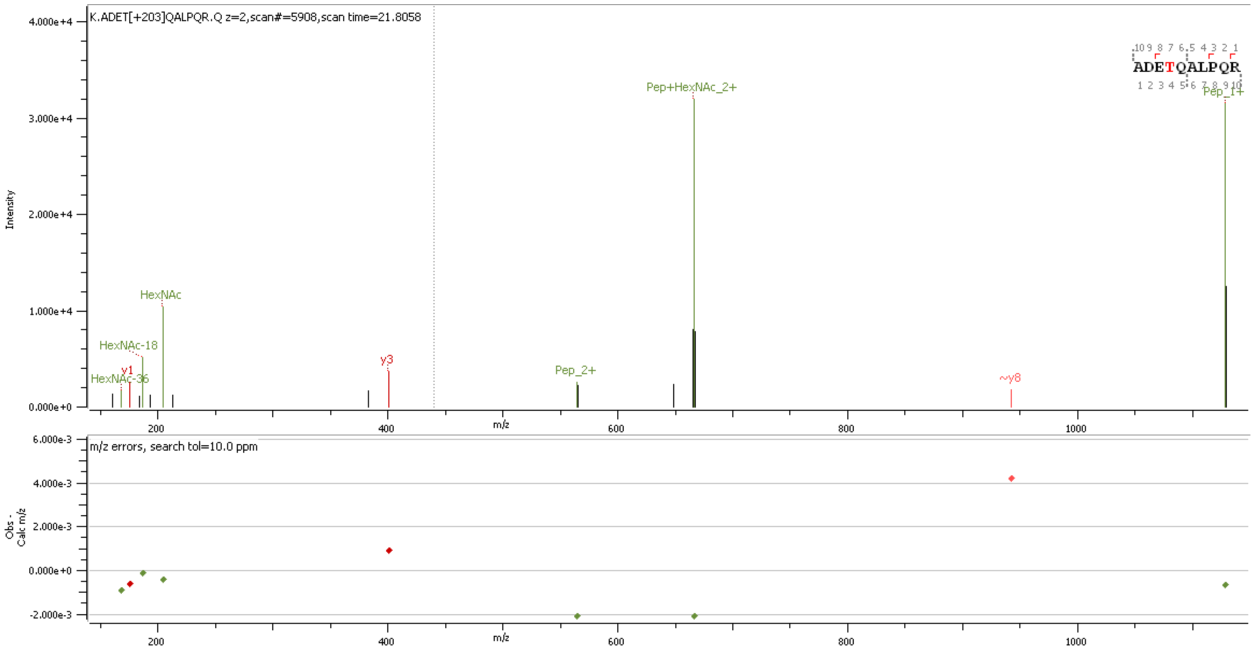


**Supp. Figure S81** HCD MS2 spectrum of O-glycopeptide [376]ADETQALPQR[385] containing HexNAc(1) at site T379

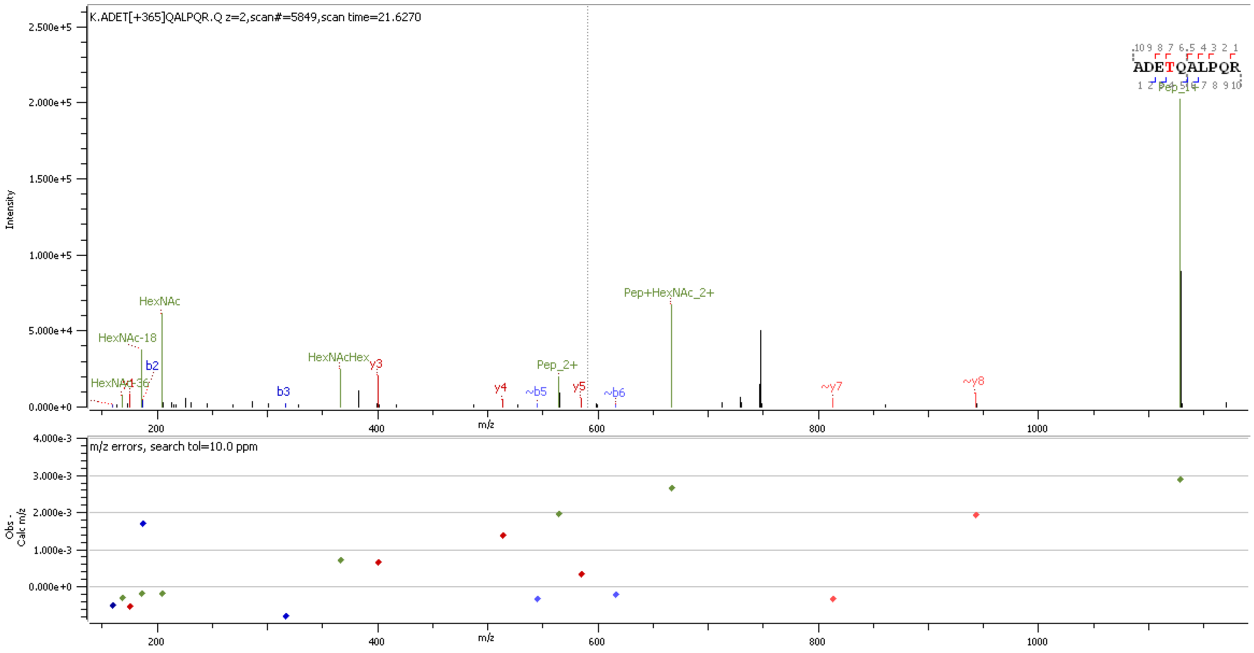


**Supp. Figure S82** HCD MS2 spectrum of O-glycopeptide [376]ADETQALPQR[385] containing HexNAc(1)Hex(1) at site T379

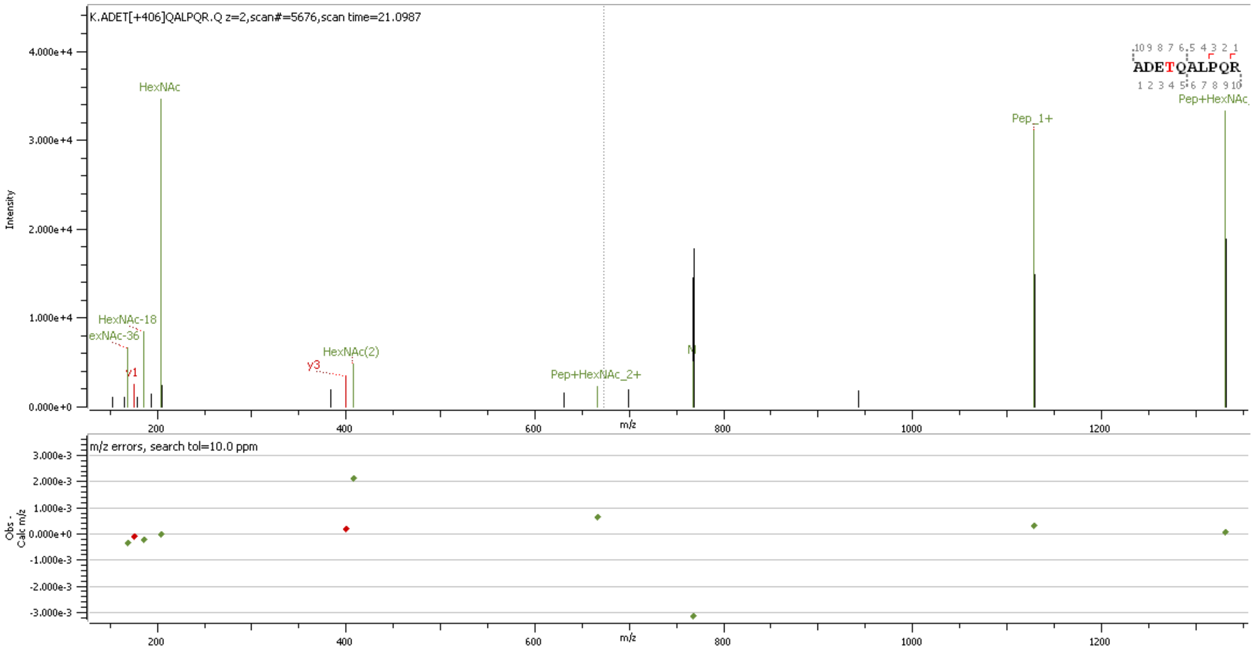


**Supp. Figure S83** HCD MS2 spectrum of O-glycopeptide [376]ADETQALPQR[385] containing HexNAc(2) at site T379

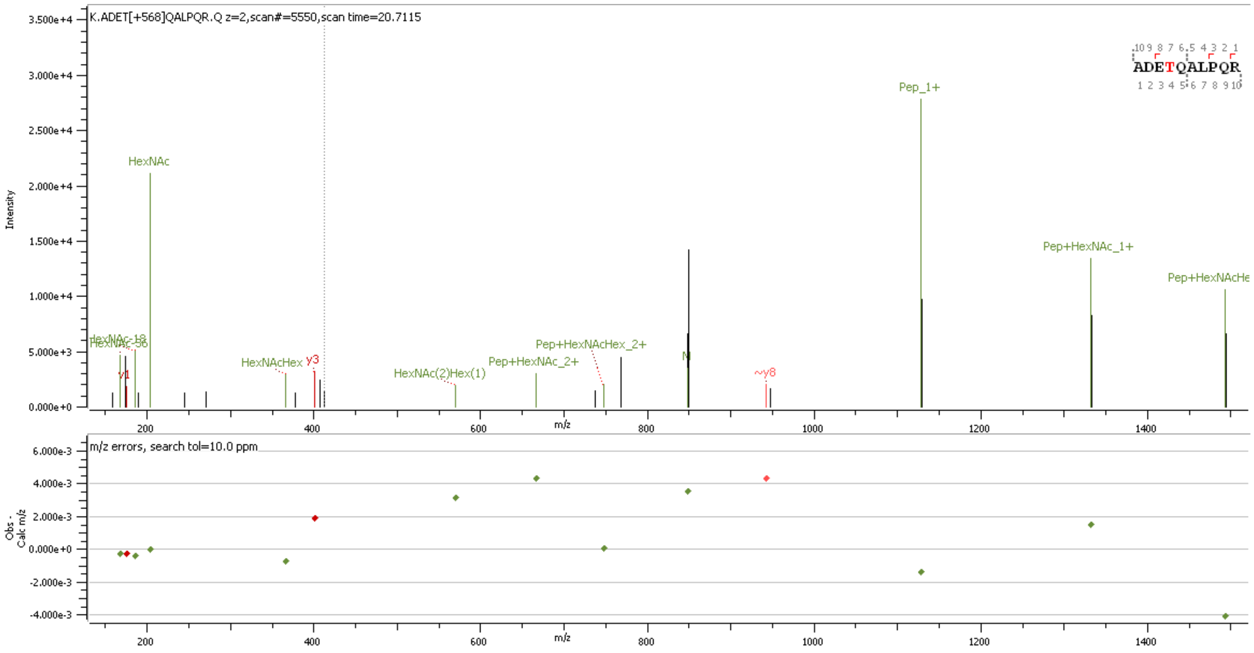


**Supp. Figure S84** HCD MS2 spectrum of O-glycopeptide [376]ADETQALPQR[385] containing HexNAc(2)Hex(1) at site T379

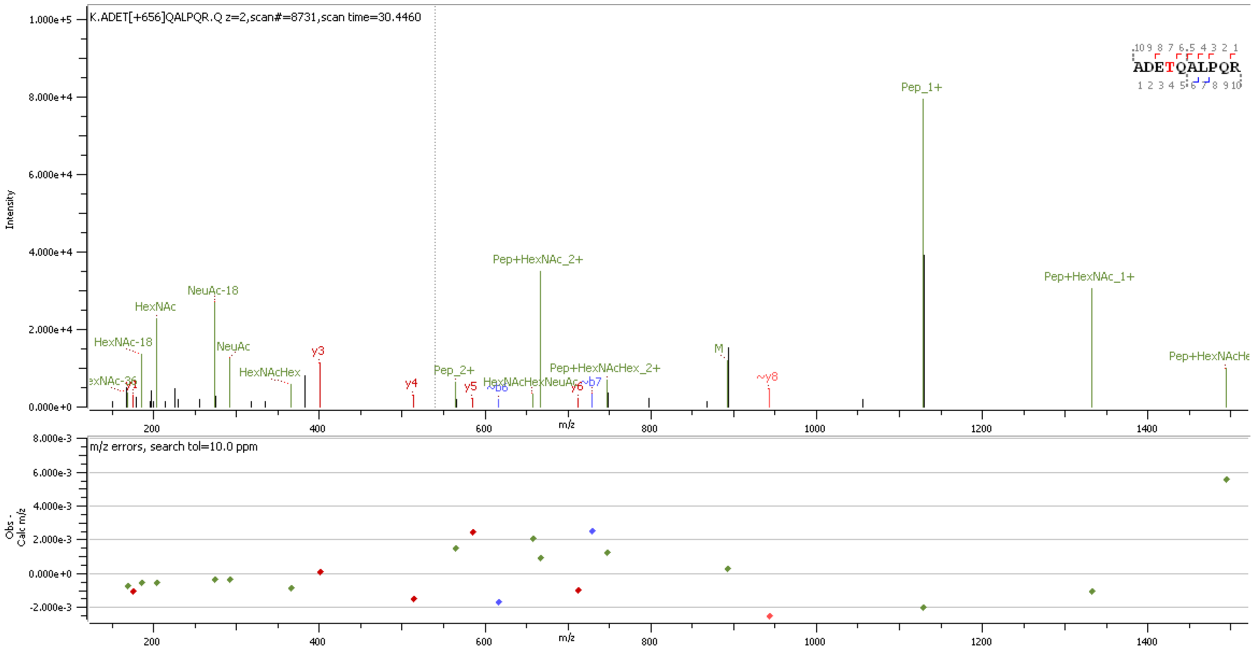


**Supp. Figure S85** HCD MS2 spectrum of O-glycopeptide [376]ADETQALPQR[385] containing HexNAc(1)Hex(1)NeuAc(1) at site T379

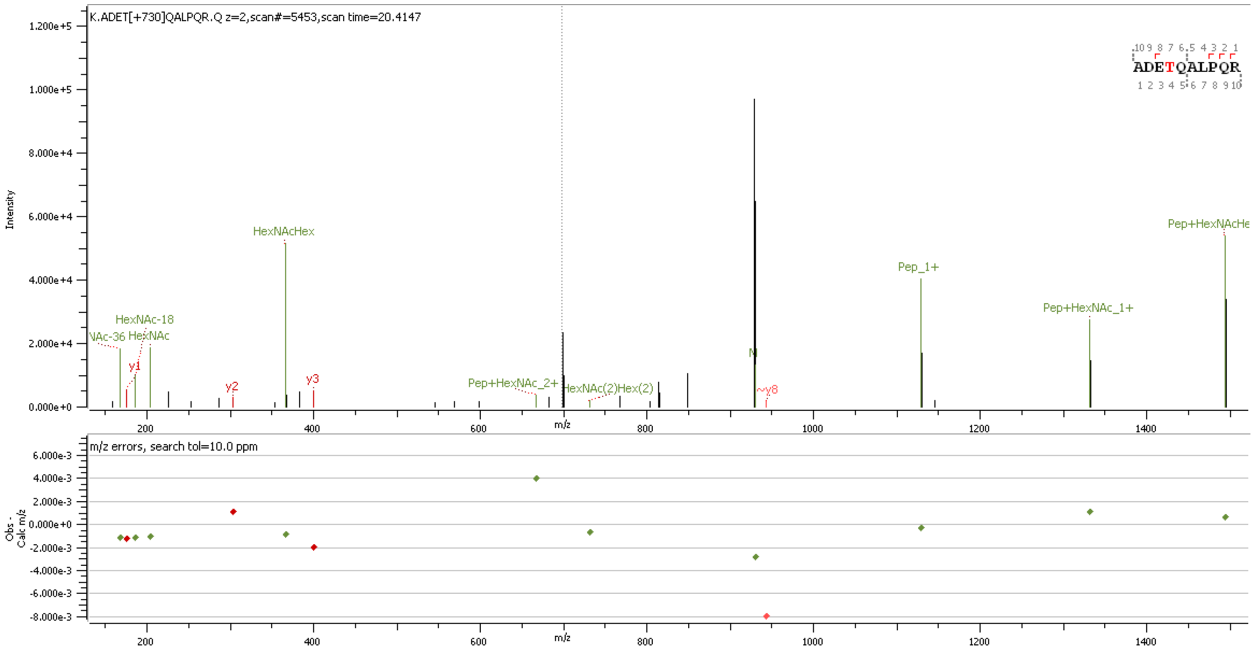


**Supp. Figure S86** HCD MS2 spectrum of O-glycopeptide [376]ADETQALPQR[385] containing HexNAc(2)Hex(2) at site T379

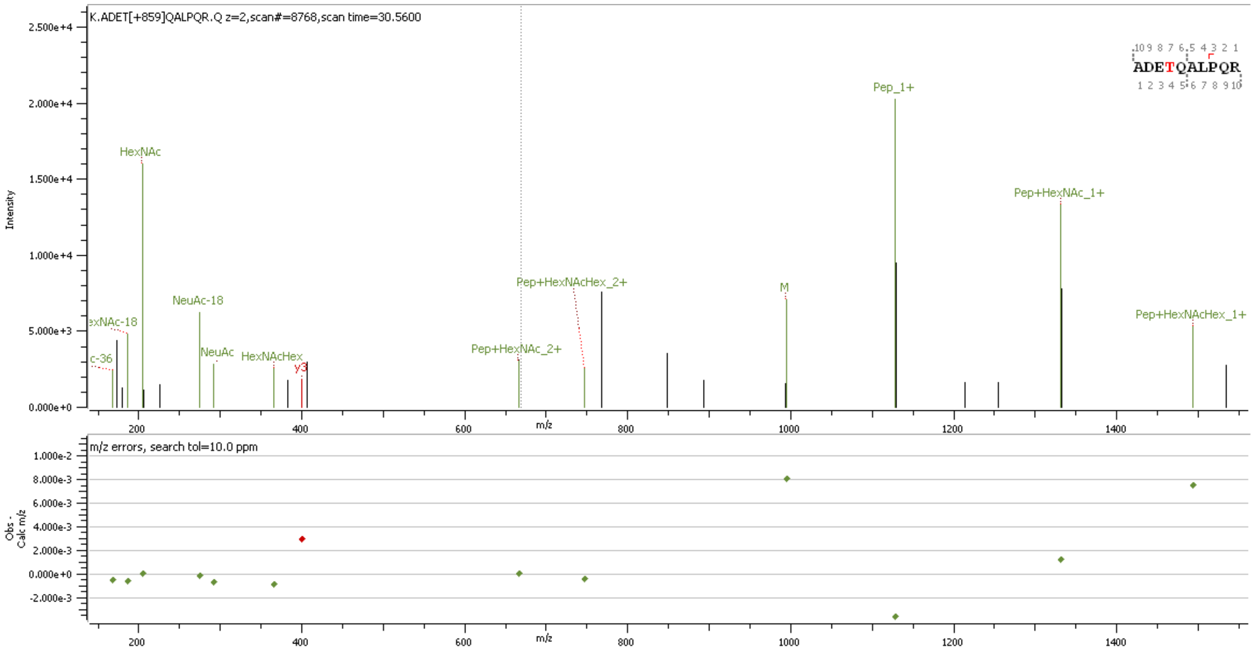

**Supp. Figure S87** HCD MS2 spectrum of O-glycopeptide [376]ADETQALPQR[385] containing HexNAc(2)Hex(1)NeuAc(1) at site T379

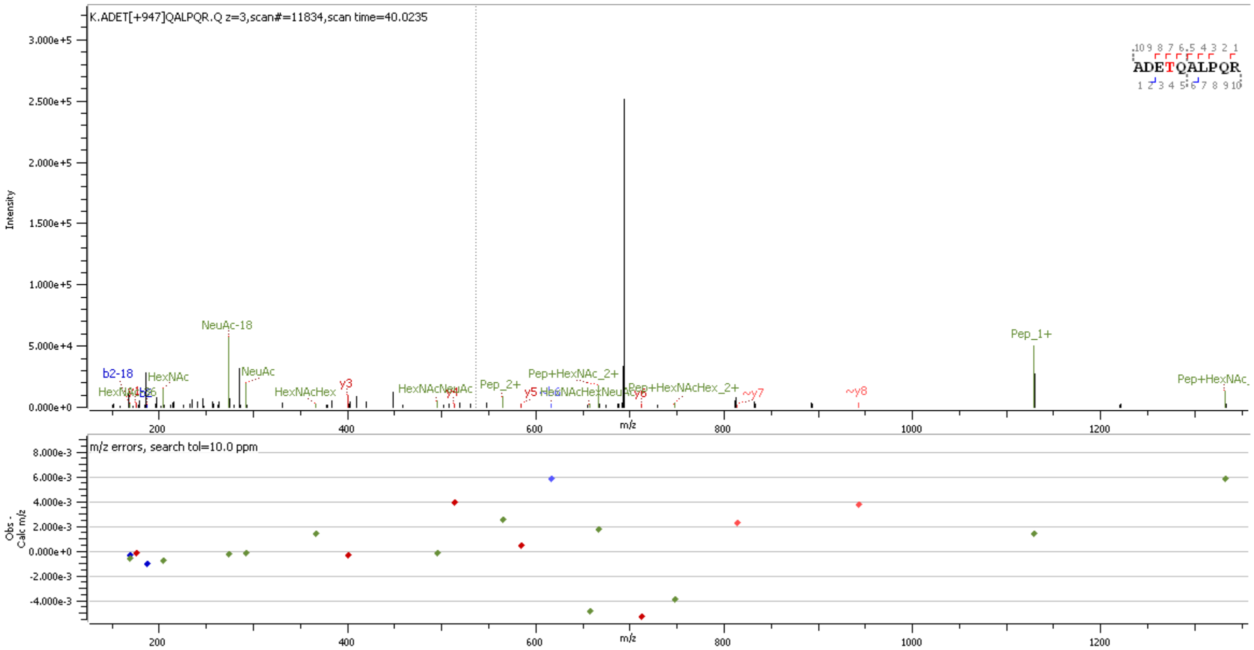

**Supp. Figure S88** HCD MS2 spectrum of O-glycopeptide [376]ADETQALPQR[385] containing HexNAc(1)Hex(1)NeuAc(2) at site T379

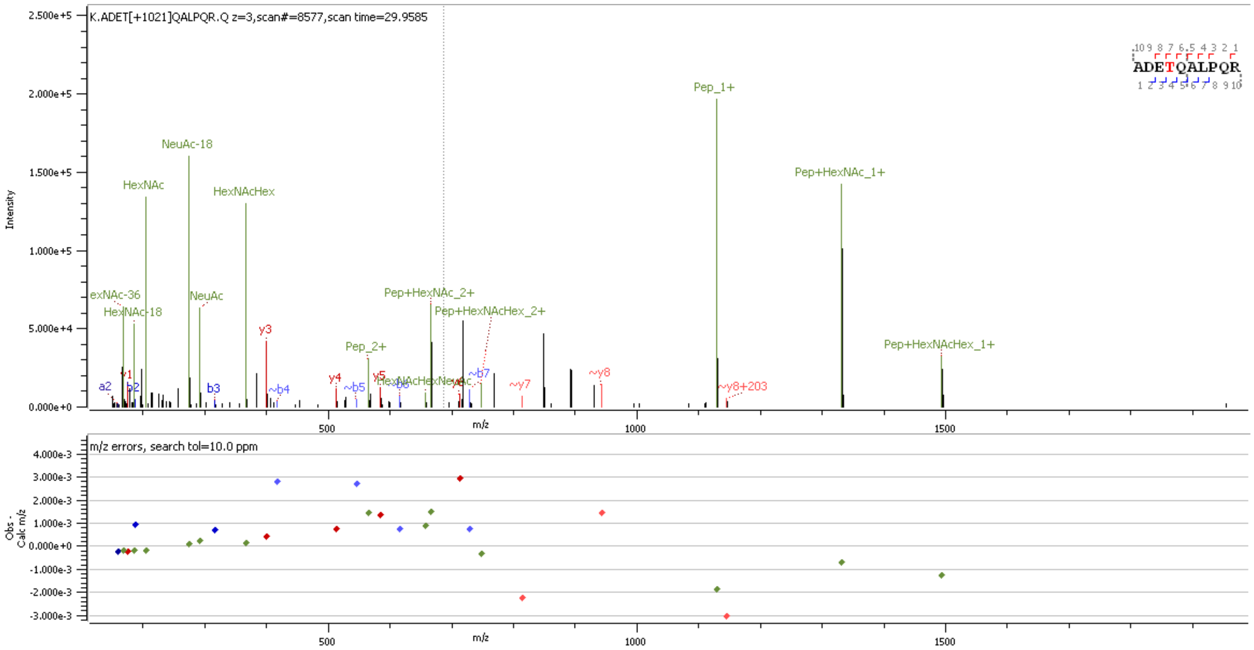


**Supp. Figure S89** HCD MS2 spectrum of O-glycopeptide [376]ADETQALPQR[385] containing HexNAc(2)Hex(2)NeuAc(1) at site T379

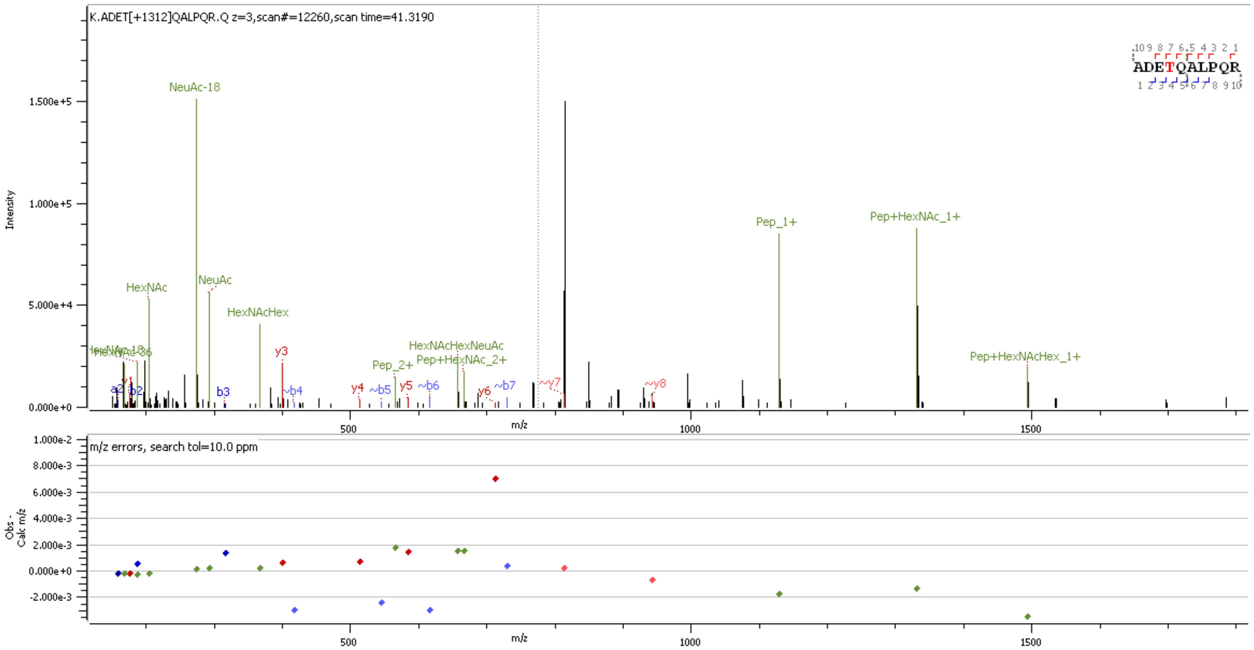


**Supp. Figure S90** HCD MS2 spectrum of O-glycopeptide [376]ADETQALPQR[385] containing HexNAc(2)Hex(2)NeuAc(2) at site T379

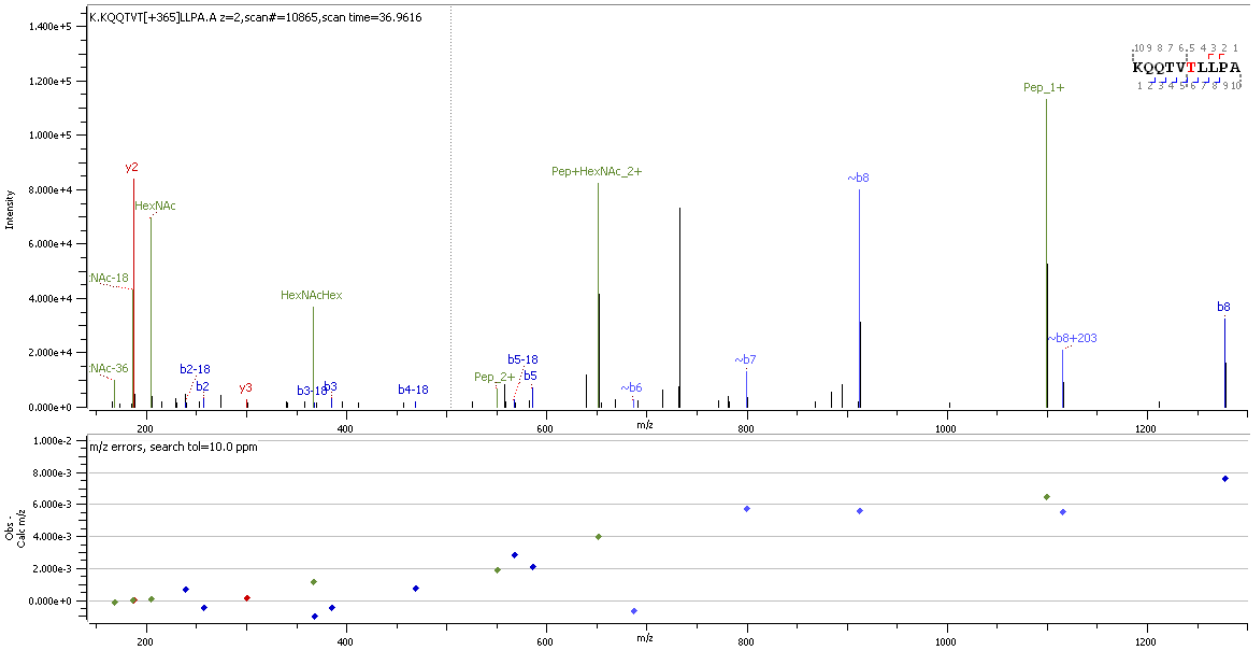


**Supp. Figure S91** HCD MS2 spectrum of O-glycopeptide $^{388}$KQQTVTLLPA$^{397}$ containing HexNAc(1)Hex(1) at site T391/393

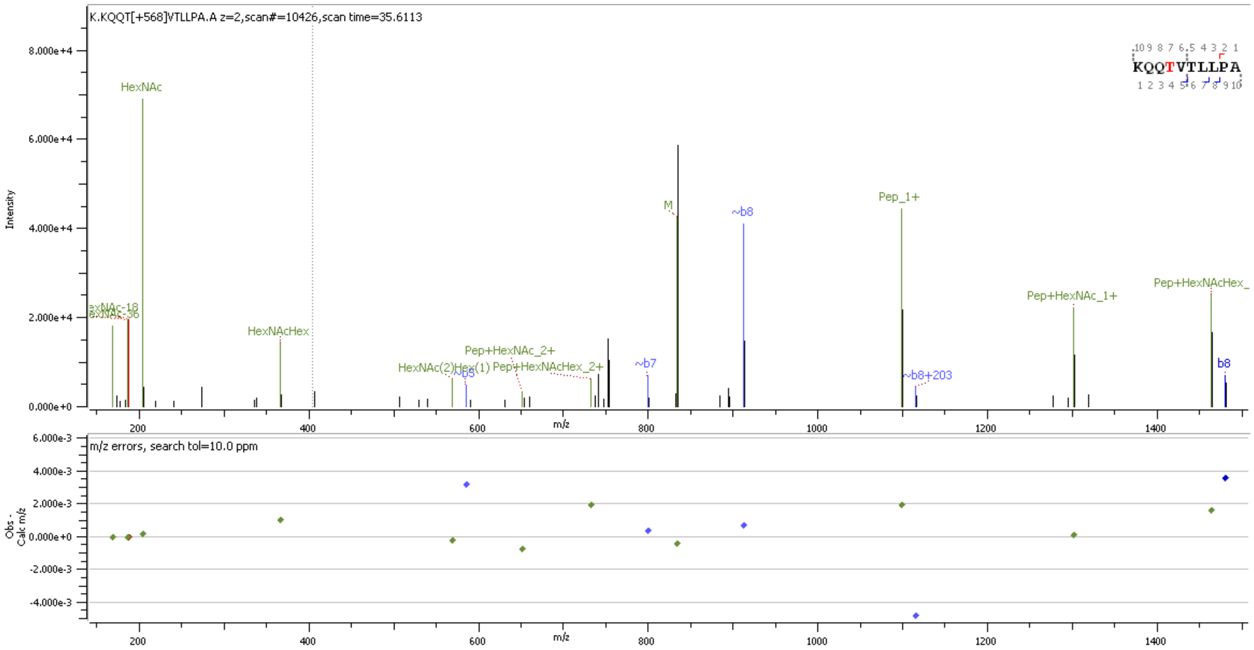


**Supp. Figure S92** HCD MS2 spectrum of O-glycopeptide $^{388}$KQQTVTLLPA$^{397}$ containing HexNAc(2)Hex(1) at site T391/393

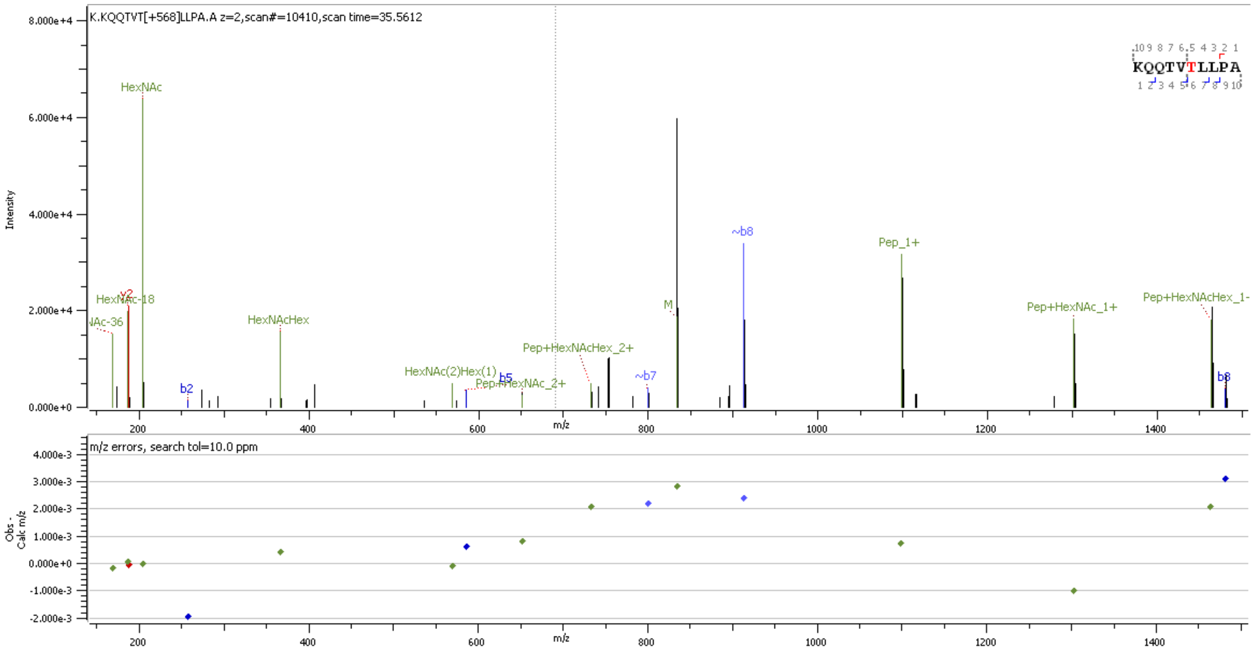


**Supp. Figure S93** HCD MS2 spectrum of O-glycopeptide [388]KQQTVTLLPA[397] containing HexNAc(2)Hex(1) at site T391/393

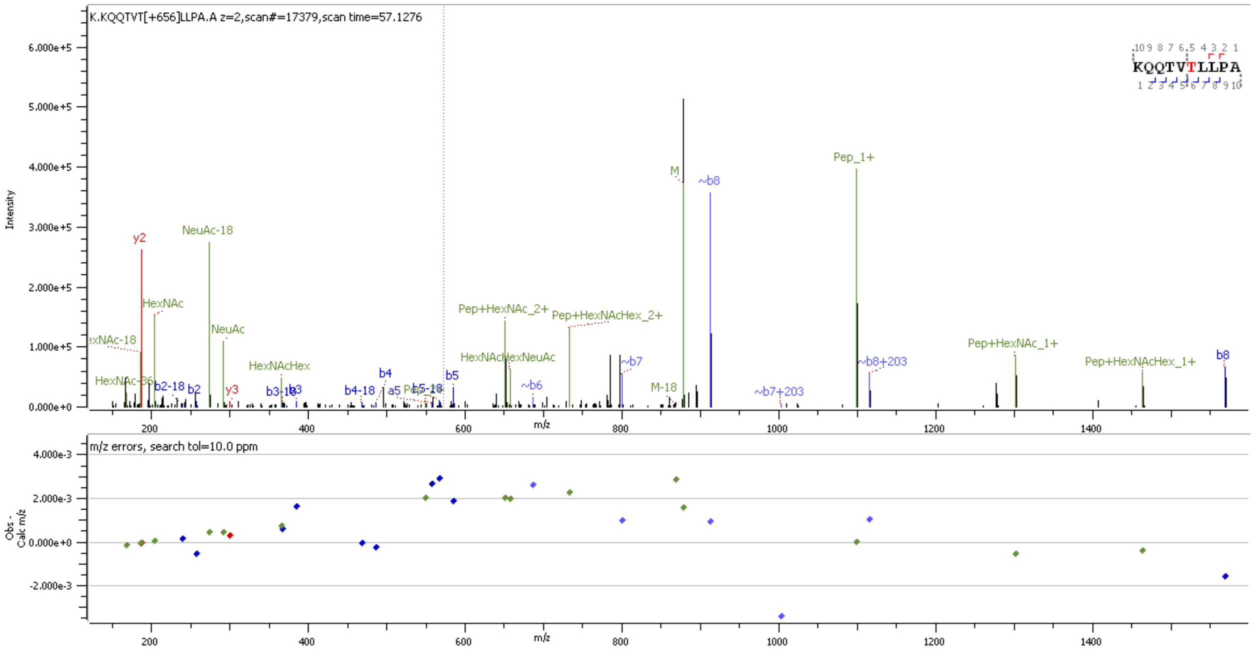


**Supp. Figure S94** HCD MS2 spectrum of O-glycopeptide [388]KQQTVTLLPA[397] containing HexNAc(1)Hex(1)NeuAc(1) at site T391/393

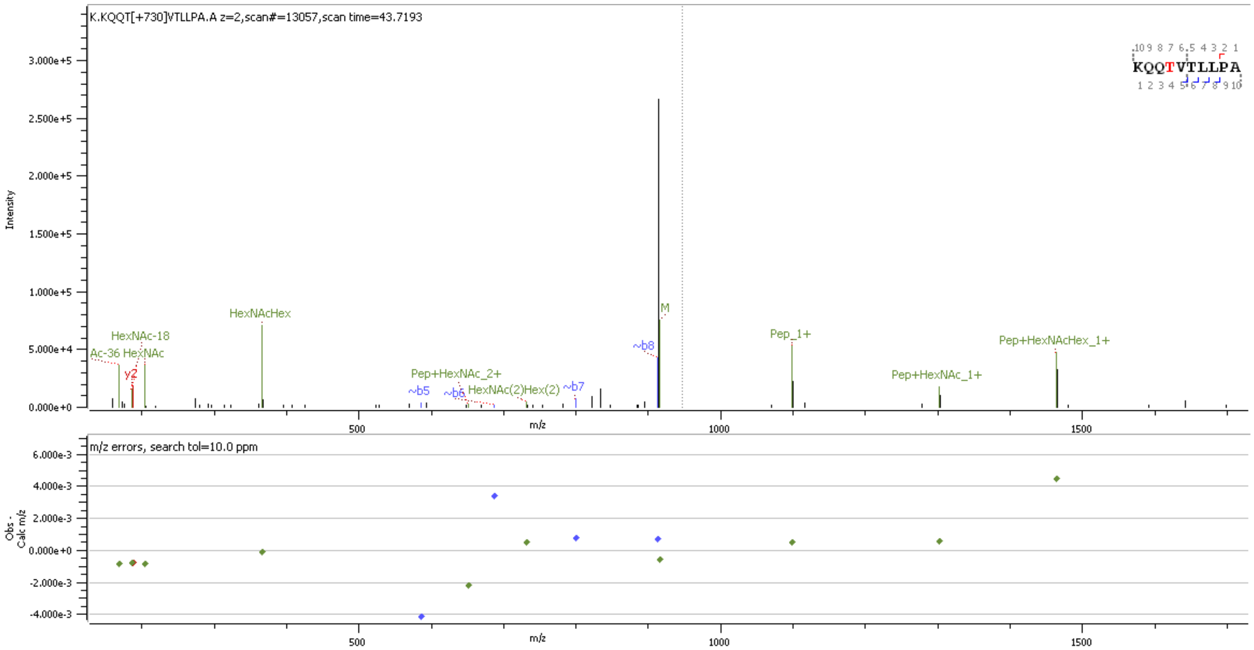


**Supp. Figure S95** HCD MS2 spectrum of O-glycopeptide $^{388}$KQQTVTLLPA$^{397}$ containing HexNAc(2)Hex(2) at site T391/393

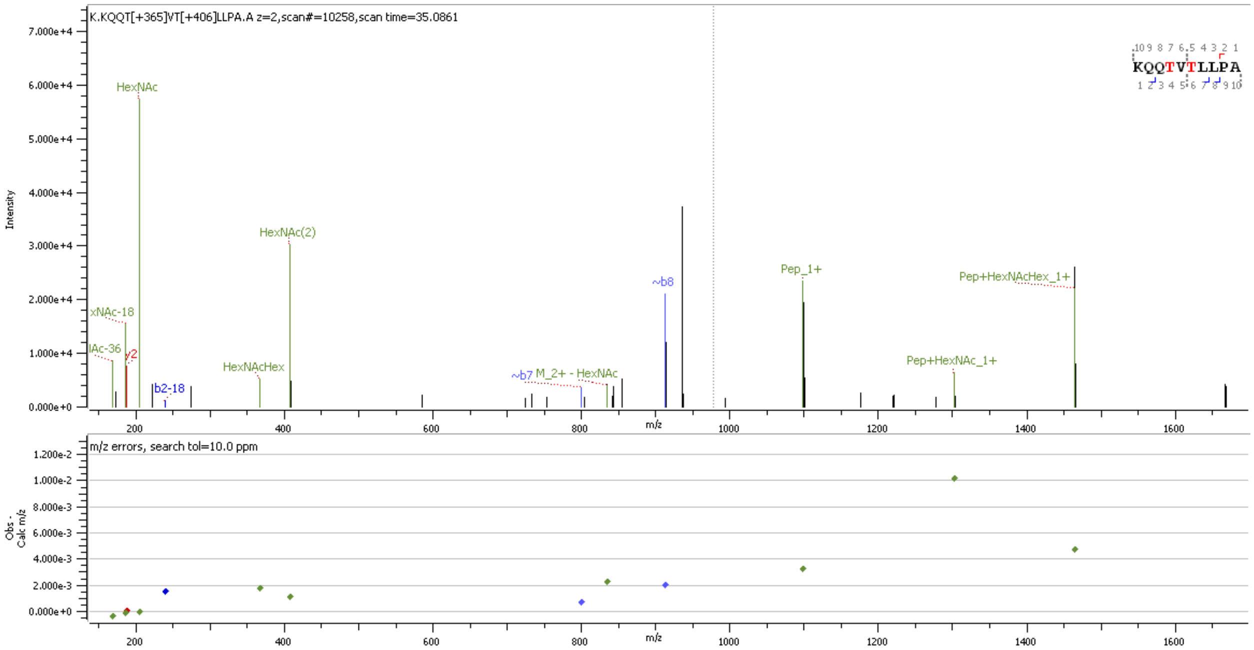


**Supp. Figure S96** HCD MS2 spectrum of O-glycopeptide $^{388}$KQQTVTLLPA$^{397}$ containing HexNAc(1)Hex(1); HexNAc(2) at site T391/393

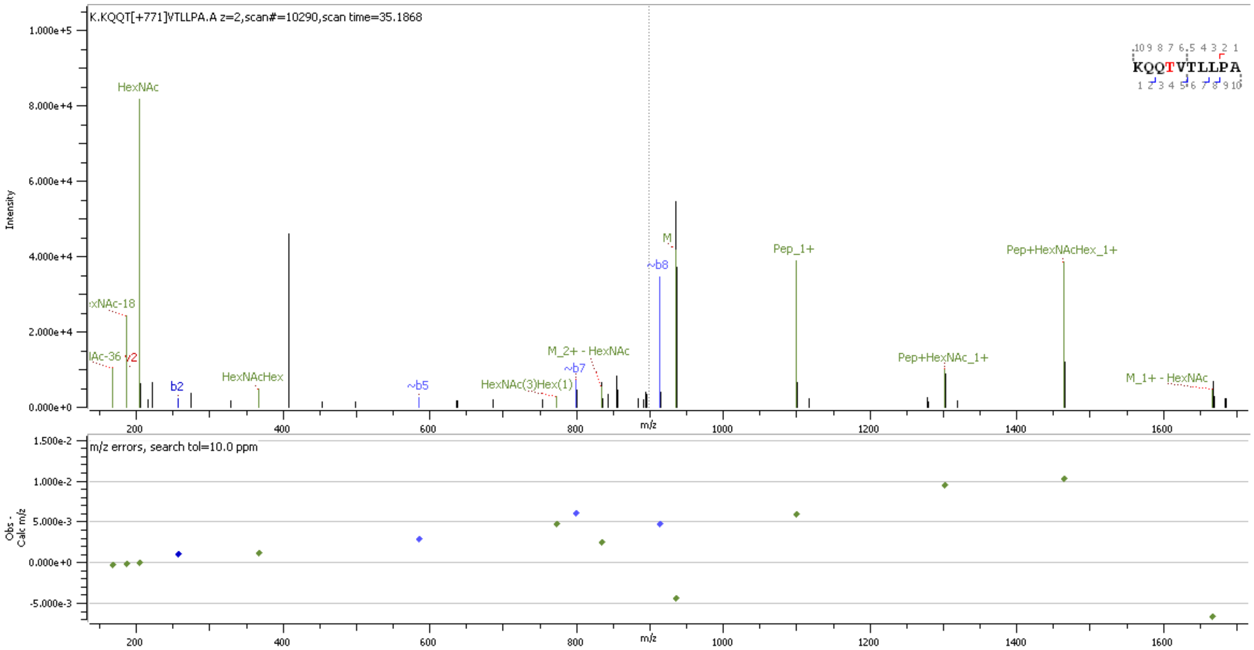


**Supp. Figure S97** HCD MS2 spectrum of O-glycopeptide [388]KQQTVTLLPA[397] containing HexNAc(3)Hex(1) at site T391/393

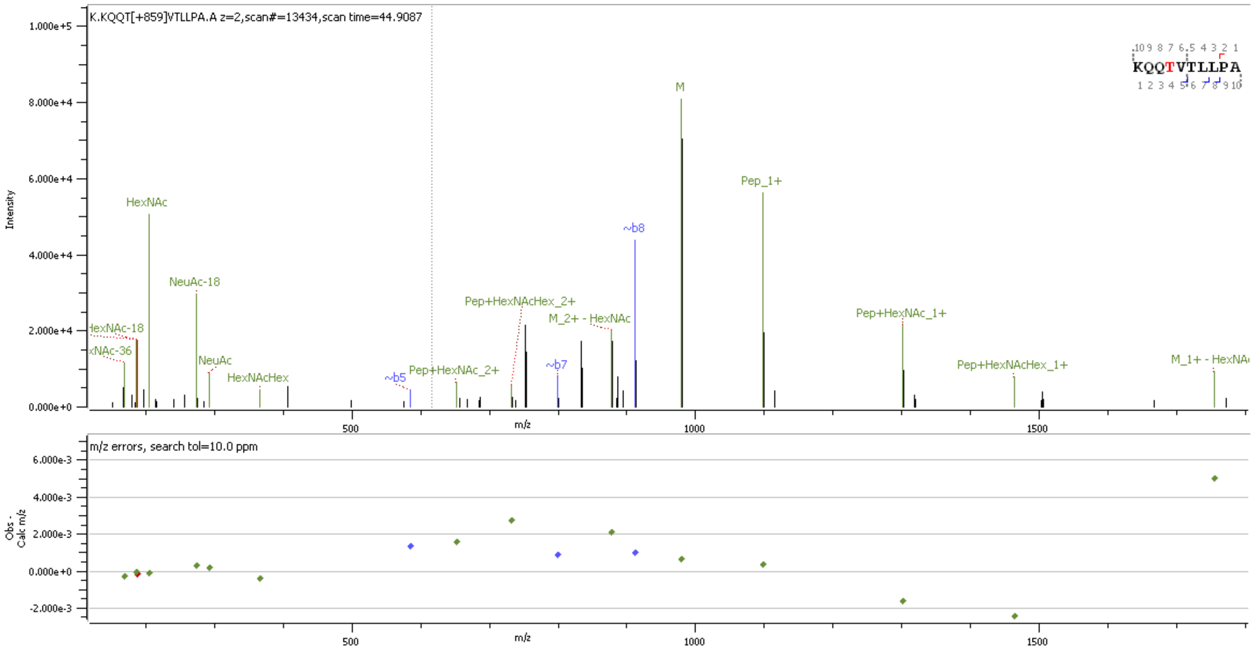


**Supp. Figure S98** HCD MS2 spectrum of O-glycopeptide [388]KQQTVTLLPA[397] containing HexNAc(2)Hex(1)NeuAc(1) at site T391/393

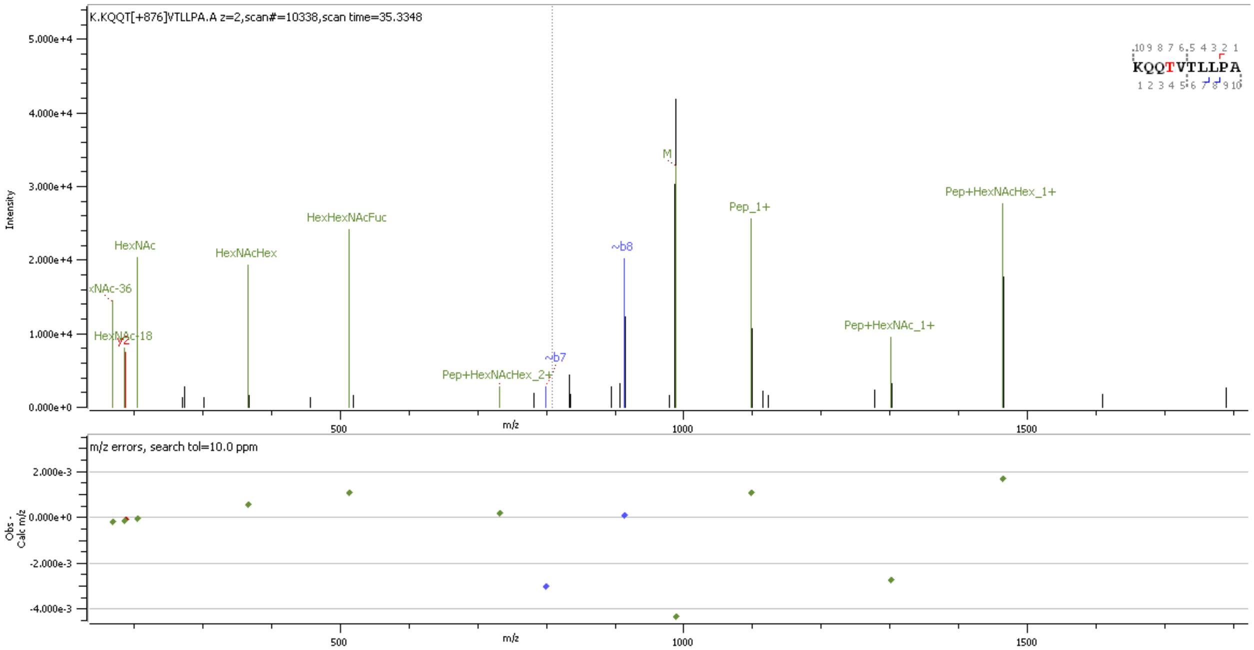


**Supp. Figure S99** HCD MS2 spectrum of O-glycopeptide [388]KQQTVTLLPA[397] containing HexNAc(2)Hex(2)Fuc(1) at site T391/393

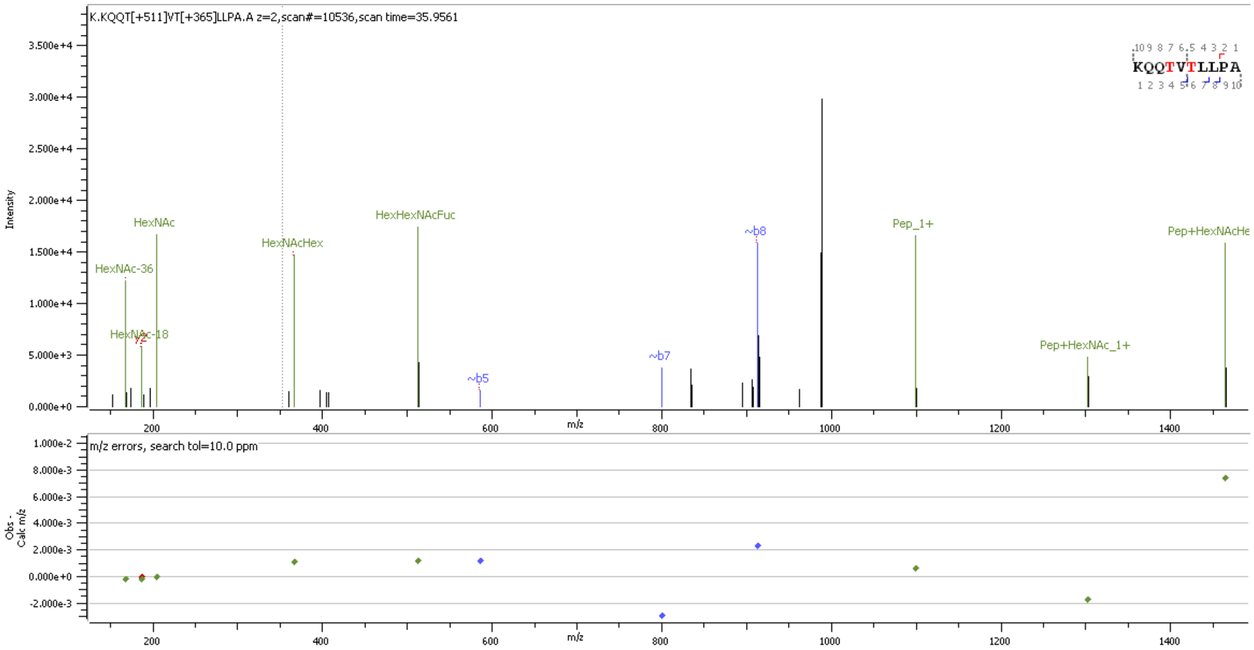


**Supp. Figure S100** HCD MS2 spectrum of O-glycopeptide [388]KQQTVTLLPA[397] containing HexNAc(1)Hex(1)Fuc(1); HexNAc(1)Hex(1) at site T391/393

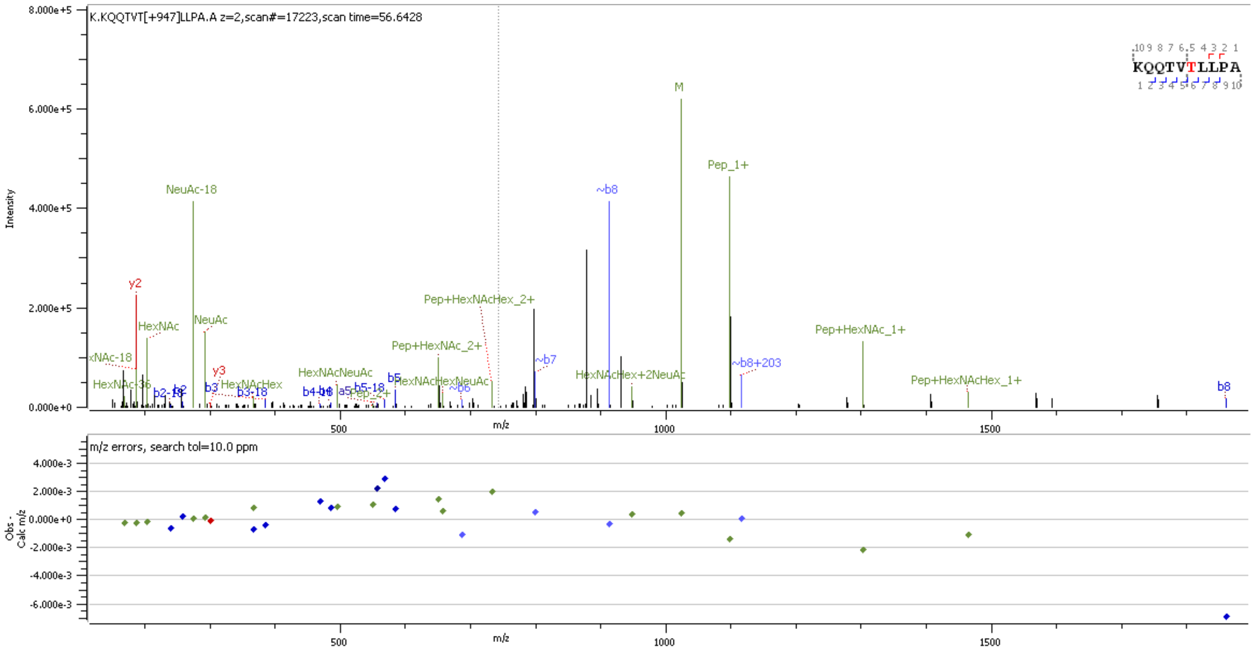


**Supp. Figure S101** HCD MS2 spectrum of O-glycopeptide $^{388}$KQQTVTLLPA$^{397}$ containing HexNAc(1)Hex(1)NeuAc(2) at site T391/393

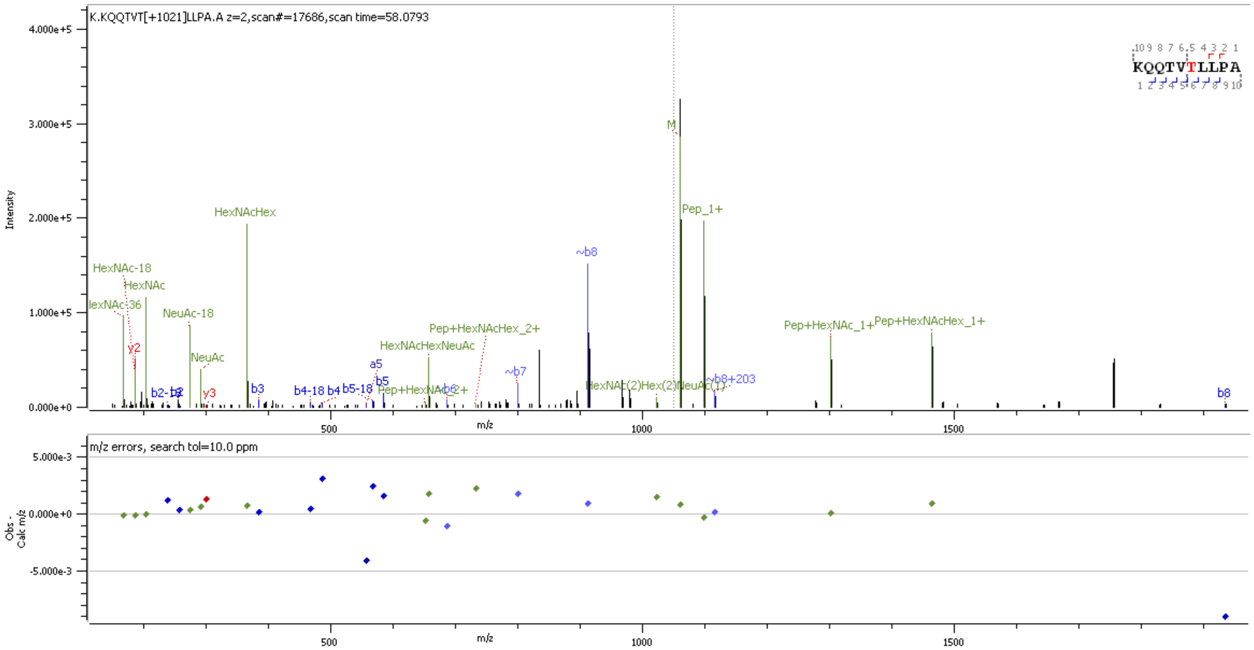


**Supp. Figure S102** HCD MS2 spectrum of O-glycopeptide $^{388}$KQQTVTLLPA$^{397}$ containing HexNAc(2)Hex(2)NeuAc(1) at site T391/393

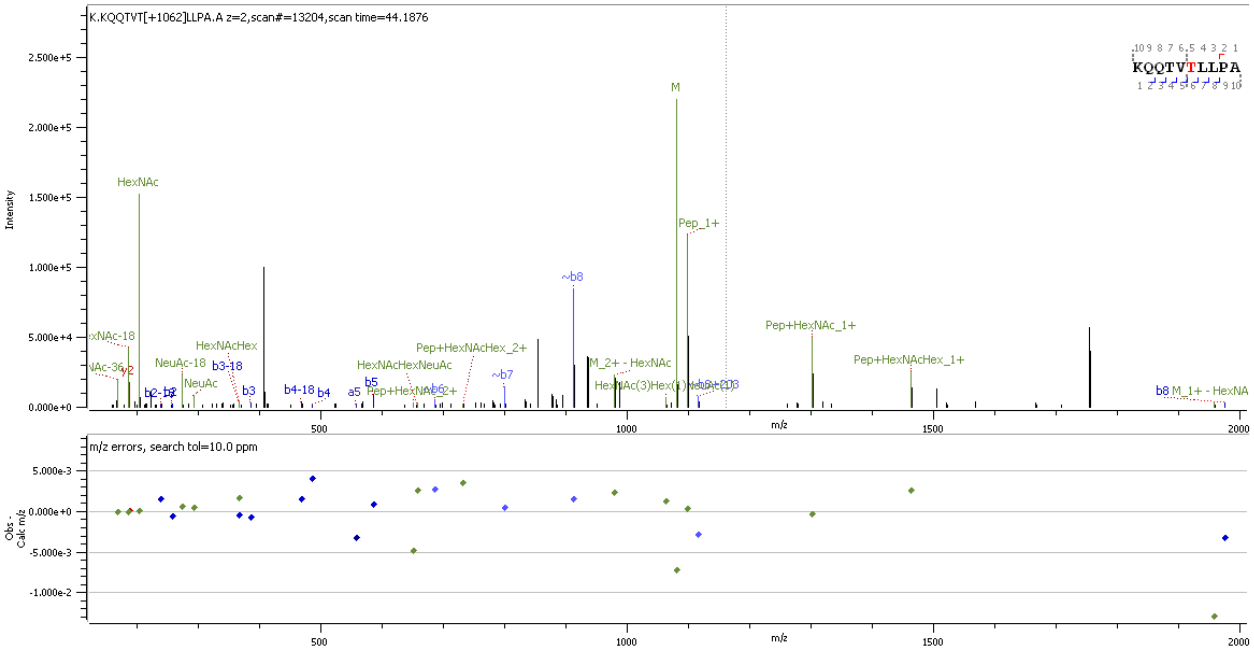


**Supp. Figure S103** HCD MS2 spectrum of O-glycopeptide $^{388}$KQQTVTLLPA$^{397}$ containing HexNAc(3)Hex(1)NeuAc(1) at site T391/393

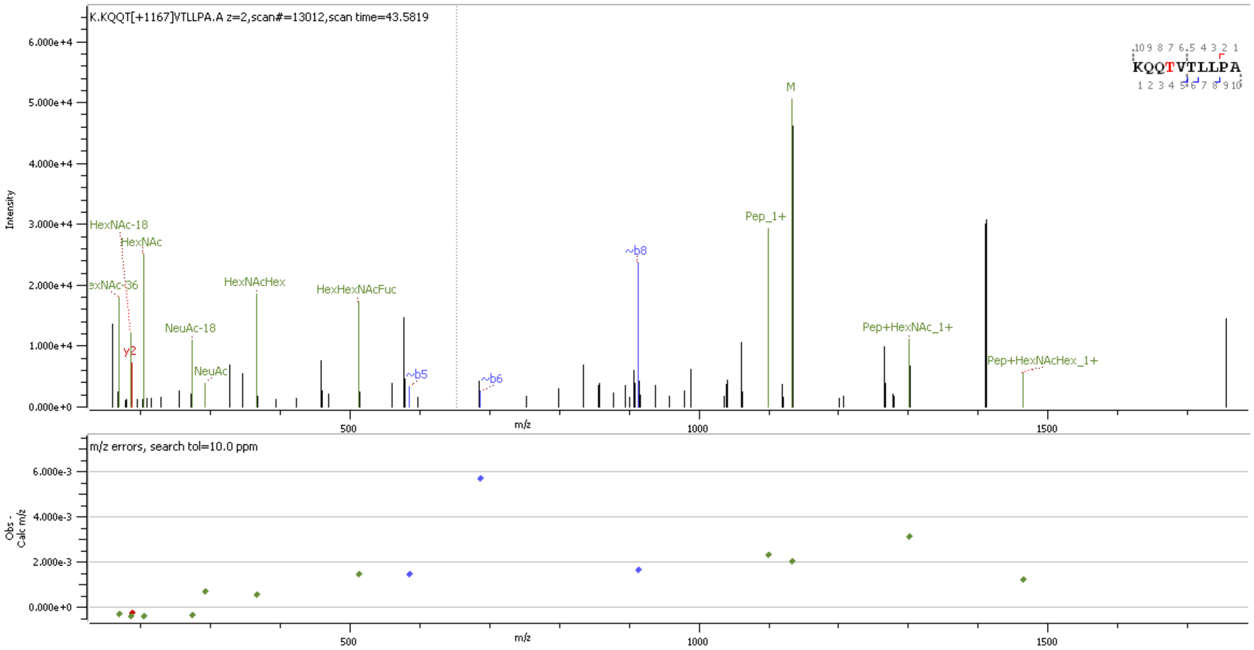


**Supp. Figure S104** HCD MS2 spectrum of O-glycopeptide $^{388}$KQQTVTLLPA$^{397}$ containing HexNAc(2)Hex(2)Fuc(1)NeuAc(1) at site T391/393

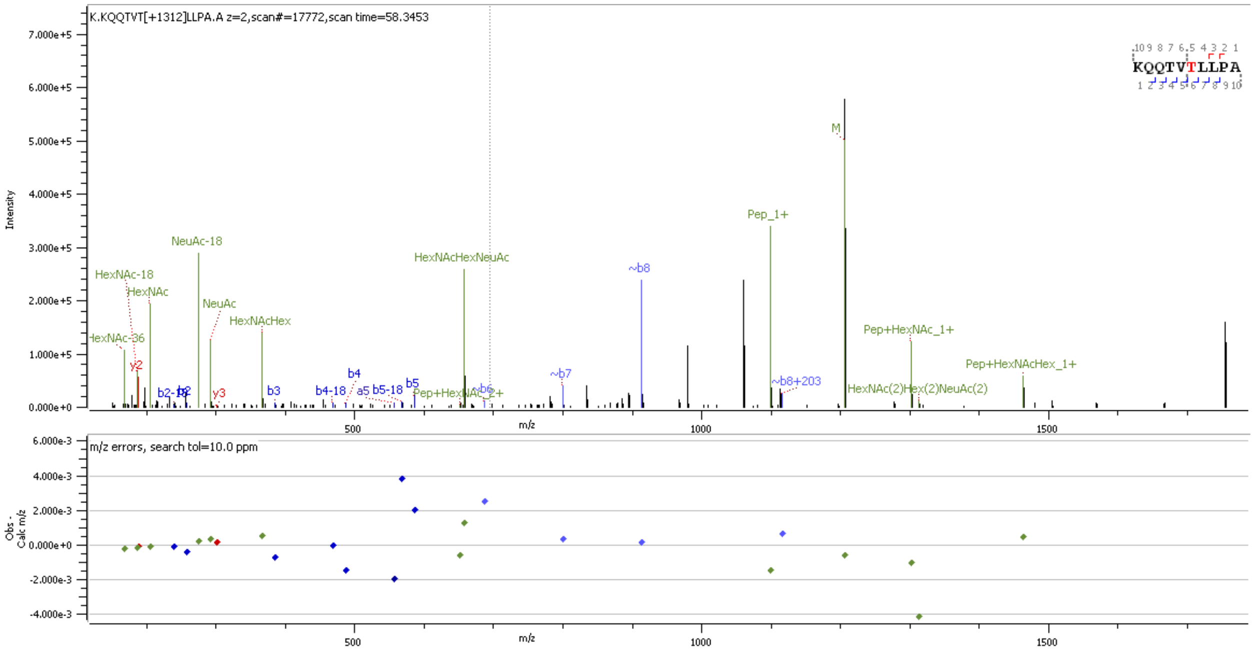


**Supp. Figure S105** HCD MS2 spectrum of O-glycopeptide $^{388}$KQQTVTLLPA$^{397}$ containing HexNAc(2)Hex(2)NeuAc(2) at site T391/393

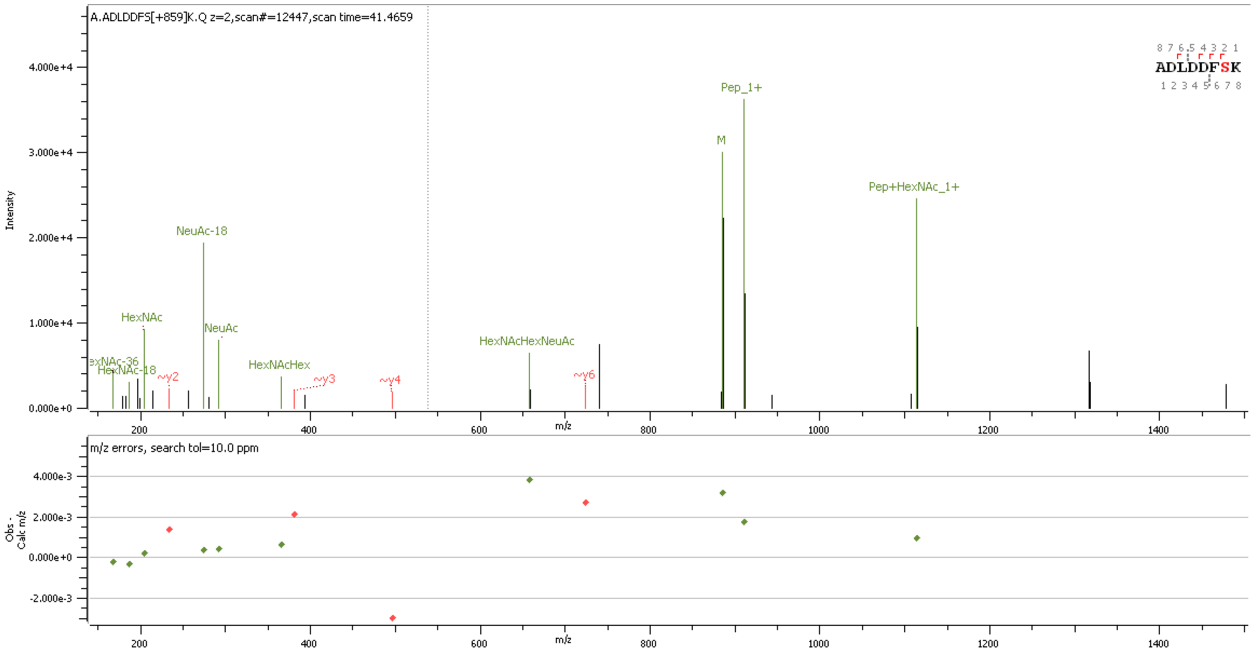


**Supp. Figure S106** HCD MS2 spectrum of O-glycopeptide $^{398}$ADLDDFSK$^{405}$ containing HexNAc(2)Hex(1)NeuAc(1) at site S404

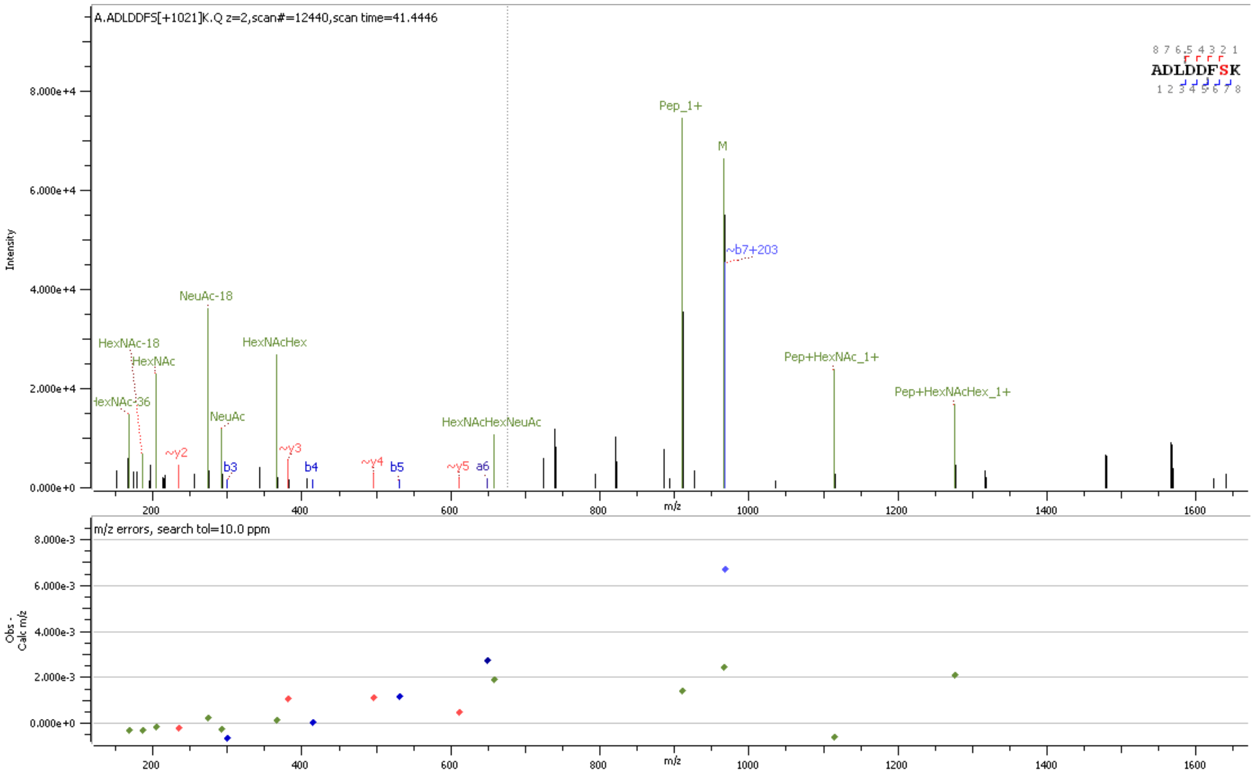


**Supp. Figure S107** HCD MS2 spectrum of O-glycopeptide [398]ADLDDFSK[405] containing HexNAc(2)Hex(2)NeuAc(1) at site S404

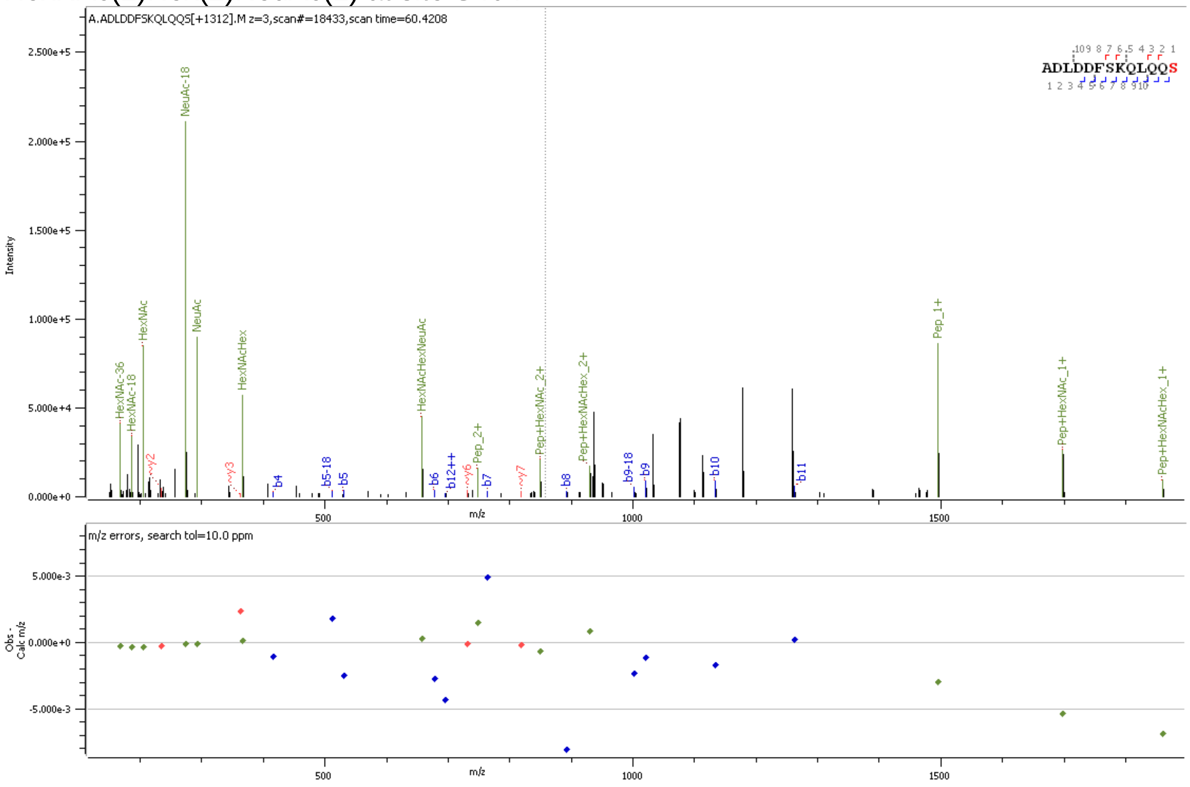


**Supp. Figure S108** HCD MS2 spectrum of O-glycopeptide [398]ADLDDFSKQLQQS[411] containing HexNAc(2)Hex(2)NeuAc(2) at site S404

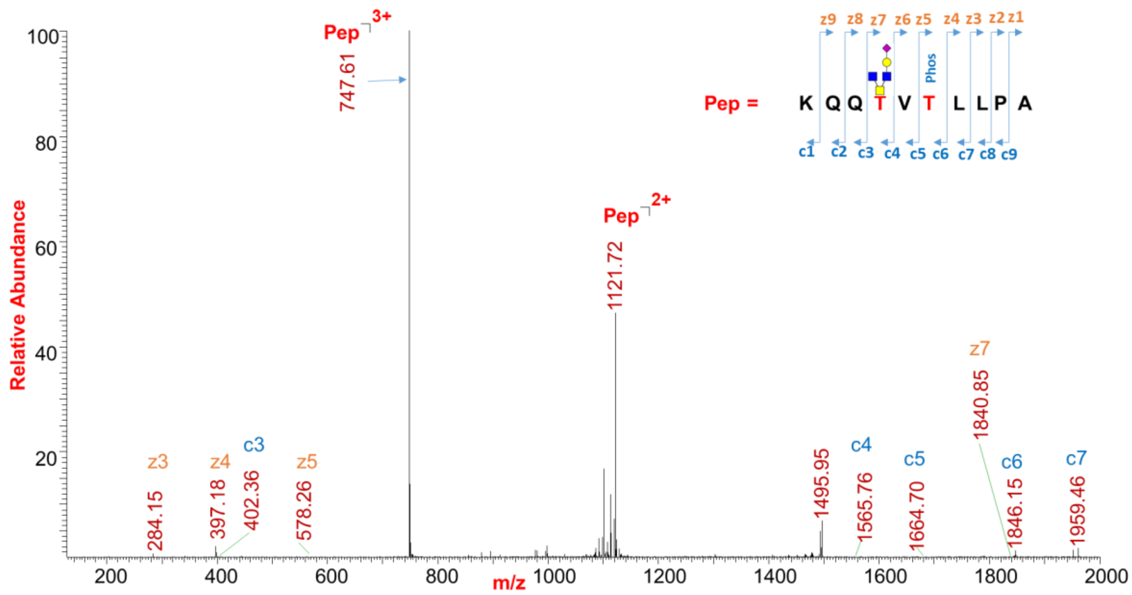


**Supp. Figure S109** ETD MS2 spectrum of O-glycopeptide [388]KQQTVTLLPA[397] containing HexNAc(3)Hex(1)NeuAc(1) at site T391 and phosphorylation at T393

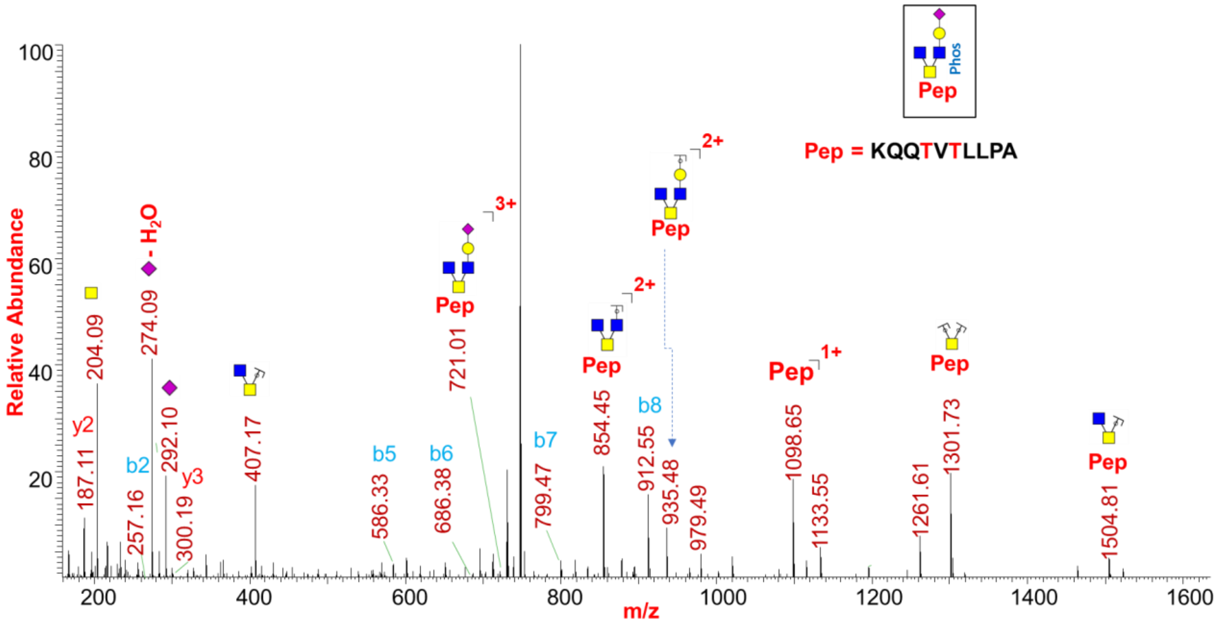


**Supp. Figure S110** HCD MS2 spectrum of O-glycopeptide [388]KQQTVTLLPA[397] containing HexNAc(3)Hex(1)NeuAc(1) at site T391 and phosphorylation at T393

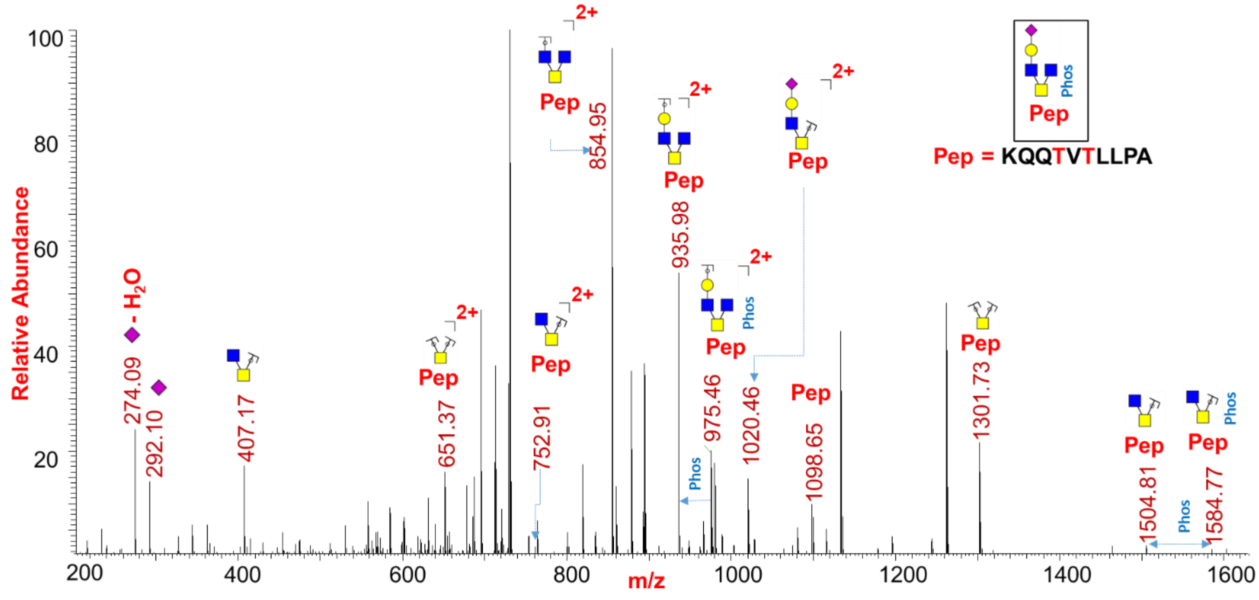


**Supp. Figure S111** CID MS2 spectrum of O-glycopeptide [388]KQQTVTLLPA[397] containing HexNAc(3)Hex(1)NeuAc(1) at site T391 and phosphorylation at T393

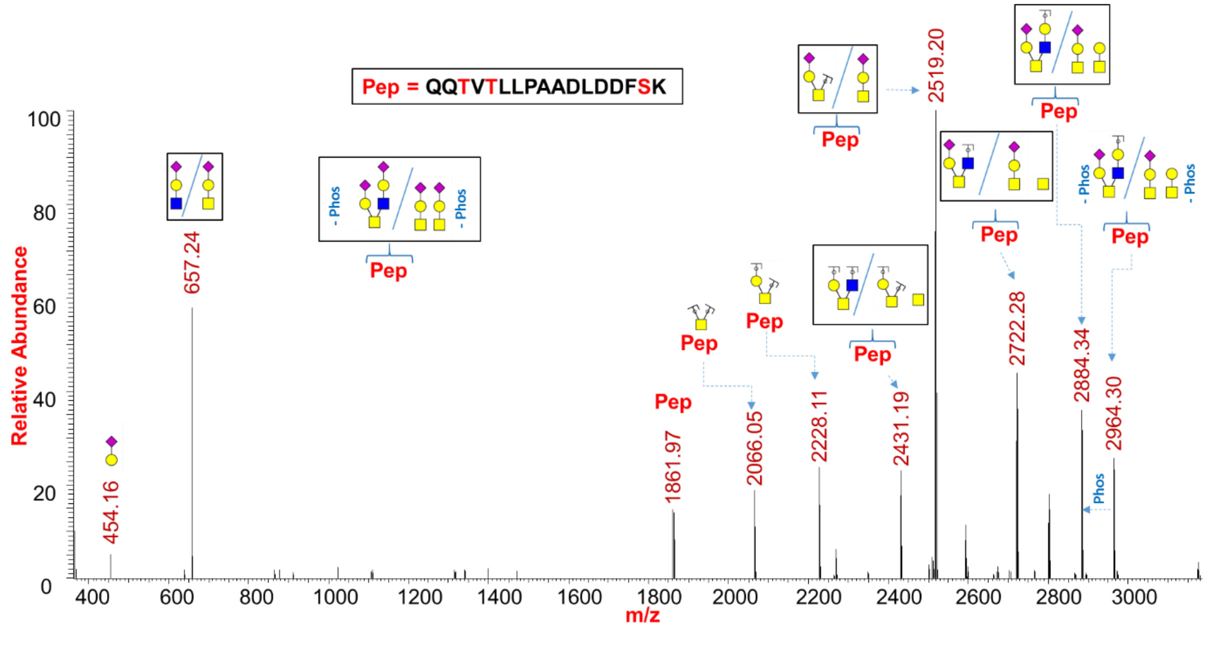


**Supp. Figure S112** CID MS2 spectrum of O-glycopeptide [389]QQTVTLLPAADLDDFSK[405] containing HexNAc(2)Hex(2)NeuAc(2) and phosphate at sites T391/T393/S404

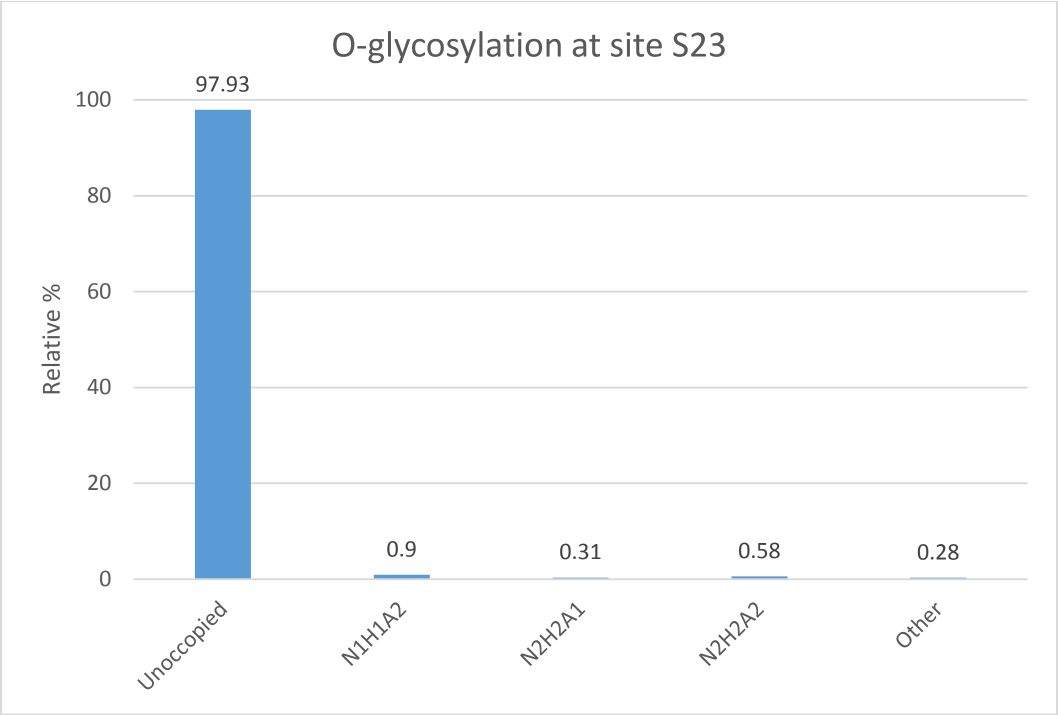


**Supp. Figure S113** Relative percentage of O-glycans at site S23

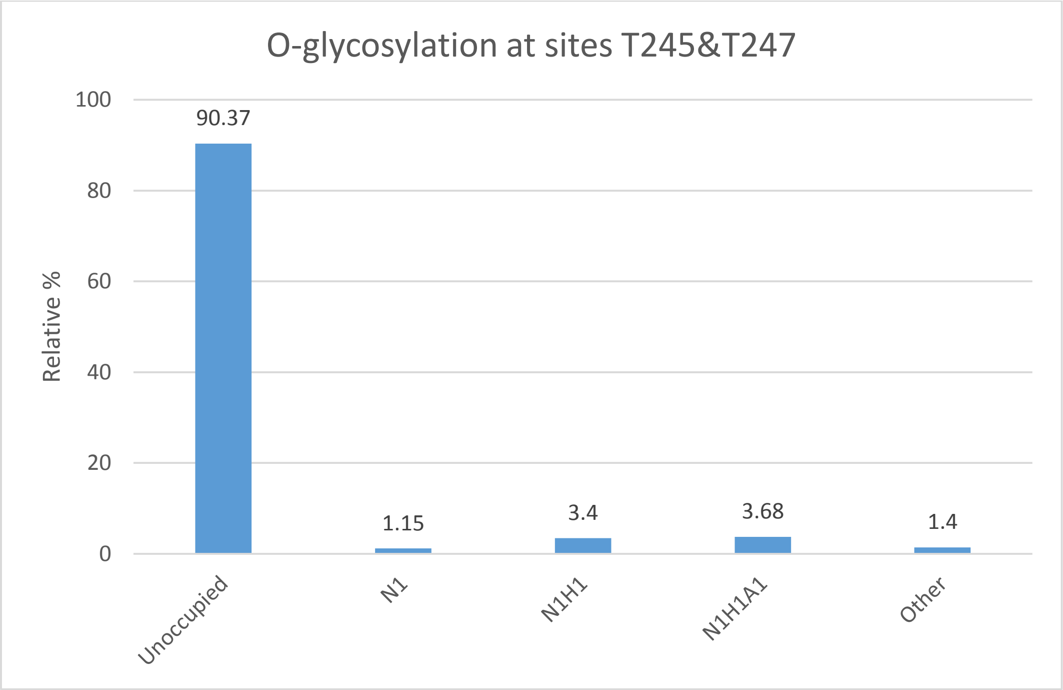


**Supp. Figure S114** Relative percentage of O-glycans at sites T245 & T247

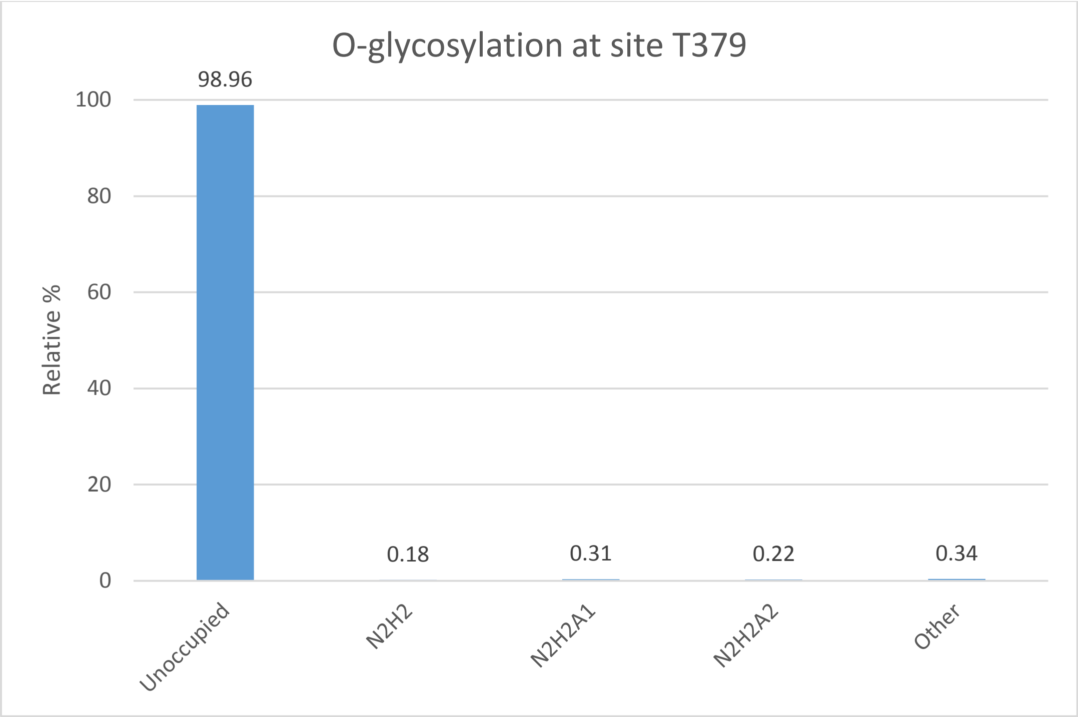


**Supp. Figure S115** Relative percentage of O-glycans at site T379
