## Supplemental Tables for "SARS-CoV-2 Nucleocapsid protein is decorated with multiple N- and O-glycans"

Supplemental Data Included: Figures S1 – S115, Supplemental Tables 1-4

Running Title: Glycosylation of Nucleocapsid (N) protein

| Table 1. Glycomic analysis of released and permethylated N-glycans via ESI-MS | | | |
|---|---|---|---|
| **Glycoform mass** | Relative Intensity | Δ mass (ppm) | Structure and Composition |
| **1579.78** | 23.1 | -2.3 | 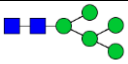 *(Hex)$_2$ + (Man)$_3$(GlcNAc)$_2$* |
| **1590.80** | 0.1 | 0.2 | 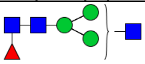 *(HexNAc)$_1$ (Deoxyhexose)$_1$ + (Man)$_3$(GlcNAc)$_2$* |
| **1620.81** | 0.2 | 0.2 | 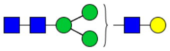 *(Hex)$_1$ (HexNAc)$_1$ + (Man)$_3$(GlcNAc)$_2$* |
| **1783.88** | 13.9 | -1.5 | 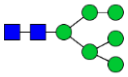 *(Hex)$_3$ + (Man)$_3$(GlcNAc)$_2$* |
| **1824.91** | 0.7 | 0.2 | 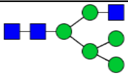 *(Hex)$_2$ (HexNAc)$_1$ + (Man)$_3$(GlcNAc)$_2$* |
| **1835.92** | 0.5 | -3.1 | 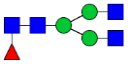 *(HexNAc)$_2$ (Deoxyhexose)$_1$ + (Man)$_3$(GlcNAc)$_2$* |
| **1865.94** | 1.0 | -3.0 | 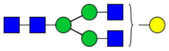 *(Hex)$_1$ (HexNAc)$_2$ + (Man)$_3$(GlcNAc)$_2$* |
| **1981.98** | 0.2 | -6.9 | 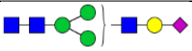 *(Hex)$_1$ (HexNAc)$_1$ (NeuAc)$_1$ + (Man)$_3$(GlcNAc)$_2$* |
| **1987.98** | 22.4 | -1.3 | 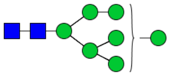 *(Hex)$_4$ + (Man)$_3$(GlcNAc)$_2$* |
| **2029.01** | 1.2 | 0.2 | 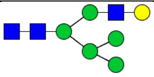 *(Hex)$_3$ (HexNAc)$_1$ + (Man)$_3$(GlcNAc)$_2$* |
| **2040.03** | 2.0 | -2.8 | 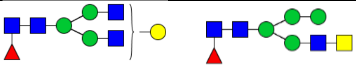 *(Hex)$_1$ (HexNAc)$_2$ (Deoxyhexose)$_1$ + (Man)$_3$(GlcNAc)$_2$* |
| **2070.04** | 2.1 | -2.7 | 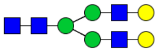 *(Hex)$_2$ (HexNAc)$_2$ + (Man)$_3$(GlcNAc)$_2$* |
| **2081.05** | 0.6 | -0.8 | 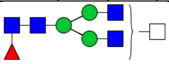 *(HexNAc)$_3$ (Deoxyhexose)$_1$ + (Man)$_3$(GlcNAc)$_2$* |
| **2111.06** | 0.2 | -1.3 | 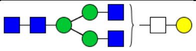 *(Hex)$_1$ (HexNAc)$_3$ + (Man)$_3$(GlcNAc)$_2$* |
| **2186.08** | 0.2 | -1.7 | 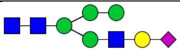 *(Hex)$_2$ (HexNAc)$_1$ (NeuAc)$_1$ + (Man)$_3$(GlcNAc)$_2$* |
| **2192.08** | 14.3 | -1.2 | 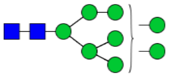 |

| | | | |
|---|---|---|---|
| | | | (Hex)$_5$ + (Man)$_3$(GlcNAc)$_2$ |
| **2214.11** | 1.9 | -2.1 | 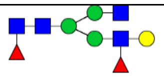<br>(Hex)$_1$ (HexNAc)$_2$ (Deoxyhexose)$_2$ + (Man)$_3$(GlcNAc)$_2$ |
| **2244.12** | 1.3 | -2.1 | 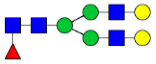<br>(Hex)$_2$ (HexNAc)$_2$ (Deoxyhexose)$_1$ + (Man)$_3$(GlcNAc)$_2$ |
| **2285.15** | 1.5 | -0.7 | 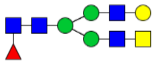<br>(Hex)$_1$ (HexNAc)$_3$ (Deoxyhexose)$_1$ + (Man)$_3$(GlcNAc)$_2$ |
| **2326.18** | 0.3 | 1.0 | 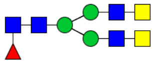<br>(HexNAc)$_4$ (Deoxyhexose)$_1$ + (Man)$_3$(GlcNAc)$_2$ |
| **2390.18** | 0.1 | -1.1 | 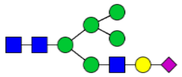<br>(Hex)$_3$ (HexNAc)$_1$ (NeuAc)$_1$ + (Man)$_3$(GlcNAc)$_2$ |
| **2396.18** | 0.4 | -1.1 | 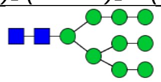<br>(Hex)$_6$ + (Man)$_3$(GlcNAc)$_2$ |
| **2401.20** | 0.7 | 0.6 | 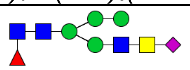<br>(Hex)$_1$ (HexNAc)$_2$ (Deoxyhexose)$_1$ (NeuAc)$_1$ + (Man)$_3$(GlcNAc)$_2$ |
| **2418.21** | 0.8 | -1.9 | 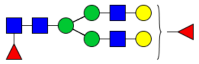<br>(Hex)$_2$ (HexNAc)$_2$ (Deoxyhexose)$_2$ + (Man)$_3$(GlcNAc)$_2$ |
| **2459.24** | 1.4 | -0.3 | 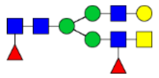 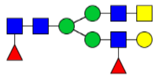<br>(Hex)$_1$ (HexNAc)$_3$ (Deoxyhexose)$_2$ + (Man)$_3$(GlcNAc)$_2$ |
| **2500.27** | 0.6 | 0.9 | 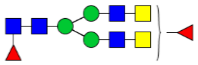<br>(HexNAc)$_4$ (Deoxyhexose)$_2$ + (Man)$_3$(GlcNAc)$_2$ |
| **2605.30** | 2.0 | 0.5 | 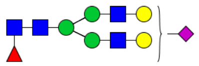<br>(Hex)$_2$ (HexNAc)$_2$ (Deoxyhexose)$_1$ (NeuAc)$_1$ + (Man)$_3$(GlcNAc)$_2$ |
| **2633.33** | 0.1 | 0.1 | 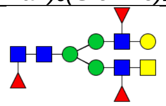<br>(Hex)$_1$ (HexNAc)$_3$ (Deoxyhexose)$_3$ + (Man)$_3$(GlcNAc)$_2$ |
| **2646.33** | 2.2 | -2.1 | 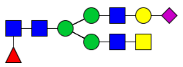 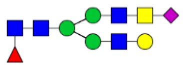<br>(Hex)$_1$ (HexNAc)$_3$ (Deoxyhexose)$_1$ (NeuAc)$_1$ + (Man)$_3$(GlcNAc)$_2$ |
| **2687.35** | 0.7 | -0.6 | 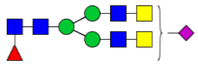<br>(HexNAc)$_4$ (Deoxyhexose)$_1$ (NeuAc)$_1$ + (Man)$_3$(GlcNAc)$_2$ |

| | | | |
|---|---|---|---|
| **2779.39** | 0.2 | 0.8 | 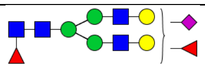 *(Hex)$_2$ (HexNAc)$_2$ (Deoxyhexose)$_2$ (NeuAc)$_1$ + (Man)$_3$(GlcNAc)$_2$* |
| **2820.41** | 0.7 | -1.7 | 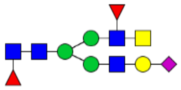 *(Hex)$_1$ (HexNAc)$_3$ (Deoxyhexose)$_2$ (NeuAc)$_1$ + (Man)$_3$(GlcNAc)$_2$* |
| **2966.47** | 0.9 | -0.9 | 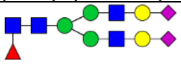 *(Hex)$_2$ (HexNAc)$_2$ (Deoxyhexose)$_1$ (NeuAc)$_2$ + (Man)$_3$(GlcNAc)$_2$* |
| **3007.50** | 1.2 | 3.8 | 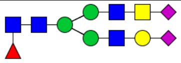 *(Hex)$_1$ (HexNAc)$_3$ (Deoxyhexose)$_1$ (NeuAc)$_2$ + (Man)$_3$(GlcNAc)$_2$* |

**Table 2. Glycomic analysis of released and permethylated O-glycans via ESI-MS**

| **Glycoform mass** | Relative Intensity | Δ mass (ppm) | Structure and Composition |
|---|---|---|---|
| **534.29** | 3.3 | 1.9 | 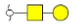 *(Hex)$_1$ (HexNAc)$_1$* |
| **575.31** | 0.9 | -3.7 | 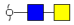 *(Hex)$_1$ (HexNAc)$_1$* |
| **895.46** | 11.2 | -3.3 | 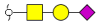 *(Hex)$_1$ (HexNAc)$_1$ (NeuAc)$_1$* |
| **983.51** | 3.2 | -5.1 | 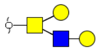 *(Hex)$_2$ (HexNAc)$_2$* |
| **1024.54** | 0.4 | -1.9 | 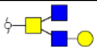 *(Hex)$_1$ (HexNAc)$_3$* |
| **1256.64** | 38.7 | -5.5 | 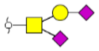 *(Hex)$_1$ (HexNAc)$_1$ (NeuAc)$_2$* |
| **1344.69** | 10.8 | 0.8 | 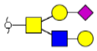 *(Hex)$_2$ (HexNAc)$_2$ (NeuAc)$_1$* |
| **1385.71** | 2.5 | -3.6 | 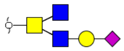 *(Hex)$_1$ (HexNAc)$_3$ (NeuAc)$_1$* |
| **1518.77** | 2.2 | -5.2 | 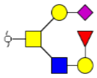 *(Hex)$_2$ (HexNAc)$_2$ (Deoxyhexose)$_1$ (NeuAc)$_1$* |
| **1705.86** | 26.8 | -1.7 | 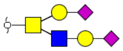 *(Hex)$_2$ (HexNAc)$_2$ (NeuAc)$_2$* |

| Table 3. N-glycopeptide Analysis | | | | | |
|---|---|---|---|---|---|
| **N-glycan site** | **Peptide Sequence** | **Composition** | **Proposed Structures** | **Glycoform Mass** | **Rel. %** |
| N47 | $^{41}$RPQGLPNNTASWF$^{53}$ | Unoccupied | Unoccupied | 1487.7390 | 46.24 |
| | | $(GlcNAc)_2$+ $(Man)_3(GlcNAc)_2(Fuc)_1$ | 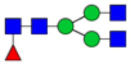 | 2933.2758 | 0.27 |
| | | $(Gal)_1(GlcNAc)_2$+ $(Man)_3(GlcNAc)_2(Fuc)_1$ | 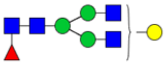 | 3095.3272 | 1.40 |
| | | $(Gal)_1(GlcNAc)_2(Fuc)_1$+ $(Man)_3(GlcNAc)_2(Fuc)_1$ | 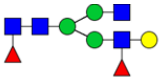 | 3241.3849 | 2.60 |
| | | $(Gal)_2(GlcNAc)_2(Fuc)_1$+ $(Man)_3(GlcNAc)_2(Fuc)_1$ | 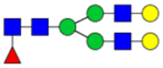 | 3257.3840 | 1.87 |
| | | $(GalNAc)_1(Gal)_1(GlcNAc)_2$+ $(Man)_3(GlcNAc)_2(Fuc)_1$ | 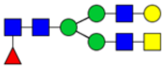 | 3298.4066 | 4.55 |
| | | $(GalNAc)_2(GlcNAc)_2$+ $(Man)_3(GlcNAc)_2(Fuc)_1$ | 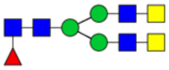 | 3339.4332 | 3.31 |
| | | $(NeuAc)_1(Gal)_1(GlcNAc)_2$+ $(Man)_3(GlcNAc)_2(Fuc)_1$ | 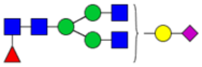 | 3386.4243 | 1.37 |
| | | $(Gal)_2(GlcNAc)_2(Fuc)_1$+ $(Man)_3(GlcNAc)_2(Fuc)_1$ | 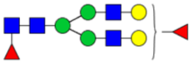 | 3404.4422 | 2.15 |
| | | $(GalNAc)_1(Gal)_1(GlcNAc)_2(Fuc)_1$ + $(Man)_3(GlcNAc)_2(Fuc)_1$ | 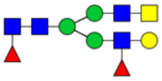 | 3445.4692 | 4.50 |
| | | $(GalNAc)_2(GlcNAc)_2(Fuc)_1$+ $(Man)_3(GlcNAc)_2(Fuc)_1$ | 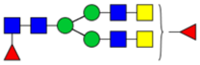 | 3486.4958 | 4.86 |
| | | $(NeuAc)_1(Gal)_2(GlcNAc)_2$+ $(Man)_3(GlcNAc)_2(Fuc)_1$ | 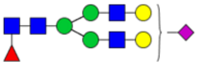 | 3549.4813 | 5.05 |
| | | $(NeuAc)_1(GalNAc)_1(Gal)_1$ $(GlcNAc)_2$+ $(Man)_3(GlcNAc)_2(Fuc)_1$ | 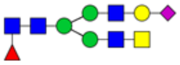 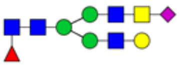 | 3590.5063 | 6.56 |
| | | $(GalNAc)_1$ $(Gal)_1(GlcNAc)_2(Fuc)_2$+ $(Man)_3(GlcNAc)_2(Fuc)_1$ | 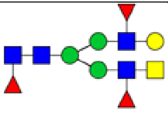 | 3591.5266 | 0.45 |

| | | | | | |
|---|---|---|---|---|---|
| | | $(NeuAc)_1$ $(GalNAc)_2(GlcNAc)_2$+ $(Man)_3(GlcNAc)_2(Fuc)_1$ | 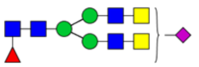 | 3631.5333 | 5.47 |
| | | $(NeuAc)_1(Gal)_1$ $(GalNAc)_1(GlcNAc)_2(Fuc)_1$+ $(Man)_3(GlcNAc)_2(Fuc)_1$ | 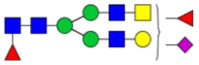 | 3736.5663 | 2.19 |
| | | $(NeuAc)_2(Gal)_2(GlcNAc)_2$+ $(Man)_3(GlcNAc)_2(Fuc)_1$ | 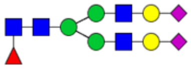 | 3840.5779 | 2.51 |
| | | $(NeuAc)_2(GalNAc)_1$ $(Gal)_1(GlcNAc)_2$+ $(Man)_3(GlcNAc)_2(Fuc)_1$ | 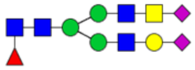 | 3881.6039 | 4.65 |
| N269 | $^{267}$AYNVTQAFGR$^{276}$ | Unoccupied | | 1126.5639 | 6.05 |
| | | $(Man)_4(GlcNAc)_2$ | 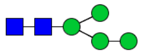 | 2181.9379 | 2.16 |
| | | $(Man)_5(GlcNAc)_2$ | 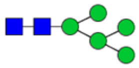 | 2343.9895 | 40.68 |
| | | ( $(Man)_6(GlcNAc)_2$ | 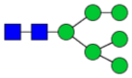 | 2506.0421 | 14.65 |
| | | $(Man)_2(GlcNAc)$+ $(Man)_3(GlcNAc)_2$ | 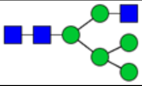 | 2547.0692 | 1.09 |
| | | (Gal) $(GlcNAc)_2$+ $(Man)_3(GlcNAc)_2$ | 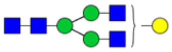 | 2588.0943 | 1.38 |
| | | $(Man)_7(GlcNAc)_2$ | 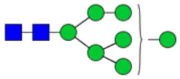 | 2668.0965 | 18.84 |
| | | $(Man)_2$(Gal) $(GlcNAc)_1$+ $(Man)_3(GlcNAc)_2$ | 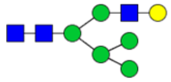 | 2709.1234 | 1.98 |
| | | $(Gal)_2$ $(GlcNAc)_2$+ $(Man)_3(GlcNAc)_2$ | 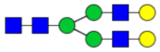 | 2750.1476 | 2.46 |
| | | $(Man)_8(GlcNAc)_2$ | 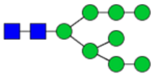 | 2830.1494 | 8.71 |
| | | $(Gal)_2(GlcNAc)_2$+ $(Man)_3(GlcNAc)_2(Fuc)_1$ | 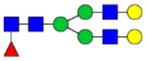 | 2896.2072 | 0.42 |
| | | $(GalNAc)_1(Gal)_1(GlcNAc)_2$+ $(Man)_3(GlcNAc)_2(Fuc)_1$ | 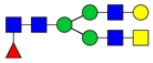 | 2937.2344 | 0.34 |

| | | | | | |
|---|---|---|---|---|---|
| | | $(Man)_9(GlcNAc)_2$ | 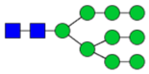 | 2992.2026 | 0.63 |
| | | $(Man)_2(NeuAc)_1(Gal)(GlcNAc)_1$+ $(Man)_3(GlcNAc)_2$ | 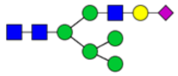 | 3000.2159 | 0.63 |

| **Table 3. O-glycopeptide Analysis** | | | | | |
|---|---|---|---|---|---|
| **O-glycan site** | **Peptide Sequence** | **Composition** | **Proposed Structures** | **Glycoform Mass** | **Rel. %** |
| S23 | $^{15}$ITFGGPSDSTGSNQN* GER$^{32}$ | Unoccopied | | 1824.7995 | 97.93 |
| | | HexNAc(1) | 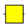 | 2027.8776 | 0.02 |
| | | HexNAc(1)Hex(1) | 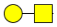 | 2189.9312<br>Full mass detected | 0.04 |
| | | HexNAc(1)Hex(1)NeuAc(1) | 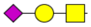 | 2481.0298<br>Full mass detected | 0.20 |
| | | HexNAc(2)Hex(2) | 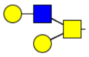 | 2555.0639<br>Full mass detected | 0.03 |
| | | HexNAc(1)Hex(1)NeuAc(2) | 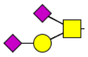 | 2771.1381 | 0.90 |
| | | HexNAc(2)Hex(2)NeuAc(1) | 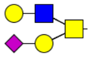 | 2846.1601 | 0.31 |
| | | HexNAc(2)Hex(2)NeuAc(2) | 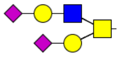 | 3137.2547 | 0.58 |
| T148 | DHIGTR | Unoccupied | | | 18.64 |

| | | | | | |
|---|---|---|---|---|---|
| | | HexNAc(1)Hex(1)NeuAc(1) | 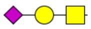 | 1354.5848 | 3.39 |
| | | HexNAc(2)Hex(2) | 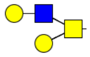 | 1428.6233 | 4.68 |
| | | HexNAc(1)Hex(1)NeuAc(2) | 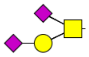 | 1645.6806 | 3.43 |
| | | HexNAc(2)Hex(2)NeuAc(1) | 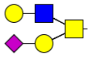 | 1719.7193 | 26.22 |
| | | HexNAc(2)Hex(2)NeuAc(2) | 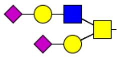 | 2010.8107 | 43.64 |
| T165 & T166 | LQLPQGTTLPK | Unoccupied | | | 0.03 |
| | | HexNAc(1) | 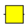 | 1398.7846 | 1.16 |
| | | HexNAc(1)Hex(1) | 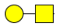 | 1560.8373 | 22.65 |
| | | HexNAc(2) | 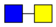 | 1601.8632 | 1.83 |
| | | HexNAc(2)Hex(1) | 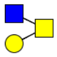 | 1763.9179 | 1.83 |
| | | HexNAc(1)Hex(1)NeuAc(1) | 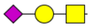 | 1851.9338 | 17.40 |
| | | HexNAc(2)Hex(2) | 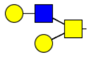 | 1925.9689 | 1.88 |
| | | HexNAc(3)Hex(1) | 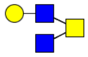 | 1966.9958 | 1.66 |
| | | HexNAc(2)Hex(1)NeuAc(1) | 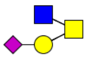 | 2055.0123 | 1.86 |

| | | | | | |
|---|---|---|---|---|---|
| | | HexNAc(1)Hex(1)NeuAc(2) | 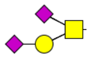 | 2144.0334 | 26.86 |
| | | HexNAc(2)Hex(2)NeuAc(1) | 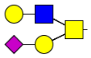 | 2217.0645 | 7.43 |
| | | HexNAc(3)Hex(1)NeuAc(1); | 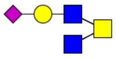 | 2258.1004 | 5.19 |
| | | HexNAc(2)Hex(2)Fuc(1)NeuAc(1) | 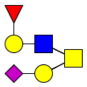 | 2363.1264 | 1.12 |
| | | HexNAc(2)Hex(2)NeuAc(2) | 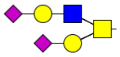 | 2508.1618 | 9.09 |
| T205 & S206 | [204]GTSPAR[209] | Unoccopied | | | 4.33 |
| | | HexNAc(1)Hex(1)NeuAc(1) | 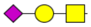 | 1244.5371<br>Full mass detected | 1.69 |
| | | HexNAc(1)Hex(1); HexNAc(1)Hex(1)<br>Or<br>HexNAc(2)Hex(2) | 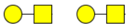<br>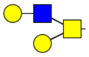 | 1318.5749 | 0.45 |
| | | HexNAc(2); HexNAc(1)Hex(1)<br>Or | 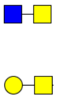 | 1359.6026 | 5.75 |

| | | | | | |
|---|---|---|---|---|---|
| | | HexNAc(3)Hex(1) | 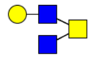 | | |
| | | HexNAc(1)Hex(1)NeuAc(2) | 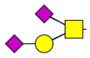 | 1535.6342 | 39.19 |
| | | HexNAc(2)Hex(2)NeuAc(1) | 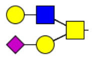 | 1609.6713<br>Full mass detected | 4.23 |
| | | HexNAc(2); HexNAc(2)Hex(2) | 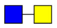<br>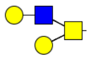 | 1724.7331 | 1.87 |
| | | HexNAc(2)Hex(2)NeuAc(2) | 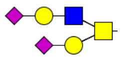 | 1900.7661<br>Full mass detected | 42.49 |
| 245T & 247T | [238]GQQQQGQTVTK[248] | Unoccupied | | | 90.37 |
| | | HexNAc(1) | 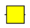 | 1405.6904 | 1.15 |
| | | HexNAc(1)Hex(1) | 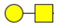 | 1567.7449 | 3.40 |
| | | HexNAc(2) | 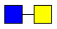 | 1608.7700<br>Full mass detected | 0.03 |
| | | HexNAc(2)Hex(1) | 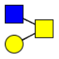 | 1770.8245<br>Full mass detected | 0.06 |

| | | | | | |
|---|---|---|---|---|---|
| | | HexNAc(1)Hex(1)NeuAc(1); | 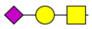 | 1858.8428 | 3.68 |
| | | HexNAc(2)Hex(2) | 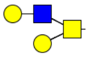 | 1932.8759<br><br>Full mass detected | 0.35 |
| | | HexNAc(2)Hex(1)NeuAc(1); | 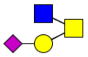 | 2061.9204<br><br>Full mass detected | 0.01 |
| | | HexNAc(1)Hex(1)NeuAc(2) | 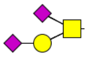 | 2149.9392<br><br>Full mass detected | 0.26 |
| | | HexNAc(2)Hex(2)NeuAc(1) | 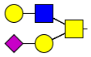 | 2223.9745 | 0.68 |
| | | HexNAc(2)Hex(2)NeuAc(2) | 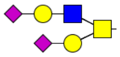 | 2515.0680<br><br>Full mass detected | 0.01 |
| T379 | $^{376}$ADETQALPQR$^{385}$ | Unoccupied | | | 98.96 |
| | | HexNAc(1) | 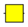 | 1331.645 | 0.01 |
| | | HexNAc(1)Hex(1) | 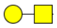 | 1493.698 | 0.10 |
| | | HexNAc(2) | 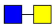 | 1534.724 | 0.01 |
| | | HexNAc(2)Hex(1) | 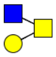 | 1696.779 | 0.03 |
| | | HexNAc(1)Hex(1)NeuAc(1); | 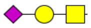 | 1784.794 | 0.09 |
| | | HexNAc(2)Hex(2) | 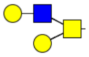 | 1858.829 | 0.18 |
| | | HexNAc(2)Hex(1)NeuAc(1); | 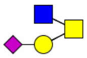 | 1987.871 | 0.01 |
| | | HexNAc(1)Hex(1)NeuAc(2) | 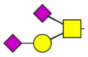 | 2075.888 | 0.09 |

| | | | | | |
|---|---|---|---|---|---|
| | | HexNAc(2)Hex(2)NeuAc(1) | 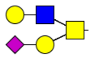 | 2149.926 | 0.31 |
| | | HexNAc(2)Hex(2)NeuAc(2) | 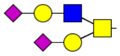 | 2441.022 | 0.22 |
| T391 & T393 | 388KQQTVTLLPA397 | Unoccupied | | 1098.6517 | 0.21 |
| | | HexNAc(1) | 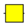 | 1301.7311 | 0.05 |
| | | HexNAc(1)Hex(1) | 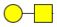 | 1463.783 | 0.04 |
| | | HexNAc(2) | 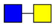 | 1504.8105 | 0.03 |
| | | HexNAc(2)Hex(1) | 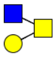 | 1666.863 | 4.52 |
| | | HexNAc(2)Hex(1); Phosphate | 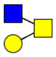 Phos | 1746.8296 | 0.02 |
| | | HexNAc(1)Hex(1)NeuAc(1) | 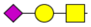 | 1754.879 | 8.19 |
| | | HexNAc(2)Hex(2) | 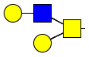 | 1828.916 | 3.73 |
| | | HexNAc(3)Hex(1) | 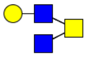 | 1869.944 | 0.65 |
| | | HexNAc(3)Hex(1); Phosphate | 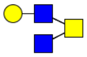 Phos | 1949.909 | 0.21 |
| | | HexNAc(2)Hex(1)NeuAc(1) | 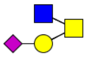 | 1957.958 | 2.00 |
| | | HexNAc(2)Hex(1)NeuAc(1); Phosphate | 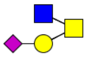 Phos | 2037.925 | 0.18 |

| | | | | | |
|---|---|---|---|---|---|
| | | HexNAc(2)Hex(2)Fuc(1) | 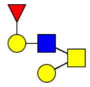 | 1974.975 | 0.21 |
| | | HexNAc(1)Hex(1)NeuAc(2) | 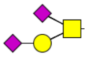 | 2045.974 | 37.75 |
| | | HexNAc(2)Hex(2)NeuAc(1) | 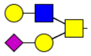 | 2120.013 | 11.26 |
| | | HexNAc(2)Hex(2)NeuAc(1); Phosphate | 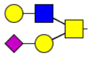 Phos | 2199.9778 | 0.04 |
| | | HexNAc(3)Hex(1)NeuAc(1) | 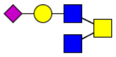 | 2161.037 | 3.40 |
| | | HexNAc(3)Hex(1)NeuAc(1); Phosphate | 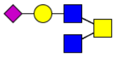 Phos | 2241.0044 | 4.48 |
| | | HexNAc(2)Hex(2)Fuc(1)NeuAc(1) | 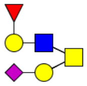 | 2266.068 | 2.24 |
| | | HexNAc(2)Hex(2)NeuAc(2) | 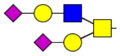 | 2411.106 | 19.63 |
| | | HexNAc(2)Hex(2)NeuAc(2); Phosphate | 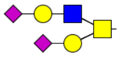 Phos | 2491.0732 | 1.15 |
| S404 | $^{398}$ADLDDFSK$^{405}$ | Unoccupied | | | 53.26 |
| | | HexNAc(2)Hex(1)NeuAc(1) | 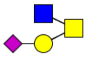 | 1769.7239 | 2.15 |
| | | HexNAc(2)Hex(2)NeuAc(1) | 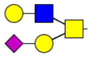 | 1931.7750 | 5.36 |
| | | HexNAc(2)Hex(2)NeuAc(2) | 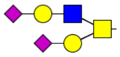 | 2222.8718 | 39.23 |
| 391T/393T/404S | $^{389}$QQTVTLLPAADLDDFSK$^{405}$ | HexNAc(2)Hex(2)NeuAc(2); Phosphate | 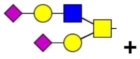 Phos | 3254.3675 | |

| | | | |
|---|---|---|---|
| Phosphorylation | | Or<br>HexNAc(1)Hex(1)NeuAc(1);<br>HexNAc(1)Hex(1)NeuAc(1); Phosphate | 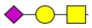<br>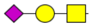<br>+<br>Phos |
